# Genetically inducible coordinators of cytokine signaling pathways for interrogating T cell motility

**DOI:** 10.64898/2026.09.21.753259

**Authors:** Yannick R. Schreiber, Madeleine R. Keller, Xavier S. Bower, Ethan C. Fuller, Jieun Woo, Nikolaos Memmos, David J. Odde, Joshua N. Leonard

## Abstract

Chimeric antigen receptor (CAR) T cell therapy is promising for treating hematologic malignancies, but extending this success to treat solid tumors is challenging. Improving T cell phenotype to address this need is desirable, and one such property is increasing T cell infiltration into tumors. A promising potential approach comprises rewiring cytokine signaling using engineered receptors to change how the T cell responds to environmental cues. However, we currently lack the tools and mechanistic understanding to iterate and improve upon such strategies. Notably, receptors that rewire signaling make it challenging to decouple paracrine effects from those conferred by the signaling inducer. To address this gap, we developed a genetically inducible toolkit of proteins termed Constitutive Activators of Motility-associated Pathways (CAMPs). Building on prior knowledge, CAMPs incorporate domains from IL5R (interleukin-5 receptor) and TNFR (tumor necrosis factor receptor) to place cytokine-associated signaling under direct genetic control, such that expression of a CAMP using a small molecule cue or a condition-responsive promoter induces CAMP signaling. We first identified receptor configurations driving constitutive signaling through targeted pathways. TNFR-based signaling modules drove NF-κB activation across diverse receptor designs, while IL5R-based signaling modules exhibited stringent requirements for membrane-proximity and organization of subunits. We engineered primary human T cells with inducible CAMP circuits, enabling us to probe pathway-specific effects on motility and transcriptomic responses. Pharmacological induction of motility-associated programs downstream of PKC (protein kinase C) was shown to be feasible and dependent on T cell activation state. CAMP induction drove an inflammatory program, particularly when signaling through both NF-κB and STAT5, but none of the conditions tested enhanced 3D motility in our assay. Altogether, our findings are consistent with a model in which migratory behavior may be coupled and regulated by multiple stimuli. These findings and new CAMP tools provide a foundation for interrogating and ultimately harnessing motility for improved T cell therapy performance.

## Introduction

Chimeric antigen receptor (CAR) T cell therapies are a promising approach for treating a range of conditions, from cancer to autoimmune disease.^1, 2^ In oncology, CAR T cells express a receptor directed against a cognate tumor antigen to confer tumor cell-directed cytotoxicity. CAR T cell therapy has proven successful for multiple hematologic malignancies, eliciting response rates of 50-90% in clinical trials.^3–6^ However, CAR T cell treatment of patients with solid tumors remains challenging.^6–9^ Improving T cell phenotype to address this need is an active area of investigation.^10–15^

One of the main challenges to solid tumor CAR T cell therapy is limited tumor penetration. CAR T cells often fail to migrate through the dense tumor stromal matrix of the solid tumor microenvironment (TME),^6^ and this problem is exacerbated by immunosuppressive cell populations and cytokines that limit anti-tumor cytotoxicity^6–9^. Some progress has been achieved by co-opting chemokine signaling to enhance chemotaxis.^16–18^ However, solid tumors frequently exhibit heterogeneous, transient, or spatially confined signaling mediator distributions, rendering gradient-dependent chemokine responses unreliable.^19, 20^ Thus, new approaches are needed.

Several recent studies point to the potential functional benefits of genetic engineering to increase T cell motility. Deletion of RASA2, important in controlling F-actin dynamics, both increased CAR T cell motility and improved tumor control in a mouse model of diffuse midline glioma.^21^ Similarly, expression of constitutively active RhoA (RhoAQ63L), predicted by a biophysical model of T cell motility to enhance T cell migration^22^, resulted in faster T cell migration and better tumor control in a mouse model of pancreatic ductal adenocarcinoma.^23^ Each of these studies exemplifies the strategy of constitutively targeting specific regulators.

Recently, Wirtz et al. reported a technology for modulating T cell motility by coopting signaling pathways, termed Velocity Receptors (VR), which is promising yet raises important questions.^24^ VRs are chimeric receptors engineered with the goal of transducing signals from tumor-associated cytokines (i.e., interleukin-5 (IL-5), tumor necrosis factor alpha (TNFα), interferon gamma (IFNγ), and interleukin-8 (IL-8)) into pathways induced by various pro-inflammatory receptor domains (i.e., IL-5 receptor (IL5R), tumor necrosis factor receptor (TNFR), and interferon gamma receptor (IFNGR)).^24^ These domains were selected based on secreted factors associated with enhanced T cell motility *in vitro*, leveraging the observation that these cytokines are produced by T cells and thus can operate in paracrine and autocrine modalities. The motivating hypothesis is that VRs lock CAR T cells into a high-motility state by binding cytokines the T cells themselves secrete, rendering the phenotype self-propelled rather than dependent on the surrounding secretory milieu. Primary human T cells co-transduced with VRs and CAR were reported to exhibit increased spontaneous migration in 3D collagen matrices, quantified as elevated mean squared displacement (MSD) and an increased fraction of cells in a high motility state. This migratory phenotype is ligand-gated rather than constitutive, so its magnitude is determined by availability of VR-activating cytokines and dictated by local cell density and tumor-intrinsic production. Anti-tumor activity varied across VR constructs and tumor models. Moreover, this initial study did not investigate the intracellular signaling pathways responsible for VR-driven motility. Thus, while this study raised the promising potential for improving CAR T cell function by modulating motility, new tools are needed to parse and modulate the various signaling phenomena that may influence motility in these models and applications.

In this study, we develop and deploy new tools for interrogating T cell motility phenomena, which we term Constitutive Activators of Motility-associated Pathway (CAMPs). Our core goal was to decouple, as much as possible, the effects of inducing signaling pathways of interest from the effects of paracrine and autocrine signaling. We thus pursued a genetically inducible design such that introduction of small molecule cue could induce a defined signaling perturbation at an experimentally determined time and level. We took a synthetic immunology approach to receptor design, systematically identifying structural and signaling features that drive activation of responses through the IL5R, TNFR1, and TNFR2 pathways. Using this validated library of CAMPs, we investigate T cell motility phenomena and related transcriptional responses to these perturbations. Overall, this work presents a useful new set of tools for investigating an important and complex phenomenon of relevance for enabling and potentially enhancing diverse T cell therapies.

## Results

### Development and validation of CAMP proteins

We focused our development of CAMPs on intracellular signaling modules and mediators with well-defined roles in enhancing T cell proliferation, persistence, and effector function. We selected the IL-5 receptor alpha and beta chains (IL5Rα, IL5Rβ) and the TNFR1 and TNFR2 intracellular signaling domains, which engage the STAT5 and NF-κB pathways, respectively. These programs augment adoptive T cell function in the context of cytokine and costimulatory receptor engineering^25–32^, and they were specifically implicated in the VR study to enhance motility^24^. We combined these modules within a single transmembrane receptor-like signaling coordinator protein (note: we avoid the term “receptor” as CAMPs are not designed to sense any ligand), and we investigated two strategies for conferring constitutive signaling (**Fig. 1a**). First, we explored the established approach of fusing intracellular domains to homodimerizing CD4 ectodomains^33,34^. Given that receptor signaling can be influenced by the identity of the transmembrane domain (TMD), we constructed CD4 ectodomain (ECD)-containing CAMP variants with TMDs derived from CD8α (CAMP-1), CD4 (CAMP-2), or CD28 (CAMP-3) (**Fig. 1b, Supplementary Figure 1**).

**Figure 1.**
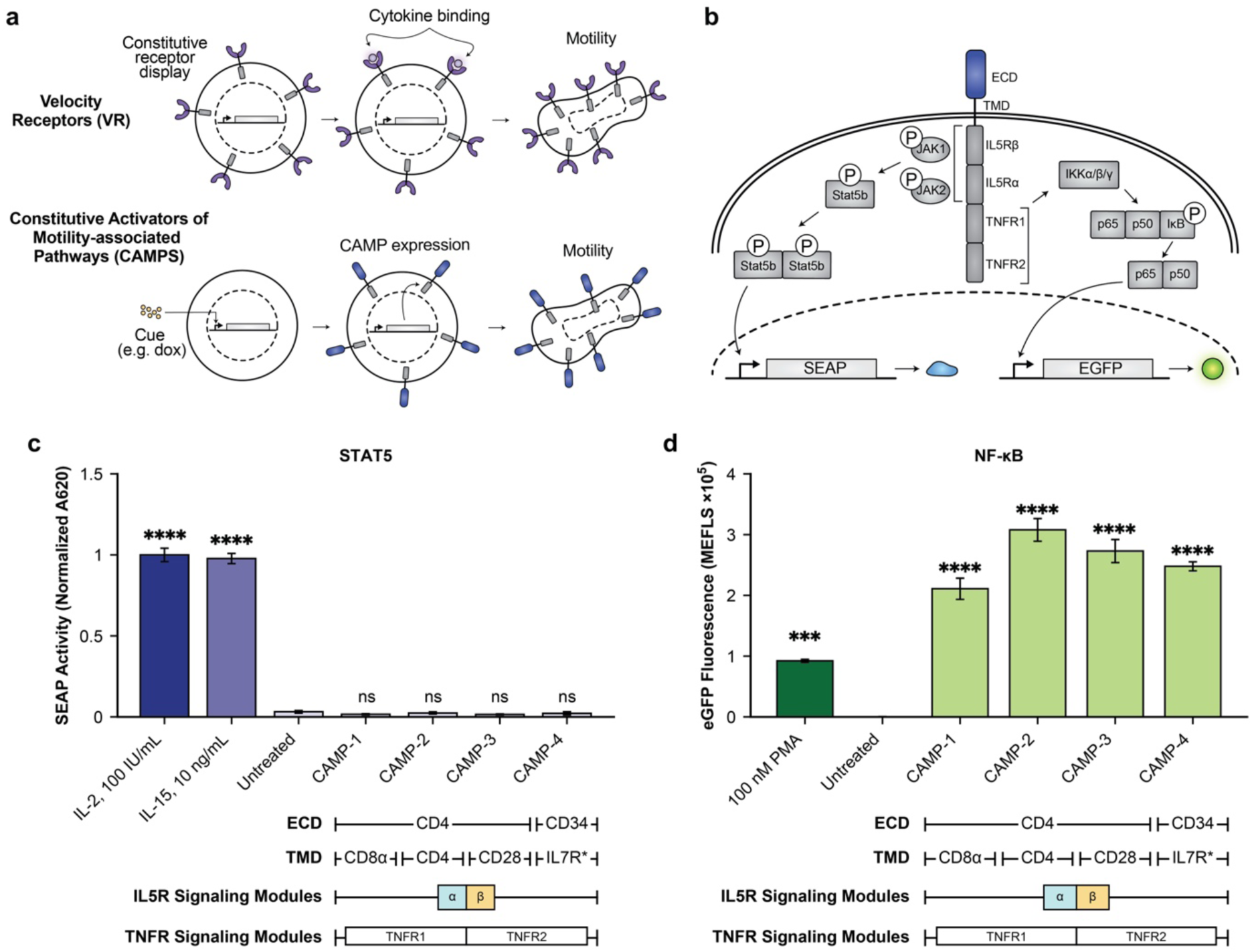
Development of constitutive activators of motility pathways (CAMPs). **(a)** Comparison of CAMPs to a prior technology, Velocity Receptors. Upon the induction of CAMP expression (e.g., via a small molecule regulated system, as shown in this example), CAMPs constitutively drive signaling through motility-associated pathways, whereas Velocity Receptors (VRs) are constitutively displayed and signal in response to cytokine binding. **(b)** CAMP architecture and reporter for monitoring signaling pathways of interest. An ECD–TMD fused to IL5R and TNFR signaling modules routes output to two reporters: JAK1/JAK2-dependent phosphorylation of STAT5b drives a SEAP reporter, and IKKα/β/γ-mediated release of p65/p50 drives an EGFP reporter. **(c,d)** Induction of STAT5 (c) and NF-κB (d) signaling via an initial set of CAMP designs. Cytokine controls (IL-2, IL-15) induce STAT5 activity, and PMA induces NF-κB. Bars show mean ± s.e.m. (n = 3); significance is versus the untreated condition by one-way ANOVA, followed by Dunnett’s multiple comparisons test to evaluate specific comparisons (***p ≤ 0.001; ****p ≤ 0.0001; ns, not significant), under the assumption of equal variance. Additional statistical details are in **Supplementary Note 2**. A repeat of the experiment shown in panels c and d is in **Supplementary** Figure 2. CAMP: Constitutive Activator of Motility Pathways; VR: Velocity Receptor, ECD: extracellular domain, TMD: transmembrane domain, JAK: Janus kinase, STAT5: signal transducer and activator of transcription 5, SEAP: secreted embryonic alkaline phosphatase (normalized A620, absorbance at 620 nm), IKK: IκB kinase, NF-κB: nuclear factor κB, EGFP: enhanced green fluorescent protein, IL5R/TNFR: interleukin-5 / tumor necrosis factor receptor, PMA: phorbol 12-myristate 13-acetate, MEFL: molecules of equivalent fluorescein.

As a second strategy for driving constitutive signaling, we explored a reported approach for engineering a constitutive IL-7 receptor by fusing CD34 ECDs to a mutated IL7R TMD (CAMP-4: CD34-IL7R*+J-α-β-T1-T2)^25^ (**Fig. 1b**). This approach forms disulfide bridges between the mutated CD34-IL7R (C7R) proteins to drive constitutive signaling. For all these CAMP designs, we hypothesized that ECD multimerization would induce proximity between paired IL5Rα/β and TNFR1/2 intracellular signaling domains, respectively, to drive constitutive signaling.

To begin exploring the CAMP concept, we built and tested an initial set of designs. These designs incorporated IL5Rα/B and TNFR1/2 signaling modules in a manner that differs from their native receptor context. We probed signaling through the expected STAT5 and NF-κB pathways, respectively^35–41^, using dedicated reporter constructs (**Fig. 1c**). These designs incorporated some choices made in the VR study^24^, and we also incorporated new designs to address gaps in knowledge as to the determinants of signaling through relevant pathways. To quantify STAT5 activity, we used a commercially available HEK-Blue CD122/132 reporter cell line engineered to detect interleukin-2 and 15 (IL-2 and IL-15) via ectopic expression of IL-2 receptor β (IL2Rβ), IL-2 receptor γ (IL2Rγ), STAT5b, and genomic incorporation of a STAT5-responsive promoter driving expression of secreted alkaline phosphatase (SEAP) for colorimetric assay analysis. NF-κB signaling was assessed using an engineered HEK NF-κB-ELAM-eGFP (HEK-NEE) reporter line, in which NF-κB activation induces eGFP expression measurable by flow cytometry.^42^ HEK-Blue CD122/132 and HEK-NEE reporter cell lines were validated by treatment, respectively, with IL-2 or IL-15, or PMA (phorbol 12-myristate 13-acetate), which activates protein kinase C (PKC) to in turn activate the mitogen-activated protein kinase (MAPK), extracellular signal-regulated kinase (ERK), and phosphoinositide 3-kinase (PI3K), engaging cascades that promote the activation of NF-κB^43–46^ (**Fig. 1c,d**). While our first set of CAMP designs induced signaling through NF-κB (**Fig. 1d**), no STAT5 signaling was detected (**Fig. 1c**). These data suggest that CAMP 1-4 can form active signaling complexes (mediated by TNFR1 and or TNFR2 domains), but the IL5R domains cannot induce signaling. We hypothesized that such disparity could be explained by incompatibility with the geometric requirements for inducing IL5R signaling, including membrane proximity, steric access to signaling domains, and subunit ordering, motivating the next design iteration.

To attempt to identify CAMP designs that confer IL5R-mediated signaling, we next constructed a set of proteins sampling key feature choices (**Fig. 2a, Supplementary Figure 3**). These constructs again sampled a range of ECD, TMD, and juxtamembrane domain (JMD) choices, and we varied the order of IL5Rα and IL5Rβ intracellular domains and introduced flexible (G4S)_3_ linkers to separate these domains and decrease steric hindrance. We also tested both wild-type IL5Rβ domains and a constitutively active IL5Rβ domain variant containing a mutation (R461C) in the first juxtamembrane residue that introduces a disulfide bridge that drives receptor association and induces constitutive signaling^47^. All constructs retained downstream TNFR1 and TNFR2 domains, separated from the IL5R domains by a (G4S)_3_ linker.

**Figure 2.**
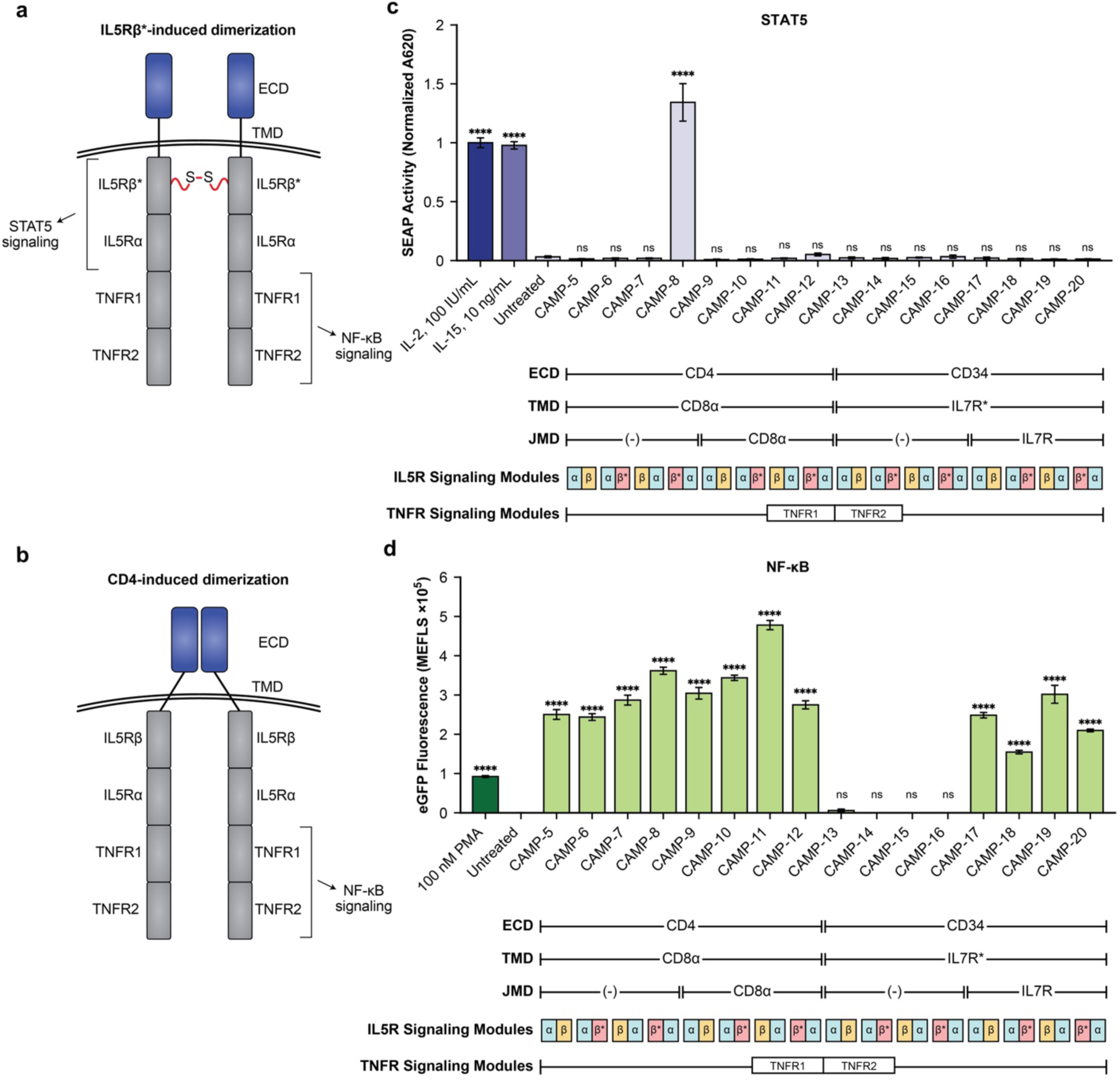
Refined CAMP designs signal through both STAT5 and NF-κB. (a,b) Summary of new CAMP designs tested to interrogate requirements for STAT5 signaling, along with proposed mechanisms. The disulfide bond (S–S) shown is hypothesized to be conferred by the mutation (R461C), which is included in constructs containing the domain designated IL5Rβ*. **(c,d)** Induction of STAT5 (c) and NF-κB (d) signaling via new CAMP designs. Bars show mean ± s.e.m. (n = 3); significance is versus the untreated condition by one-way ANOVA, followed by Dunnett’s multiple comparisons test to evaluate specific comparisons (****p ≤ 0.0001; ns, not significant), under the assumption of equal variance. Additional statistical details are in **Supplementary Note 2**. A repeat of the experiment shown in panels c and d is in **Supplementary** Figure 4. ECD: extracellular domain; TMD: transmembrane domain, JMD: juxtamembrane domain, IL5R: interleukin-5 receptor, TNFR: tumor necrosis factor receptor, SEAP: secreted embryonic alkaline phosphatase (normalized A620), NF-κB: nuclear factor κB, PMA: phorbol 12-myristate 13-acetate, MEFL: molecules of equivalent fluorescein.

Functional analysis of this next set of CAMP designs identified strict requirements for inducing STAT5 signaling and some constraints on NF-κB signaling, yielding viable CAMP constructs. Across the sixteen rationally designed architectures tested, STAT5 activation was observed in only a single construct, CAMP-8 (**Fig. 2c**), while many designs signaled through NF-κB (**Fig. 2d**). CAMP-8 positioned the mutant IL5Rβ R461C signaling domain immediately proximal to the membrane, followed by a flexible (G4S)_3_ linker and the IL5Rα signaling domain. All alternative configurations, including reversed subunit orders containing the same R461C mutation, failed to elicit STAT5 signaling. We also confirmed that CAMP-8 effectively traffics to the cell surface, supporting its proposed mechanism for signal induction (**Supplementary Figure 5**). Induction of NF-κB signaling was observed across many variations, but constructs incorporating the CD34 ECD fused to the mutant IL7R TMD required inclusion of the IL7R JMD to enable NF-κB activation; removal of the JMD abolished signaling, indicating that membrane-proximal motifs remain critical for TNFR signaling in certain contexts. These results demonstrate that IL5R-mediated STAT5 signaling is highly sensitive to receptor geometry, requiring precise membrane-proximal positioning and subunit ordering, whereas TNFR-mediated NF-κB signaling is comparatively permissive across a broad range of contexts. Importantly, this study also yielded vetted CAMP designs that are competent for inducing signaling through the pathways of interest.

### Extracellular CAMP domains modulate CAMP signaling but are not required

Having identified signaling-competent CAMP designs, we next probed the role of the ECD. Our findings thus cannot determine whether the ECD is required or modulates signaling intensity or selectivity through the pathways of interest, and so we built a series of new designs to investigate these phenomena **(Supplementary Figure 6**). To address whether multimerizing ECDs are required for signaling, we generated variants of CAMP-8 in which the multimerizing CD4 ECD was replaced with ECDs that exist as monomers from either CD34 (CAMP-21) or IL10RB (CAMP-22), or the ECD was removed entirely (CAMP-23). Interestingly, all ECD variants of CAMP-8 retained the ability to induce signaling via STAT5 (**Fig. 3a**). The magnitude of STAT5 signaling varied across constructs, and STAT5 signaling was preserved even in the absence of an ECD. All constructs also retained robust NF-κB signaling activity (**Fig. 3b**). These findings indicate that ECD-mediated clustering is not strictly required for constitutive CAMP signaling given functional intracellular design choices. ECDs do appear to modulate signaling magnitude, with CD34 ECD-containing CAMP-21 conferring reduced signaling through both pathways compared to other designs (possibly indicating a stability or trafficking defect). The ability of a minimal receptor lacking an ECD to signal constitutively highlights the intrinsic sufficiency of intracellular domain organization in driving downstream STAT5 and NF-κB signaling.

**Figure 3.**
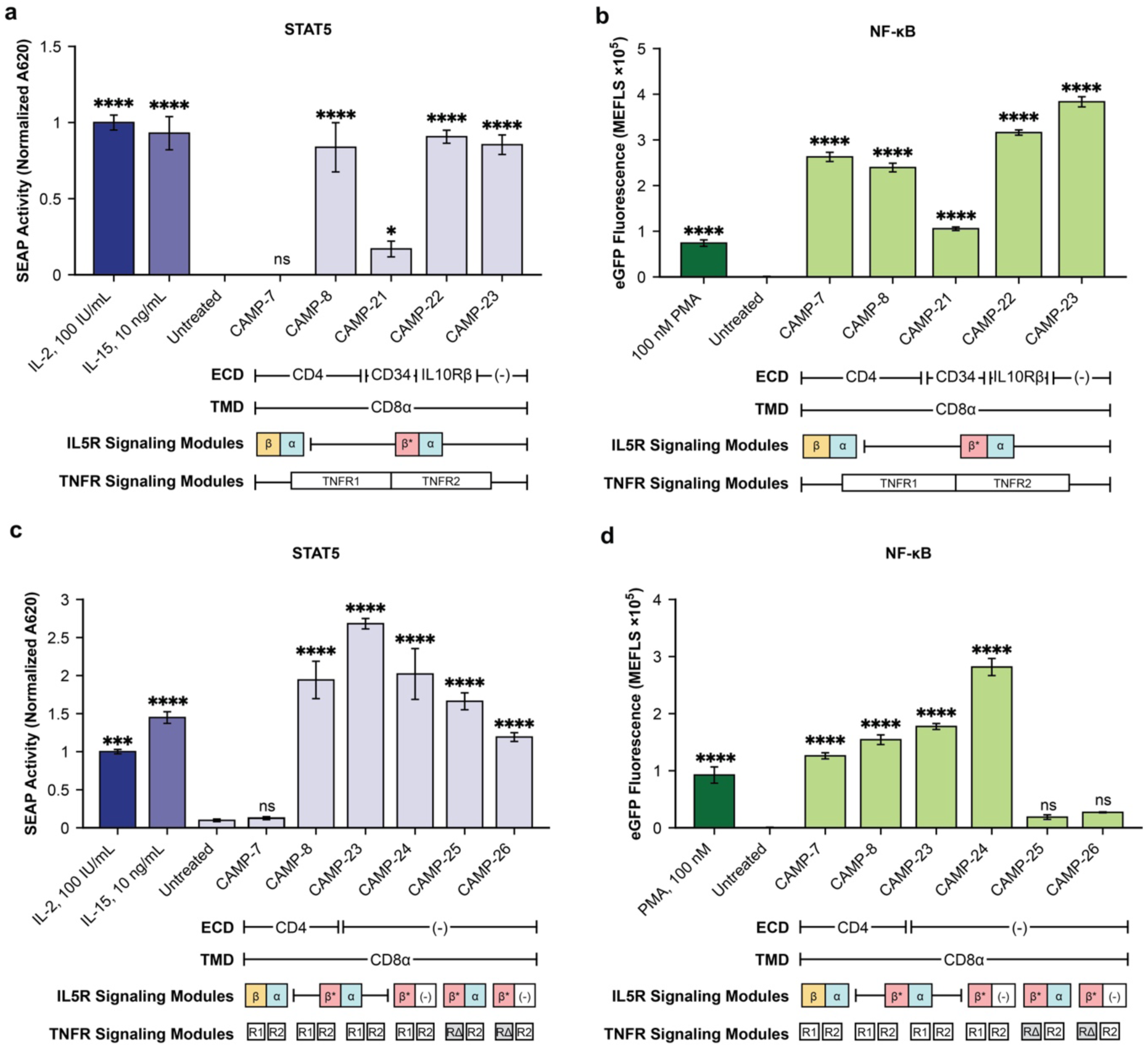
Extracellular domain and TNFR death domain choices shape CAMP signaling. (a,b) Induction of STAT5 (a) and NF-κB (b) signaling via CAMP designs probing the role of the ECD. **(c,d)** Induction of STAT5 (c) and NF-κB (d) signaling via CAMP designs probing the role of intracellular domains. Bars show mean ± s.e.m. (n = 3); significance is versus the untreated condition by one-way ANOVA, followed by Dunnett’s multiple comparisons test to evaluate specific comparisons (*p ≤ 0.05; ***p ≤ 0.001; ****p ≤ 0.0001; ns, not significant), under assumption of equal variance. Additional statistical details are in **Supplementary Note 2**. A repeat of the experiments shown in panels a–d is in **Supplementary** Figure 7. ECD: extracellular domain; TMD: transmembrane domain. IL5R: interleukin-5 receptor, IL10Rβ: interleukin-10 receptor β, TNFR: tumor necrosis factor receptor, TNFRΔDD1: TNFR1 death domain deletion variant (RΔ), SEAP: secreted embryonic alkaline phosphatase (normalized A620), NF-κB: nuclear factor κB, PMA: phorbol 12-myristate 13-acetate, MEFL: molecules of equivalent fluorescein.

### IL5Ra is dispensable for constitutive STAT5 signaling whereas TNFR1 death domain is required for NF-κB signaling

After identifying which extracellular components are necessary for CAMP-mediated induction of STAT5 and NF-κB signaling, we next sought to determine which elements of each signaling module are necessary or sufficient to induce signaling. We found that the IL5Rβ domain must contain the R461C mutation and be immediately proximal to the membrane to induce STAT5 signaling (**Fig. 2a**), but it remains unknown whether the downstream IL5Rα domain is necessary for activating this pathway. Additionally, the TNFR1 signaling module contains a native TNFR death domain (TNFRDD), which is responsible for recruitment of the Tumor necrosis factor Receptor Type 1-Associated DEATH Domain (TRADD) protein, activating apoptotic, necroptotic, and NF-κB pathways^48^. Apoptosis and necroptosis are undesirable pathways to activate in the context of cell-based therapeutics and could complicate motility studies. Therefore, we investigated whether this TRADD-binding death domain could be deleted from the TNFR1 module without ablating the ability to induce (potentially useful) NF-κB signaling.

To test these mechanistic questions, we constructed three new CAMP proteins that are variants on CAMP-23. One construct omitted the IL5R*a* domain (CAMP-24), a second omitted the TNFRDD (CAMP-25), and a third construct contained deletions of both the IL5R*a* and TNFRDD modules (CAMP-26) (**Supplementary Figure 6**). CAMPs omitting the IL5R*a* signaling module induced STAT5 signaling, indicating it is dispensable (**Fig. 3c**). CAMPs omitting the TNFRDD exhibited substantially reduced NF-κB signaling, indicating a critical role for this domain (**Fig. 3d**). Altogether these findings pinpoint the components required for robust CAMP-mediated induction of STAT5 and NF-κB signaling arms, positioning us to deploy CAMPs for functional studies.

### Developing an assay for quantifying primary human T cell motility phenomena

We next sought to develop an assay for evaluating T cell motility. As several approaches with differing methodologies have been reported, we sought to validate a method by evaluating primary T cell motility phenomena prior to moving toward CAMP-mediated perturbations. We adopted a standard 3D collagen matrix model, which is commonly used for cell migration studies (**Fig. 4a**).^49–52^ Using time-resolved phase contrast microscopy, we observed movement of T cells within this 3D collagen matrix within a single collapsed XY plane. To analyze these data, we developed a computational pipeline using the TrackMate plugin^53, 54^ within FIJI/ImageJ^55^ software (**Supplementary Notes 3,4**). From TrackMate-resolved cell movement tracks, we quantified single-cell mean-squared displacement (MSD) to quantify migration. To establish baseline conditions for our assay, we evaluated motility across a range of cell plating densities, which has been reported to increase cellular motility,^24^ and we explored several assay variations involving activation of various signaling pathways (**Fig. 4b, Supplementary Figure 8a)**. Cells were either activated with anti-CD3/CD2/CD28 ImmunoCult reagent 24 h prior to encapsulation in the gels (pre-activated) or immediately after encapsulation within gels (in-assay activated); in all cases, 48 h after gel encapsulation, cells were imaged for one hour. Since PMA treatment has been reported to increase primary T cell motility via PKC-driven signaling,^56^ we also treated cells with 50 ng/mL of PMA immediately prior to imaging. Since there is no one agreed-upon way to quantify motility of a population, we analyzed our data in several ways (described in detail in **Methods, Supplementary Note 3**). While some prior work identifies T cells as falling into low-and high-motility subpopulations using a fixed threshold (e.g., a MSD of 25 μm^2^ for the VR study^24^), we found this method to poorly describe variation in our data. Instead, we quantified (i) the frequency of cells falling into low-and high-motility subpopulations by fitting nominally bimodal MSD distributions using a Gaussian mixture model, as previously described^57, 58^; (ii) the central tendency (geometric mean) MSD for each identified subpopulation; and (iii) the overall arithmetic mean motility of each condition, which we defined relative to a reference condition (which is given a value of unity) within each experiment (i.e., relative motility). Using these metrics, we evaluated the motility phenomena of interest. We also characterized two technical confounds that could shape how these metrics should be interpreted: track-length filtering, which biases surviving cells toward higher MSD (**Supplementary Figure 9**), and variation in per-well cell recovery, which moves state occupancy but not state position (**Supplementary Figure 10**). Track length distributions and per-well recovery variation were not found to create bias of a scale sufficient to alter our conclusions.

**Figure 4.**
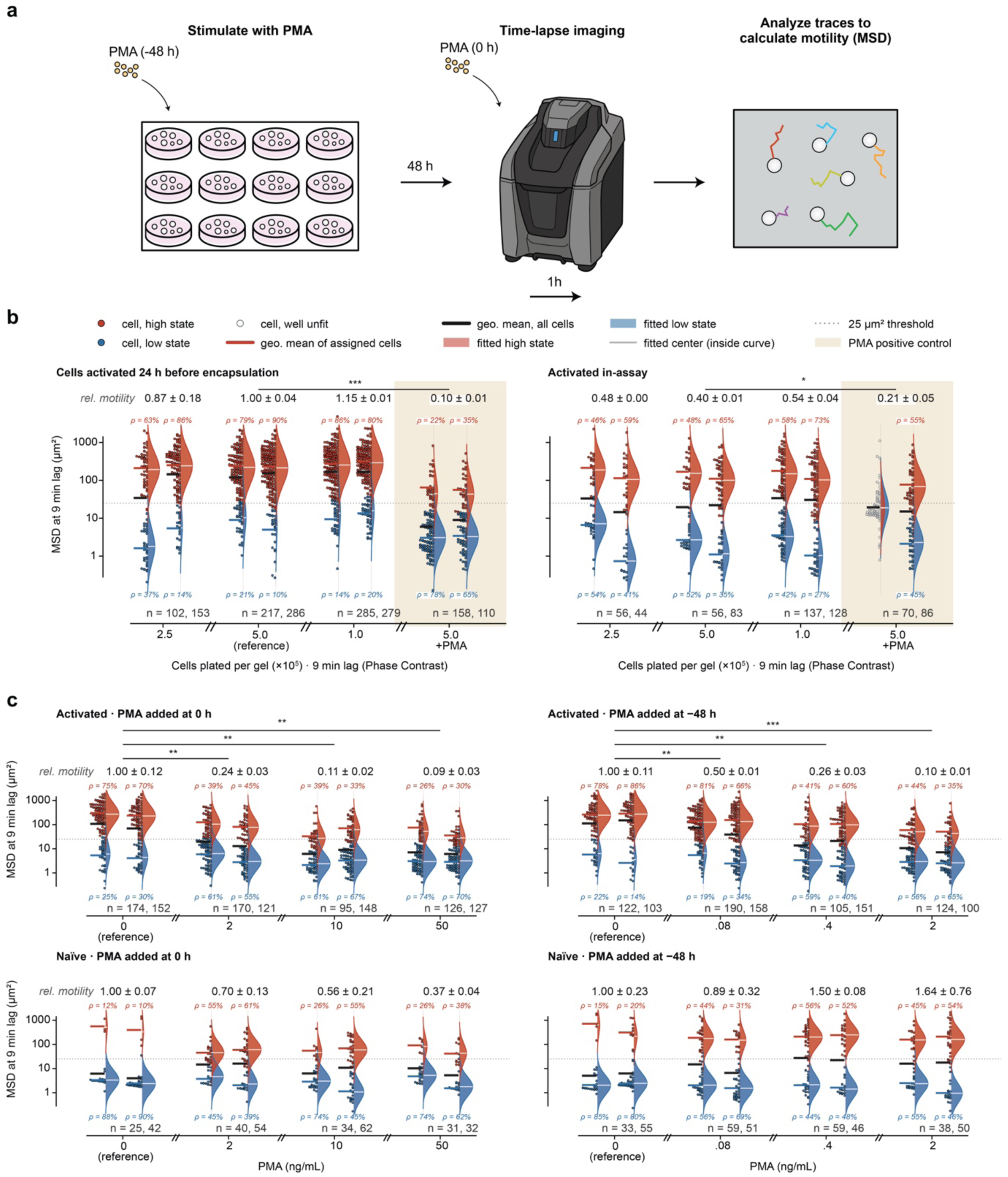
Development of an assay for studying primary human T cell motility. **(a)** Experimental schematic exemplifying a general T cell motility analysis workflow. **(b)** Interactions between plating density and timing of activation on motility. Unmodified primary human T cells were activated with ImmunoCult either 24 h before encapsulation or at the time of encapsulation, and some samples were stimulated with PMA (50 ng/mL) as indicated. **(c)** Interactions between activation and PMA treatment regimens on T cell motility. Naïve cells were never activated with ImmunoCult, while activated cells were treated 24 h before encapsulation. In (b) and (c), each condition is shown as a per-cell swarm colored by fitted motility state (see legend) beside its fitted two-state mixture, with wells analyzed individually. Relative motility for each condition is calculated as the mean of the per-well arithmetic mean MSDs (excluding cells that yielded no estimate) relative to the equivalent value calculated for the relevant reference condition ± SEM across wells (reference conditions are indicated, with a single condition used for panel (b) and individual reference conditions defined for each subplot in (c)). n = cells per well identified and used in the MSD calculation. Significance: pooled-error model on log10 per-well arithmetic mean MSD, with one within-condition variance estimated from all wells of the panel (8/4 residual df in panels b/c); each PMA condition is compared against its prespecified reference and p values are corrected within each panel by Benjamini–Hochberg (* q<0.05, ** q<0.01, *** q<0.001, **** q<0.0001; ns, not significant). n = 2 wells per condition. Additional statistical details are in **Supplementary Note 2**. MSD: mean squared displacement; PMA: phorbol 12-myristate 13-acetate.

Our results generally support the use of this methodology to analyze motility phenomena, and additionally suggest a few interesting features of T cell motility. First, we observed at most a modest effect of plating density: relative motility varied by no more than 1.3-fold across a four-fold range of plating densities (2.5 × 10^5^ to 1 × 10^6^ cells per gel) in gels containing both pre-activated and in-assay activated T cells (**Figure 4b, Supplementary Figure 8a**). Any such effect is therefore much smaller than the effects of activation timing and PMA treatment reported herein. For both in-assay activated and pre-activated cells, treatment with PMA led to lower relative motility (**Figure 4b**). Because relative motility reduces each condition to a ratio against its respective control, we also examined motility by comparing the two fitted states (**Supplementary Figure 8b-c**). We noted a few distinct qualitative trends through which PMA appears to act. In activated cells, PMA reduced motility of the high motility state, while reducing the share of cells occupying that state. In naïve cells, PMA instead raised the frequency of cells in the high motility state. We also noted that the naïve untreated samples contained too few cells in the high motility state to draw conclusions about the properties of this subpopulation. These observations guide our subsequent analyses.

One possible reason for this result is that these conditions differ somewhat from prior studies in which treatment of naïve (rather than ImmunoCult-activated) T cells with PMA immediately prior to imaging drove induced motility^56^. Thus, we also compared the responses of activated and naïve T cells to PMA across various doses and timings. In the “acute” PMA regimen, cells were dosed immediately prior to time course imaging, and in the “chronic” PMA regimen, cells were dosed with lower concentrations of PMA (to reduce toxicity) 48 h prior to time course imaging. Pre-activated T-cells were more motile than naïve cells, which was most noticeable in the greater frequency of high-motility cells (**Fig. 4c**). The addition of PMA led to reduced relative motility in activated T cells under both acute and chronic PMA regimens, whereas PMA treatment generally increased the frequency of high-motility cells in the naïve cell samples, consistent with prior reports^56^ (**Fig. 4c**). Thus, these observations reconcile apparent discrepancies in PKC-induced T cell motility programs across T cell activation states and support the use of our pipeline to investigate CAMP-mediated motility phenomena.

### Developing a method for engineering inducible CAMP expression in primary T cells

We sought a system for engineering primary human T cells to investigate the impact of CAMP-mediated signaling. We selected a small molecule-inducible gene expression system to provide external control over the timing and degree of CAMP expression. Constitutive expression of pro-migratory signaling receptors has the potential to confound phenotypic analysis through chronic activation or adaptation during *ex vivo* cell culture. An inducible system avoids this problem and, moreover, enables characterization of the same engineered cell sample with or without CAMP expression. The total cargo size for a CAMP (e.g., 2.9 kB for CAMP-24) plus inducible machinery (e.g., pTRE-3G which is doxycycline (dox)-inducible) exceeds the nominal capacity of lentiviral payloads, so we selected a transposon vector. PiggyBac was selected due to its large cargo capacity, high degree of stable genomic integration, and compatibility with GMP manufacturing and clinical use.

Given these goals, we designed and validated a transposon-based T cell engineering workflow. Our transposon (∼17.5 kB in length) includes three separate transcriptional units: (i) a Tet response element (TRE) promoter driving CAMP expression, (ii) a constitutive CAG promoter driving expression of a 4αβ chimeric cytokine receptor (selection marker) linked via a T2A self-cleaving peptide to a mNeonGreen (marker of successful engineering), and (iii) a human EF1α promoter driving expression of the Tet-On 3G transactivation protein (**Fig. 5a**). The 4αβ receptor enables enrichment of engineered cells.^59^ By fusing the IL-4 receptor ectodomain to the βc signaling domain shared by the IL-2 and IL-15 receptors, 4αβ delivers an IL-2/IL-15 signal upon binding to IL-4, enabling selective *ex vivo* expansion and enrichment of engineered T cells by applying IL-4 and withdrawing IL-2 (starving unmodified cells of this signal), without affecting cytotoxicity.^59^ Since 4αβ-mediated selection is most effective during the proliferative phase of primary T cell expansion, primary T cells were electroporated with the PiggyBac transposon and transposase, and then IL-4 was added concurrently with CD3/CD28-stimulation to initiate selective expansion of engineered T cells from the outset of culture (**Fig. 5b**). This workflow was informed by prior work demonstrating improved transgene integration when transposon electroporation and integration precedes activation-driven proliferation in T cells.^60^ Cells were expanded under IL-4 selection for 10-14 days, after which integration efficiency was assessed using flow cytometry (**Fig. 5c**). We consistently achieved approximately 35% engineered cells, which is excellent given the large construct size. To our knowledge, this represents one of the largest inducible synthetic transgene systems for engineering primary human T cells using a translatable manufacturing approach.^60–62^ Moreover, this validated engineering method provides a tool for investigating CAMP-induced phenomena.

**Figure 5.**
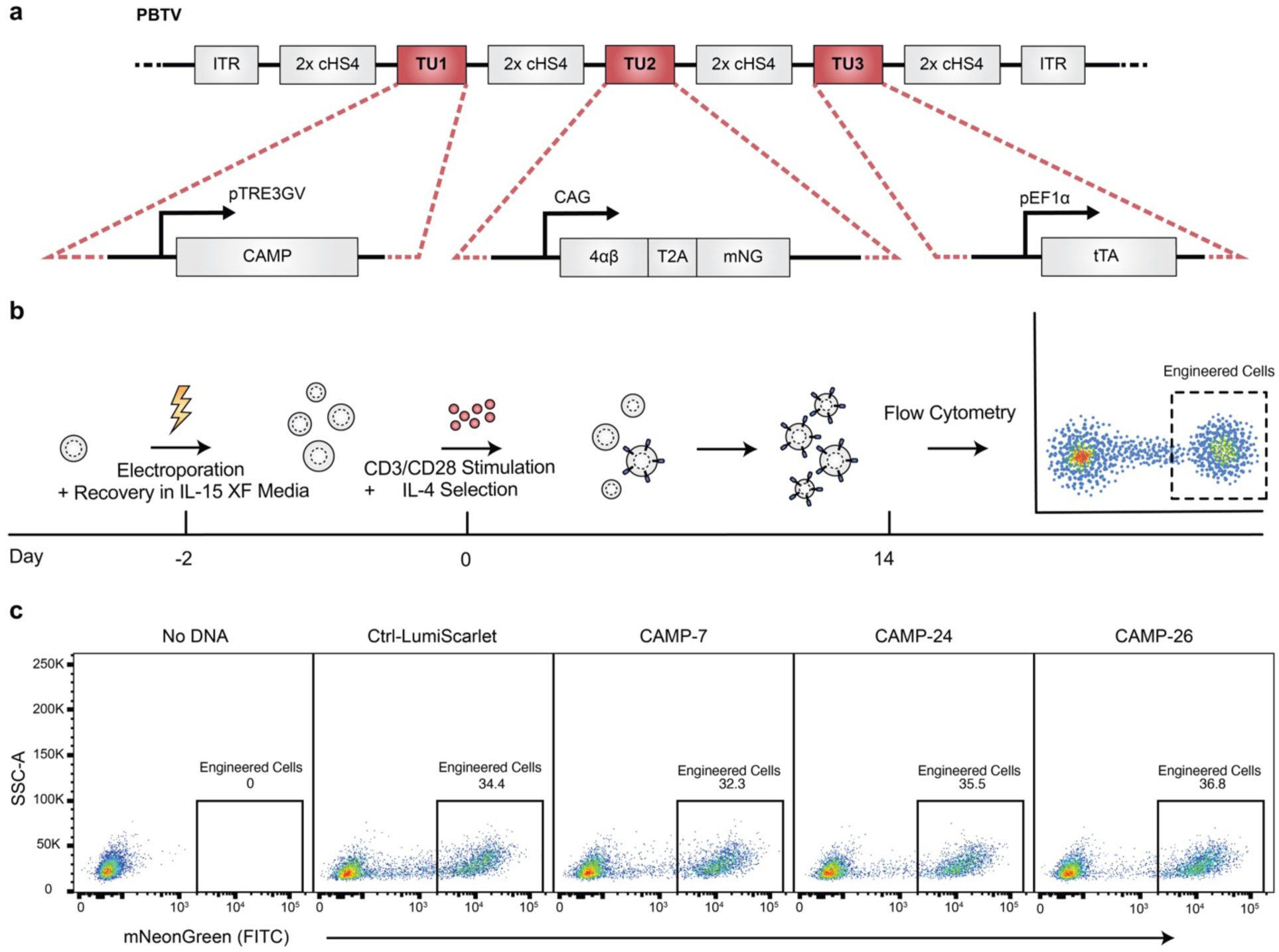
Vector design and generation of engineered primary T cells. **(a)** PiggyBac Transposon Vector architecture. The PBTV construct carries three insulated transcription units between ITRs, with details shown in expanded views in lower row, separated by 2x cHS4 insulators. **(b)** Engineering timeline. Cells are electroporated and recovered in IL-15 XF medium (day − 2), stimulated via CD3/CD28 and IL-4-selected (day 0), and analyzed by flow cytometry (day 14). **(c)** Quantification of T cell engineering outcomes (percent engineered) for an exemplary run. PBTV: piggyBac transposon vector; ITR: inverted terminal repeat. TU: transcription unit; cHS4: chicken β-globin hypersensitive site-4 insulator; tTA: tetracycline-controlled transactivator; T2A: self-cleaving 2A peptide; mNG: mNeonGreen; SSC-A: side-scatter area; FITC: fluorescein isothiocyanate channel.

### CAMP-induced T cell phenotypes and transcriptional states

Having validated the necessary CAMP technology and methods, we next investigated how induction of STAT5 and/or NF-κB signaling impacts T cell state as defined by two outputs: (i) motility and (ii) transcriptome changes (**Fig. 6a**). We selected a subset of CAMPs that signal through NF-κB only (CAMP-7), STAT5 and NF-κB (CAMP-24), and STAT5 only (CAMP-26) and engineered inducible T cell lines. We also generated a control line which is similarly engineered and selected but in which dox induces expression of LumiScarlet. Engineered cells were embedded in 2 mg/mL collagen gels, tracked by single-cell imaging, and analyzed as in **Figure 4** (**Fig. 6b**). The same cells were then recovered from the same gels for bulk RNA-sequencing and transcriptomic analysis (**Fig. 6c, Supplementary Figures 11-14**), such that motility and transcriptional readouts were obtained from the same cells under matched conditions. This process was repeated for two independent engineering runs using donor-matched T cells.

**Figure 6.**
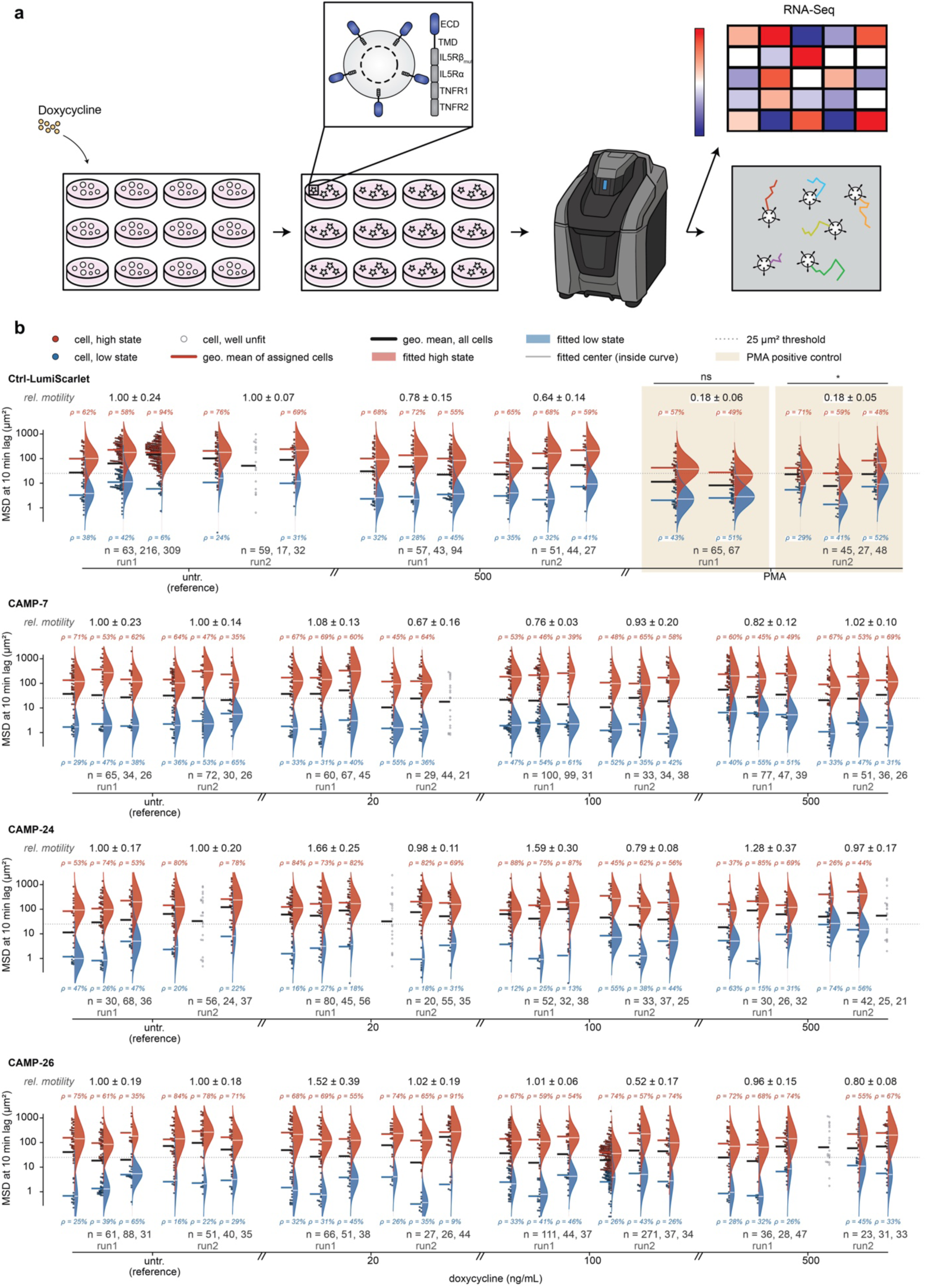
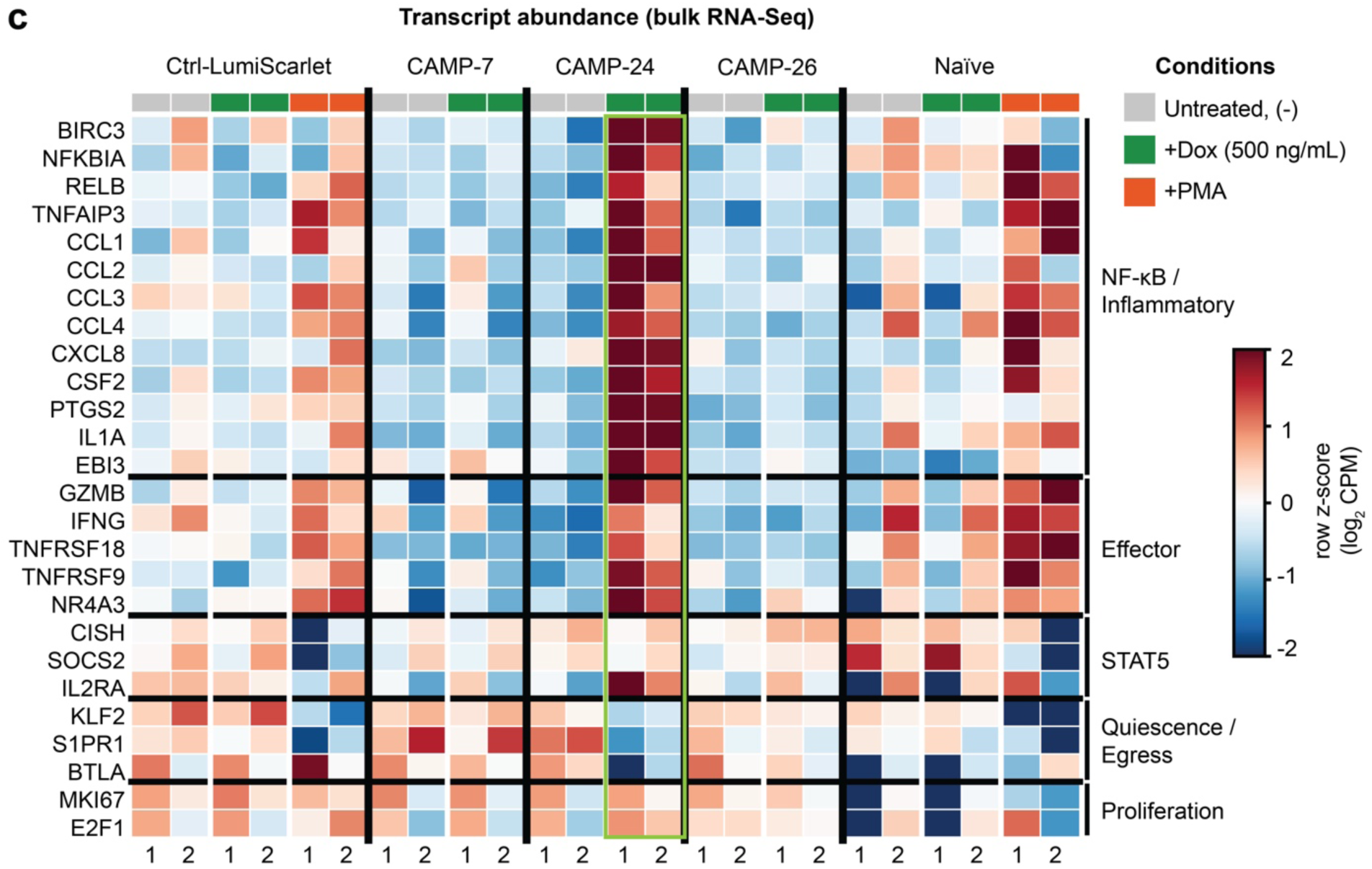
Modulation of T cell state by inducing CAMP expression. **(a)** Overall workflow for investigating effects of CAMP expression on T cell motility and transcriptomic state. **(b)** T cell motility analysis. Motility, relative motility, and high-and low-motility populations were quantified as in Figure 4. Cells were treated with dox (as indicated) and/or PMA (0.4 ng/mL) for 60 h prior to imaging (1h) followed by harvest of cells for RNA-seq. **(c)** Transcriptomic analysis of T cells after CAMP induction and/or PMA activation (see Fig. 6 **continued**). This analysis includes all samples characterized for motility in panel (b) (run number at bottom of plot), plus an additional control set conducted in parallel on naïve (non-engineered) T cells and treated as indicated, matching conditions in (b). Differential gene expression was calculated relative to the untreated condition within each run. Genes were grouped by program (bin labels on right side of heat map). Dox-induced expression of CAMP-24 (signaling via both STAT5 and NF-κB) confers a reproducible signature (highlighted in light green box). Additional statistical details are in **Supplementary Note 2**. Dox: doxycycline; GMM: Gaussian mixture model; MSD: mean squared displacement; RNA-seq: RNA sequencing; CPM: counts per million; PMA: phorbol 12-myristate 13-acetate.

Our combined analysis revealed new insights into the effects and interactions of STAT5 and NF-κB-induced signaling in T cells. In general, all of our engineered lines (including Ctrl-LumiScarlet) exhibited motility distributions that exhibit the higher motility observed when non-engineered T cells were activated (**Fig. 4**). Induction of CAMP expression did not substantially reduce or enhance motility, using our setup and metrics, whereas PMA treatment of Ctrl-LumiScarlet cells reduced motility (**Fig. 6b**), as seen with activated non-engineered cells (**Fig. 4**). Interestingly, only induction of CAMP-24 (which activates both STAT5 and NF-κB) produced a significant, reproducible transcriptional response (1,334 genes at FDR < 0.05; 274 at FDR <0.05 and |log2FC| > 1; **Fig. 6c, Supplementary Figure 12, Supplementary Data 1**), yielding the highest concordance between runs (**Supplementary Figure 11**). CAMP-24 induced transcription along both arms of NF-κB signaling: the canonical feedback regulators *NFKBIA*, *TNFAIP3*/A20, *BCL3*, and *NFKBIE,* and the non-canonical mediators *RELB* and *NFKB2* were all significantly upregulated, along with NF-κB target *BIRC3*, consistent with TNFR1/TNFR2 signaling modules within the receptor. This was accompanied by expression of genes associated with the inflammatory secretome, with significant induction of cytokine and chemokine genes *CSF2* (GM-CSF), *CCL2*, *CCL1*, *CCL3*, *CCL4*, *CXCL8*, *IL1A*, *IL13*, *IL22*; the prostaglandin synthase *PTGS2* (COX-2); and the IL-27/IL-35 subunit *EBI3*. Induction also produced a significant effector and activation signature: the cytotoxic protease granzyme B (*GZMB*), *IFNG*, the costimulatory receptors *TNFRSF18*/GITR and *TNFRSF9*/4-1BB, the AP-1 factor *JUN*, and *EGR2* and *NR4A3*, accompanied by significant down-regulation of the quiescence and lymph-node-egress genes *KLF2*, *S1PR1*, *IL7R*, and *SELL*/CD62L, together suggesting a resting-to-effector transition. Notably, the response did not engage two other transcriptional programs. The canonical STAT5 target genes *CISH* and *SOCS2* remained unchanged, and there was no evidence of cell-cycle activation (*MKI67*, the *MCM* helicases, *CDK1*, *TOP2A* all showed no significant change; only *MYC* and *E2F1* rose modestly). Altogether, CAMP-enabled disentanglement of these interacting pathways revealed an important insight; at least within the context of these inducible, sustained signaling systems, signaling through both STAT5 and NF-κB yields a stronger response than signaling through either pathway alone, and this combined response resembles NF-κB-mediated patterns more so than canonical STAT5-driven responses.

## Discussion

This study developed and employed CAMPs—a new set of genetic tools enabling research toward the goal of enhancing T cell motility to improve therapeutic outcomes. CAMPs provide new technology for activating specific motility-associated pathways of interest in a manner that is largely decoupled from complicating influences, especially compared to previously available technologies. These findings shed new light on T cell responses associated with key pathways of interest—STAT5 and NF-κB—and provide a foundation for deploying CAMPs in diverse contexts for probing and illuminating the complex phenomenon of T cell motility.

A key accomplishment of this study was addressing gaps in our understanding of how to engineer membrane proteins to drive signaling through motility-associated pathways. We focused on STAT5 and NF-κB pathways, which are implicated in enhanced adoptive T cell function and have been identified as important in receptor engineering studies.^25–32^ While the VR study motivating much of this work also implicated these pathways in enhanced motility, it did not identify VR features necessary to induce such signaling.^24^ Our systematic CAMP engineering helped to close these gaps.

Our data are consistent with a model in which TNFR signaling is comparatively tolerant to variations in receptor architecture and can be engaged across multiple clustered receptor configurations. However, in constructs with CD34 ECDs fused to IL7R TMDs, it appears that the presence of an intracellular JMD region is necessary for TNFR signaling. TNFR signaling requires the aggregation of receptor trimers to recruit signaling complexes in hexagonal networks, forming intracellular membrane-proximal lattices.^63^ Loss of the JMD on IL7R constructs may prevent recruitment of adaptor proteins and abrogate signaling. Alternatively, the IL7R TMD may require its native JMD for proper helix exit and orientation. The mutated IL7R used in this study contains an unpaired cysteine residue within the JMD, which is a common mechanism for induction of constitutive IL7R signaling often found in hematologic malignancies.^25, 64^ Thus, removing this JMD removes unpaired cysteines and precludes disulfide bond formation between IL7R chains, which could ablate constitutive signaling.

In contrast, IL5R signaling is geometrically constrained. The native IL5Rβ receptor, also known as colony stimulating factor 2 receptor beta, complexes with ligand-specific alpha chains to form dodecameric complexes that cause associated JAK2 kinases to trans-phosphorylate and induce signaling.^47, 65^ Only CAMP-8 and its derivatives, which contain a mutant IL5Rβ immediately proximal to the membrane, enabled constitutive signaling. Like the mutant IL7R domain, R461C IL5Rβ includes an unpaired cysteine that is hypothesized to drive formation of IL5Rβ complexes to induce signaling. Given the requirements for IL5Rβ domains to form dodecameric complexes and TNFR domains to form hexagonal lattices, it is possible that CAMP-8 and its derivatives benefit from reciprocally supporting intracellular IL5Rβ and TNFR inter-chain interactions that drive the robust STAT5 and NF-κB signaling observed.

T cell motility is modulated—both up and down—by a variety of factors, and this study provided new insight into how the timing and nature of such signals in the pathways of interest impact motility. In general, our findings argue against a simple model of “motility enhancing factors” and concord with current models in which T cell migration is governed by a push and pull between stop and go signals.^66^ For example, pharmacological activation of PKC using PMA modulated motility of primary T cells in a manner that depends on activation state; PMA reduced motility of activated T cells, while it increased motility of naïve T cells. Induction of CAMPs driving STAT5 and/or NF-κB did not substantially reduce or enhance motility of engineered cells, whereas PMA treatment reduced motility. We do not attribute this difference to the magnitude of signaling through NF-κB, as CAMPs induced stronger activation of this pathway than did PMA (in HEK-NEE cells). This suggests that other downstream mediators of PKC signaling may drive an alternative, stronger “Stop” signal than does NF-κB signaling alone. We noted that our engineered T cells exhibited a more motile phenotype (prior to any stimulation) than naïve T cells, and it is possible that modulating the engineering method to make the basal state less motile could change the motility response to CAMP induction. Finally, CAMP induction drove a distinct transcriptional program when STAT5 and NF-κB were co-activated, compared to activating each pathway alone. That response most closely resembles a canonical NF-κB response. A possible contributing factor is that CAMP-24 (which signals through both pathways) conferred somewhat greater NF-κB activation (in HEK-NEE cells) compared to the NF-κB-specific CAMP-7. We note that this initial study explored only a small set of conditions regarding timing, sequence, and magnitude of signals that can help interrogate T cell behaviors, establishing CAMPs as a tool for probing these complex phenomena in new ways.

It is interesting to consider how the CAMP mechanism compares to alternative genetic approaches for studying such phenomena and the implications for the dynamics accessible with each approach. As discussed above, VR signaling is self-propelling after autocrine or paracrine production of cytokines drives sustained VR signaling.^24^ However, abrogation or absence of sufficient cytokine production precludes VR signaling. CAMP induction provides sustained signaling until the driver of CAMP expression is removed. Both VR and CAMP signaling differ from an alternative strategy of direct overexpression of downstream transcription factors (e.g., constitutively active STAT5 or NF-κB variants); both VR and CAMP engage endogenous regulatory circuitry, driving pathway activation while preserving native feedback and attenuation mechanisms.^67, 68^ A distinguishing feature of CAMPs is that one may directly modulate the timing and duration of signal induction, either in vitro or in vitro, using externally applied cues including small molecule inducers and extensible to other approaches including optogenetics^69^. It will be useful to compare these signaling interventions with direct genetic modulation of downstream regulators of cell mechanics, including the dynamics of F-actin or myosin II, which can benefit motility and tumor control.^22, 23^ For each of these mechanisms, investigating how the timing, control, and integration of these interventions impacts short-and long-term T cell function will help to evaluate translational utility.

Although our focus in developing CAMPs was investigation of motility phenomena, it is possible that these or subsequent CAMPs could be deployed therapeutically, and here we raise related considerations. Although constitutive receptor signaling, in general, can risk exhaustion or uncontrolled activation, careful engineering of signaling strength, pathway, and duration can selectively enhance beneficial T cell functions without compromising viability or effector control.^25, 70^ Placing CAMP expression under the control of natural or engineered promoters (e.g., a NFAT reporter responsive to T cell receptor signaling) could enable engineering T cells to induce CAMP-driven behaviors in response to local environmental cues. While determining which type of CAMP activity could benefit T cell function in vivo requires further investigation, the modularity of CAMP as a genetic “part” enables its ready incorporation into synthetic biology strategies such as AND/OR logic gates, tumor-sensing modules, or environment-responsive modules to restrict timing and location of CAMP-induction, facilitating exploration of the potential utility of this approach.

There exist many opportunities to build upon the foundation established in this study, and here we highlight a few. Motility experiments used T cells from a single donor, and motility may vary by donor. An assay of the type used may underestimate motility; accumulation of cells atop the gel suggests that the most motile fraction of cells may have migrated through the matrix and arrested at the gel-medium interface where paracrine signaling is maximal, nutrient and oxygen availability is highest, and adhesive substrate necessary for continued migration is absent. Therefore, the most motile cell populations may be underrepresented in this analysis. Use of microscopy methods that directly resolve migration in the z-axis and measure trans-matrix transit could aid in evaluating this possibility. The most exciting opportunity is extension of these *in vitro* analyses to *in vivo* studies, including both the external (e.g., small molecule) and environment-induced expression strategies discussed above. CAMPs are a uniquely enabling tool for dissecting the contributions of individual pathways to interesting and important T cell motility phenomena including tumor infiltration and anti-tumor activity in preclinical models.

## Methods

### General DNA assembly

Plasmid cloning was mainly carried out using standard PCR and restriction enzyme–based methods, employing Vent DNA Polymerase (New England Biolabs, NEB), Taq DNA Polymerase (NEB), and Phusion DNA Polymerase (NEB). Restriction enzymes (NEB; Thermo Fisher), T4 DNA Ligase (NEB), Antarctic Phosphatase (NEB), and T4 Polynucleotide Kinase (PNK; NEB) were also used. In addition, Golden Gate and Gibson assembly techniques were applied where appropriate. Most plasmids were transformed into chemically competent TOP10 *E. coli* (Thermo Fisher) and cultured at 37 °C, while integration vectors were introduced into chemically competent Stable *E. coli* (NEB) and grown at 30 °C.

### Cloning CAMP vectors

CAMP plasmids for use in HEK-Blue CD122/132 and HEK-NEE cells were constructed using pcDNA-based backbones. Restriction sites were strategically selected to facilitate modular exchange of components via restriction enzyme cloning. In short, CAMP DNA containing the ECD of interest was ordered from Twist to contain a 5’ NheI site and a 3’ BbsI site. Each of the IL5Rα and IL5Rβ domain combinations used in this study were ordered as single constructs from Twist with BsaI overhangs complementary to the 5’ ECD DNA and the 3’ TNFR DNA. TNFR1/2 fusion constructs were supplied by Twist and contained a 5’ BbsI cut site complementary to the previous IL5Rα and IL5Rβ domain cut site, as well as a 3’ NotI site located after the stop codon. These three DNA fragments were independently digested by and assembled using standard restriction enzyme cloning. A comprehensive list of all plasmids generated and used in this study and their plasmid maps are provided in **Supplementary Data 2**, and corresponding plasmid maps are available in the Data Availability section.

### DNA plasmid preparation

TOP10 or NEB Stable *E. coli* cells were cultured overnight in 50 mL of LB medium containing the appropriate selective antibiotic. Plasmid DNA was then extracted using the ZymoPURE II plasmid midiprep kit (Zymo, D4201) according to the manufacturer’s instructions. In each experiment, all receptor variants being compared were prepared using the same protocol. For certain transposon vectors and reporter landing pad integration vectors, 10 mL cultures of NEB Stable *E. coli* were grown overnight in LB with the appropriate antibiotic, and plasmid DNA was isolated using the ZymoPURE plasmid miniprep kit (Zymo, D4210) following the manufacturer’s instructions.

### Golden Gate assembly of CAMP in PiggyBac Transposon Vector (PBTV) plasmids

mMoClo integration vectors were constructed using a BbsI-HF-mediated Golden Gate assembly. Each 20 µL reaction contained 2 µL 10× T4 ligase buffer, 2 µL 10× BSA (1 mg/mL), 0.8 µL BbsI-HF, 0.8 µL T4 DNA ligase (400 U/µL), 20 fmol of the integration vector backbone (pHIE426), and 40 fmol of each transcription unit and linker plasmid. The reaction was first incubated at 37 °C for 15 min, followed by 55 thermocycling iterations (37 °C for 5 min and 16 °C for 3 min). A final incubation was performed at 37 °C for 15 min, 50 °C for 5 min, and 80 °C for 10 min to terminate the reaction. The mixture was then cooled to room temperature (approximately 20–25 °C) (or held at 4 °C if run overnight), placed on ice, and immediately transformed into bacteria.

### Transient transfection of HEK-Blue CD122/132 and HEK-NEE cells

HEK293FT cells were transiently cotransfected using the calcium phosphate method. Cells were seeded at a minimum density of 1.0 × 10⁵ cells per well in 24-well plates (0.5 mL DMEM). After ∼24 h, when cells had adhered and reached ∼50% confluency, transfection was performed. Plasmid DNA (up to 500 ng for 24-well plates) was diluted in H₂O, and 2 M CaCl₂ was added to a final concentration of 0.3 M. This mixture was added dropwise to an equal volume of 2x HEPES-buffered saline (280 mM NaCl, 50 mM HEPES, 1.5 mM Na_2_HPO_4_), gently mixed by pipetting four times, incubated for 4 min, and then mixed more vigorously by pipetting ten times. Subsequently, 100 µL (24-well) of the transfection mixture was added dropwise to the cells, followed by gentle swirling of the plates.

CAMP plasmid amounts were normalized by copy number to achieve 2.5 × 10^10^ plasmids per well (24-well), corresponding to approximately 250 ng per well. Total DNA input was kept constant at 500 ng across conditions using empty vector filler DNA (pHIE298). A complete list of plasmid quantities is provided in **Supplementary Data 2**. The following day, media was aspirated and replaced with fresh medium.

### Signal quantification from HEK-Blue CD122/132 cells

Secreted embryonic alkaline phosphatase (SEAP) activity was quantified using the QUANTI-Blue colorimetric assay according to the manufacturer’s instructions. QUANTI-Blue working solution was prepared immediately before use by combining QUANTI-Blue reagent, QUANTI-Blue buffer (InvivoGen, rep-qbs), and nuclease-free water at a ratio of 1.8:1.8:176.4 (v/v/v), followed by incubation at room temperature for 10 min. Twenty microliters of cell culture supernatant were added to each well of a clear flat-bottom 96-well plate, followed by the addition of 180 μL of QUANTI-Blue working solution. Plates were incubated at 37°C for at 4 h to allow color development, after which absorbance was measured at 620 nm using a microplate reader. SEAP activity was determined from the absorbance values, with higher optical density corresponding to increased alkaline phosphatase activity. Raw absorbance values were corrected by subtracting the background signal obtained from untreated cell supernatants. The standard error of the mean (SEM) for the background-corrected values was calculated using standard error propagation. Background-corrected SEAP activity was then normalized to the dsRE2 control-transfected cells treated with 100 IU/mL IL-2 by applying a constant scaling factor such that the normalized value of the reference condition equaled 1. The corresponding SEM values were scaled by the same normalization factor.

### Signal quantification from HEK-NEE cells

Cells were harvested 36–48 h after transfection and at least 24 h after medium replacement. Cells were washed with PBS (pH 7.4), detached using 0.05% trypsin–EDTA for 5 min, and quenched with culture medium. Cell suspensions were transferred to FACS buffer (PBS pH 7.4, 2–5 mM EDTA, and 0.1% BSA), centrifuged at 150g for 5 min, and resuspended in FACS buffer containing 3 µM DAPI. HEK-NEE cells expressing mNeonGreen (mNG) and dsRedExpress2 (dsRE2) were used as compensation controls for the FITC and PE–Texas Red channels.

Flow cytometry methods are described below. For these assays, data were gated to exclude debris, doublets, and dead cells (DAPI+). dsRE2-positive cells, identified by PE–Texas Red fluorescence, were analyzed for eGFP expression in the FITC channel. Exemplary gating strategies are in **Supplementary Figures 15,16**. NF-κB signaling was quantified by measuring eGFP expression driven by the NF-κB ELAM promoter. Mean fluorescence intensity (MFI) was calculated for the relevant fluorescence channels within the transfected population and averaged across three biological replicates. Background fluorescence from dsRE2-only control cells was subtracted, and error was propagated through subsequent calculations.

eGFP MFI values were converted to molecules of equivalent fluorescein (MEFLs) using UltraRainbow calibration particles (Spherotech, URCP-100-2H) measured in each flow cytometry experiment (**Supplementary Figure 17**). Calibration curves were generated by plotting experimentally measured MFI values against manufacturer-provided MEFL values, and linear regression was used to determine the MFI-to-MEFL conversion factor. Uncertainty was propagated through the conversion calculation.

### Flow cytometry

Flow cytometry was performed using a BD LSR Fortessa special order research product (Northwestern Single Cell Genomics Core). Laser and filter configurations are listed in **Supplementary Table 6**. Approximately 3,000–10,000 single transfected cells were analyzed per sample. Transfected cells were identified using fluorescent protein reporters encoded on transfection plasmids (e.g., dsRedExpress2) or constitutive mNeonGreen expression from transposon constructs.

Data were analyzed using FlowJo version 10. Fluorescence signals were compensated for spectral overlap. Cell populations were identified by forward and side scatter, and singlets were selected using FSC-A versus FSC-H gating. Live cells were identified by exclusion of DAPI-positive cells. Transfected or engineered cells were defined based on fluorescence gates established using appropriate negative controls, including empty-vector transfections or unmodified cells, with gates set to include no more than 1% of nonfluorescent control populations.

### Cell culture

The HEK293FT cell line was obtained from Thermo Fisher, Life Technologies (RRID: CVCL_6911). HEK-Blue cells were purchased from InvivoGen (hkb-il2bg). The HEK-NEE cell line was generated in our laboratory and authenticated by treating cells with 50 nM PMA and analyzing eGFP expression via flow cytometry.

HEK293FT, HEK-Blue CD122/132, and HEK-NEE cells were maintained in DMEM (Gibco, 31600-091) supplemented with 10% FBS (Gibco, 16140-071), 6 mM L-glutamine (2 mM from Gibco, 31600-091; 4 mM from Gibco, 25030-081), penicillin (100 U/µL), and streptomycin (100 µg/mL; Gibco, 15140-122) at 37 °C in a 5% CO₂ incubator. Cells were subcultured every 2–4 d at a 1:5 to 1:20 ratio using trypsin–EDTA (Gibco, 25300-054). All three HEK cell lines were cultured under these identical conditions.

HEK293FT, HEK-Blue CD122/132, and HEK-NEE cell lines were confirmed Mycoplasma-free using the MycoAlert Mycoplasma Detection Kit (Lonza, LT07-318).

### Primary human T cell culture

Primary human T cells were obtained from Stemcell Technologies as quarter leukopaks (70500.2). These cells were immediately processed upon receipt using Stemcell EasySep Pan-T Cell isolation kit (17951). T cells were cryopreserved in Cryostor CS10 (Stemcell Technologies, 100-1061) cryogenic preservation medium. After thawing, T cells were cultured in Immunocult-XF T Cell Expansion Medium (Stemcell Technologies, 10981), supplemented with 1% penicillin (100 U/µL) and streptomycin (100 µg/mL; Gibco, 15140-122). After thawing and during passage of cells without the 4αβ chimeric cytokine receptor, culture medium was supplemented with 20 IU/mL IL-2 (Peprotech 200-02-50UG). Cultures undergoing selection with the 4αβ chimeric cytokine receptor were supplemented with 30 ng/mL IL-4 (Peprotech, 200-04-250UG).

### Primary human T cell electroporation

Primary human T cells were electroporated one day after thawing. T cells were removed from their culture vessels and centrifuged at 200 x g for 10 minutes. Culture media was aspirated, and cells were resuspended in Lonza electroporation buffer P3 using 20 µL of buffer per 1 million cells. 1 million T cells were electroporated per well using a Lonza 4D 16-well electroporation system with pulse code FI-115. Unless otherwise indicated, 600 ng of transposon vector was electroporated per reaction, along with 240 ng of Hyperactive PiggyBac (HyPB) transposase plasmid, for a total of 840 ng of DNA per reaction. Two days after electroporation, cells were stimulated using 25 µL/mL CD3/CD28 Immunocult Activator (Stemcell Technologies, 10971) and cultured in Immunocult-XF T Cell Expansion Medium (Stemcell Technologies, 10981) supplemented either with 20 IU/mL IL-2 (Peprotech 200-02-50UG) or 30 ng/mL IL-4 (Peprotech, 200-04-250UG) for cultures undergoing selection with the 4αβ chimeric cytokine receptor.

### T cell migration studies

For experiments examining the effects of cell density or PMA treatment, cells receiving stimulation were treated with 25 µL/mL ImmunoCult™ Human CD2/CD3/CD28 T Cell Activator (Stemcell Technologies, 10970) either 24 h before or immediately before collagen encapsulation, as indicated for each experiment. T cells were maintained in ImmunoCult™-XF T Cell Expansion Medium (Stemcell Technologies, 10981) supplemented with 100 IU/mL IL-2 (PeproTech, 200-02-50UG).

Collagen gels were prepared at a final collagen concentration of 2 mg/mL using rat tail collagen type I (Corning, CLS354236). Because the supplied collagen stock concentration varied between lots (typically 3.45–3.75 mg/mL), reagent volumes were adjusted accordingly to maintain the final collagen concentration and gel volume. For collagen supplied at 3.45 mg/mL, 105 µL of T cells (5E5 cells, unless otherwise specified) was mixed with 105 µL of 0.22 µm-filtered reconstitution buffer (100 mL Milli-Q water, 4.8 g HEPES, and 2.2 g sodium bicarbonate). This mixture was combined on ice with 290 µL of collagen and mixed thoroughly by pipetting. For collagen supplied at 3.75 mg/mL, 115 µL of T cells (5E5 cells, unless otherwise specified) was mixed with 115 µL of reconstitution buffer and combined on ice with 270 µL of collagen. In both cases, collagen was neutralized with 1 M NaOH (1% v/v) immediately before polymerization, yielding a final gel volume of 500 µL containing 2 mg/mL collagen. The resulting 500 µL mixture of 2 mg/mL collagen gel was gently pipetted to avoid bubble formation and evenly distributed across the bottom of a 12-well plate. Gels were allowed to polymerize for 1 h at 37 °C in a 5% CO₂ incubator.

After polymerization, 0.5 mL of Immunocult™-XF medium supplemented with 100 IU/mL IL-2 (Peprotech, 200-02-50UG) was added to the top of the gels. For experiments with doxycycline-inducible CAMP constructs, the indicated masses of doxycycline were added to cells 48 h prior to gel encapsulation to ensure sufficient expression of CAMP proteins. 48 h after gel encapsulation, imaging was performed using a Keyence BZ-X800E microscope with the timelapse software suite. Plates were maintained in a stage-top incubator (TOKAI HIT, INU-K1W-F1) at 37 °C, 5% CO₂, and controlled humidity. Phase-contrast images were acquired every 3 min for 1 h using a 20X objective. A Z-stack of 11 images over a vertical axis range of 50 µm was taken with every image capture.

### T cell migration studies with CAMP induction

In experiments where CAMP constructs were induced with doxycycline, CAMP-engineered T cells at least 2 weeks-post CD3/CD28 Immunocult™ stimulation (Stemcell Technologies, 10971) were prepared as described above for 3D migration assays unless otherwise indicated. Cells were encapsulated in 2 mg/mL collagen gels at a density of 5×10^5^ live cells/gel. Following 1 h of polymerization at 37°C and 5% CO₂, 0.5 mL of Immunocult™-XF supplemented with 100 IU/mL IL-2 and the indicated doxycycline concentration was added to the top of each gel to induce CAMP expression. 48 h after gel encapsulation, imaging was performed using a Keyence BZ-X800E microscope with the timelapse software suite. Plates were maintained in a stage-top incubator (TOKAI HIT, INU-K1W-F1) at 37 °C, 5% CO₂, and controlled humidity. Images were acquired every 2 min for 1 h using a 20X objective lens with both phase contrast microscopy and fluorescence microscopy using a GFP filter cube (Keyence, OP-87763). A Z-stack of 11 images over a vertical axis range of 50 µm was taken with every image capture.

### Spot detection and single-cell tracking

Images were processed using the Keyence BZ-X800 Analyzer software to generate .AVI file videos that could be processed for cell tracking. Briefly, *.gci files from each well were processed using the Time Lapse analysis suite. To remove image borders interfering with downstream analysis, image stabilization was disabled during processing of image series. Time-lapse sequences were acquired at 11 Z-planes every 2 min for 1 h (30 frames) for doxycycline-inducible T cell experiments, and every 3 min for 1 h (20 frames) for experiments involving T cells that were not engineered. Best-focus composite images from these processed Z-stacks were exported as AVI stacks using the Keyence BZ-X analysis software. The spatial calibration was 1.02939 µm/pixel for images collected for experiments containing doxycycline-inducible T cells, and 0.68626 µm/pixel for images collected using T cells that were not engineered. These calibration values were verified against a stage micrometer. Because mean-squared displacement scales with the square of this calibration, each channel was analyzed with its own measured pixel size. These images were subsequently converted into *.AVI videos to be analyzed in FIJI image analysis software using the TrackMate plugin.

Time-lapse stacks were imported into Fiji/ImageJ^55^ and analyzed with TrackMate v7^53, 54^. In each frame, cells were detected with the Laplacian-of-Gaussian detector. Within time-lapse videos with a spatial calibration of 0.68626 µm/pixel and a capture interval of 3 min, detection parameters included an estimated object diameter of 36 µm. For time-lapse videos with a spatial calibration of 1.02939 µm/pixel and a capture interval of 2 min, detection was achieved using an estimated object diameter of approximately 20 µm for the GFP channel and 24 µm for phase contrast (corrected by a factor of 1.5X from the 0.68626 µm/pixel videos), with median filtering and sub-pixel localization enabled. A quality threshold was applied to exclude background and sub-cellular debris; the threshold was selected by inspection of the spot-quality distribution and confirmed by overlaying detected spots on the source images. Detected spots in videos with a spatial calibration of 1.02939 µm/pixel were linked into single-cell trajectories using the Linear Assignment Problem tracker (LAP)^71^ with a maximum frame-to-frame linking distance of approximately 45 µm, a gap-closing distance of approximately 55 µm, and a maximum frame gap of 2 (permitting bridging across a single consecutive missed detection). For videos with a spatial calibration of 0.68626 µm/pixel and a capture interval of 2 min, these parameters were divided by a factor of 1.5X to adjust for image pixel size and 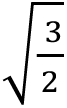 to account for MSD time lag differences, yielding a corrected linking distance of 25 µm, a gap-closing distance of approximately 30 µm, and a maximum frame gap of 2. Track-segment splitting and merging were disabled, as T cells do not divide over this timescale. Spot centroid coordinates were exported in calibrated (µm) units for downstream analysis. Identical detection and tracking parameters were used across all conditions and replicates within a given channel and imaging run.

### Image analysis pipeline automation

The cell tracking pipeline was automated using agentic coding tools (**Supplementary Notes 3,4**). The manual TrackMate workflow in the Fiji GUI was documented as an annotated screenshot series (movie import, LoG spot detection, quality filtering, and LAP tracking). This was fed into Claude Code (Opus 4.7, xhigh effort) with additional user guidance, which reproduced it exactly as a headless Jython macro (trackmate_batch.py) that exports per-spot statistics, track overlays, and the reopenable TrackMate session for each input movie. The macro was wrapped in a Makefile that autodetects the Fiji installation, discovers input .avi files across flat or nested folder layouts, and runs the tracking-then-MSD stages end-to-end, enabling reproducible, one-command (i.e. make -B parameter=new_value) reprocessing. First, the automated pipeline was adjusted and demonstrated to match the manual results. This pipeline was then used to explore and improve choices for detection and tracking parameters (e.g., spot-quality threshold) with the context of how they affect entire datasets, as opposed to the single-sample scale of manual Fiji processing.

### Mean-squared displacement calculation

For each trajectory in the doxycycline-induced GFP fluorescence images, the two-dimensional mean-squared displacement (MSD) was computed at a fixed time lag τ of 10 min (5 frames) as the time-averaged squared displacement over all overlapping windows of that lag within the track, MSD(τ) = ⟨ |r(t + τ) − r(t)|² ⟩, where r(t) is the cell centroid position at time t and the average is taken over all frame pairs separated by τ for which both endpoints were detected. For each trajectory in the initial phase contrast microscopy study, MSD calculations utilized a fixed time lag of 9 min (3 frames) with each frame collected every 3 min.

All trajectories yielding at least one valid τ-lag window were included; no minimum track-length filter was applied, so that faster cells, which are more prone to premature track termination, were not systematically excluded. Positions were converted to micrometers prior to computing displacements, and MSD is reported in µm². Because imaging captured a single two-dimensional plane of a three-dimensional matrix, displacements reflect the in-plane projection of cell motion. MSD computation was implemented in Python 3 using NumPy and pandas.

### Motility classification and motile fraction

Consistent with the definition of a motile cell as one that translocates more than approximately its own radius over the observation window, cells were classified as motile when their 10-min-lag MSD exceeded R² = 25 µm², where R = 5 µm approximates the T-cell radius. For each well, the motile fraction was computed as the percentage of tracked cells meeting this criterion. The motile fraction, rather than the population median or mean MSD, was used as the primary motility metric as the per-cell MSD distributions spanned several orders of magnitude and were bimodal, comprising an arrested subpopulation near the localization limit and a distinct motile subpopulation; in this two-state regime, a central-tendency statistic reflects the relative size of the two subpopulations rather than a characteristic cell behavior, whereas the motile fraction directly quantifies the property under study and is stable across wells as a proportion.

### Gaussian mixture model characterization

Because variants and even replicate wells of the same variant differed substantially in their mean motility, a fixed MSD threshold (25 µm², adopted from Johnston et. al^24^) classified cells inconsistently across wells; modest well-to-well shifts in the overall distribution moved fractions of cells across the cutoff, inflating replicate variability in the estimated motile fraction, adding to the demonstrated cell density effects. Therefore, we used an alternative approach in which we assumed that the overall population comprised two subpopulations (high and low motility), but the average motility of each subpopulation was assumed to (potentially) vary between conditions. To this end, we used a two-component Gaussian mixture model, modeling the log₁₀-transformed per-cell time-averaged MSD as a distribution comprising low-and high-motility states, fit by expectation-maximization. Settings used: msd_overlap_partial; phase contrast images used a lag of 3 frames (3 min/frame => 540 s); GFP channel images used a lag of 5 frames (2 min/frame => 600s); cells below 0.1 µm² were excluded as localization-floor artifacts. The component width was estimated based on maximum likelihood across all cells in each channel’s full dataset, yielding: σ= 0.373 (95% CI 0.356–0.386) for phase contrast images, σ = 0.309 (0.294–0.326) for GFP channel images, in log₁₀ units; then the value was held fixed during per-well fitting. Width was modelled as biological state width in addition to a per-cell measurement term whose magnitude follows from the number of independent lag windows in that cell’s track (the diffusivity-independent CV expected for a time-averaged MSD estimator^72–75^), and σ was estimated by maximum likelihood jointly with the per-well means. Fixing rather than freely fitting this width prevented the low-motility component from broadening to absorb the cells of intermediate MSD, which we interpret not as a distinct third state but as cells that spent a moderate proportion of the imaging time course in each of the two states and therefore fall between the two population means. Fits are per-well with free means and weights; the component width is the only fixed parameter, and it is fixed per channel (phase contrast or GFP channel), not per condition. Therefore, the per-well high-motility fraction (the mixture weight) is a threshold-free readout robust to whole-well shifts in mean motility.

### Cell extraction from collagen gel and preparation for RNA-Sequencing

T cells encased in collagen gel and analyzed via time-lapse microscopy were liberated from the gel using a 2 mg/mL solution of Collagenase IV (Gibco, 17104-019). Cellular and gel debris were removed by wash and centrifugation, and resulting cell pellets were lysed and stored in 50 µL of ZymoShield (Zymo, R1100). Samples were then submitted to Plasmidsaurus for RNA-Sequencing. For each construct, untreated and 500 ng/mL doxycycline-treated samples were sequenced, with additional PMA-treated samples for dox-inducible LumiScarlet cells and naive cells, yielding 12 libraries per run. The full experiment was performed across two independent engineering-and-sequencing runs derived from a single donor, for a total of 24 libraries.

### Read processing and gene-level quantification

Sequencing reads were processed using the Plasmidsaurus RNA-seq analysis pipeline. FASTQ quality was assessed with FastQC v0.12.1, followed by read trimming and filtering using fastp v0.24.0 with poly-X tail trimming, 3′ quality-based trimming, a minimum Phred score of 15, and a minimum read length of 50 bp. Filtered reads were aligned to the reference genome using STAR v2.7.11 with non-canonical splice junction removal and unmapped read output, followed by coordinate sorting with samtools v1.22.1. PCR and optical duplicates were removed using UMI-based deduplication with UMIcollapse v1.1.0. Alignment metrics, strand specificity, and genomic feature distribution were assessed using RSeQC v5.0.4 and Qualimap v2.3 and summarized with MultiQC v1.32. Gene-level counts were generated using featureCounts (Subread v2.1.1) with strand-specific counting, fractional assignment of multi-mapping reads, exon and 3′ UTR features, and grouping by gene_id. Counts were annotated with gene biotype and additional metadata from the reference GTF file. Inter-sample correlation, read mapping statistics, and principal component analysis are shown in **Supplementary Figure 14.**

Differential expression between doxycycline-induced (500 ng/mL) or PMA-treated (0.4 ng/mL) and untreated cells was determined per construct across both sequencing runs (n = 2 per condition; run-to-run concordance shown in **Supplementary Figure 13**) using edgePython (v0.2.5), a Python implementation of edgeR, with TMM normalization and the negative-binomial quasi-likelihood F-test; genes with Benjamini–Hochberg FDR < 0.05 and |log2 fold change| > 1 were considered differentially expressed. Pre-ranked gene set enrichment analysis was performed with GSEApy (v0.12) against the MSigDB Hallmark, Reactome, and Gene Ontology Biological Process collections, with gene sets at FDR q < 0.25 considered enriched. Analyses were performed in Python using NumPy, pandas, SciPy, scikit-learn, and Matplotlib, as well as on the Plasmidsaurus RNA-Seq analysis portal.

## Abbreviations

AP-1: Activator protein 1 (transcription factor)
BCL3: BCL3 transcription coactivator (B-cell lymphoma 3-encoded protein)
BIRC3: Baculoviral IAP repeat-containing protein 3
BSA: Bovine serum albumin
BZ-X: Keyence BZ-X series all-in-one fluorescence microscope
BZ-X800E: Keyence BZ-X800E fluorescence microscope controller/analyzer unit
C7R: Constitutively active IL-7 cytokine receptor
CAG: CMV early enhancer/chicken β-actin promoter
CAMPs: Constitutive Activators of Motility-associated Pathways
CAR: Chimeric antigen receptor
CCL: C-C motif chemokine ligand
CD: Cluster of differentiation
CD34-IL7R: CD34–interleukin-7 receptor fusion construct
CD122/132: IL-2 receptor β chain (CD122) and common γ chain (CD132)
CDK1: Cyclin-dependent kinase 1
cHS4: Chicken hypersensitive site 4 insulator element
CISH: Cytokine-inducible SH2-containing protein
CLS354236: Corning Collagen I, rat tail (Sigma-Aldrich catalogue number)
COX-2: Cyclooxygenase-2
CPM: Counts per million
CS10: CryoStor 10% DMSO
CSF2: Colony-stimulating factor 2 (gene encoding GM-CSF)
CXCL8: C-X-C motif chemokine ligand 8 (interleukin-8)
DAPI: 4′,6-diamidino-2-phenylindole
DMEM: Dulbecco’s Modified Eagle Medium
DMSO: Dimethyl sulfoxide
DNA: Deoxyribonucleic acid
dsRE2: dsRedExpress2
E2F1: E2F transcription factor 1
EBI3: Epstein-Barr virus-induced gene 3 (IL-27/IL-35 subunit)
ECD: ectodomain
EDTA: Ethylenediaminetetraacetic acid
EF1α: Elongation factor 1-alpha (promoter)
eGFP: Enhanced green fluorescent protein
EGR2: Early growth response protein 2
ELAM: Endothelial leukocyte adhesion molecule (E-selectin); ELAM minimal promoter
ERK: extracellular signal-regulated kinase
FACS: Fluorescence-activated cell sorting
FBS: Fetal bovine serum
FDR: False discovery rate
FITC: Fluorescein isothiocyanate
FSC-A: Forward scatter area
FSC-H: Forward scatter height
G4S: Glycine–serine linker, (Gly₄Ser)ₙ
GFP: Green fluorescent protein
GM-CSF: Granulocyte-macrophage colony-stimulating factor
GMM: Gaussian Mixture Model
GMP: Good Manufacturing Practice
GSEApy: GSEApy — Python implementation of Gene Set Enrichment Analysis
GTF: Gene transfer format (genome annotation file format)
GUI: Graphical user interface
GZMB: granzyme B
HEK: Human embryonic kidney 293 (cells)
HEK293FT: Human embryonic kidney 293FT (cells)
HEKNEE: HEK NFκBELAMeGFP
HEPES: 4-(2-hydroxyethyl)-1-piperazineethanesulfonic acid
HyPB: Hyperactive PiggyBac
IFNG: Interferon gamma (gene symbol)
IFNGR: interferon gamma receptor
IFNγ: Interferon gamma (protein)
IKK: IκB kinase
IKKα/β/γ: IκB kinase subunits α, β and γ (γ = NEMO)
IL: Interleukin
IL2Rβ: IL-2 receptor β
IL2Rγ: IL-2 receptor γ
IL5R: IL-5 receptor
IL5R/TNFR: IL-5 receptor / tumor necrosis factor receptor (chimera)
IL5Rβ: IL-5 receptor β chain (common β chain, CSF2RB)
IL7R: Interleukin-7 receptor (α chain, CD127)
IL10RB: Interleukin-10 receptor subunit beta (gene symbol)
IL10Rβ: Interleukin-10 receptor β subunit (protein)
ITR: Inverted terminal repeat
IκB: Inhibitor of κB
JAK: Janus kinase
JMD: juxtamembrane domain
JUN: Jun proto-oncogene, AP-1 transcription factor subunit
KLF2: Krüppel-like factor 2
LAP: Linear Assignment Problem tracker
LB: Lysogeny broth (Luria-Bertani medium)
LoG: Laplace of Gaussian
log2FC: Log₂ fold change
LSR: BD LSR-series flow cytometer
MAPK: mitogen-activated protein kinase
MEFLs: molecules of equivalent fluorescein
MFI: Mean fluorescence intensity
Milli-Q: Milli-Q ultrapure water (Millipore/Merck)
MKI67: Marker of proliferation Ki-67
mMoClo: Mammalian Modular Cloning
mNG: mNeonGreen
MSD: mean squared displacement
MSigDB: Molecular Signatures Database
MYC: MYC proto-oncogene, bHLH transcription factor
NEB: New England Biolabs
NF-kB: Nuclear factor kappa-light-chain-enhancer of activated B cells
NFAT: Nuclear factor of activated T cells
NFKB2: Nuclear factor kappa B subunit 2 (p100/p52)
NFKBIA: NF-κB inhibitor alpha (IκBα)
NFKBIE: NF-κB inhibitor epsilon (IκBε)
NR4A3: Nuclear receptor subfamily 4 group A member 3
OP-87763: Keyence BZ-X GFP filter cube (catalogue number)
Pan-T: Pan-T cell (total CD3⁺ T cells)
PBS: Phosphate-buffered saline
PBTV: PiggyBac Transposon Vector
PCA: Principal Component Analysis
pcDNA: pcDNA plasmid expression vector series (Invitrogen)
PCR: Polymerase chain reaction
PI3K: phosphoinositide 3-kinase
PiggyBac: PiggyBac transposon system
PKC: protein kinase C
PMA: phorbol 12-myristate 13-acetate
PNK: Polynucleotide kinase
PTGS2: Prostaglandin-endoperoxide synthase 2 (COX-2)
pTRE-3G: pTRE-3G plasmid — third-generation tetracycline response element promoter
QUANTI-Blue: QUANTI-Blue colorimetric SEAP detection reagent (InvivoGen)
RNA: Ribonucleic acid
RNA-Seq: RNA sequencing
RRID: Research Resource Identifier
RΔ: TNFR1 death domain deletion variant
S1PR1: Sphingosine-1-phosphate receptor 1
S-S: Disulfide bond
SEAP: Secreted embryonic alkaline phosphatase
SELL/CD62L: Selectin L / L-selectin (CD62L)
SOCS2: Suppressor of cytokine signaling 2
SSC-A: Side scatter area
STAR: Spliced Transcripts Alignment to a Reference (RNA-seq aligner)
STAT5: Signal transducer and activator of transcription 5
STAT5b: Signal transducer and activator of transcription 5B
T2A: Thosea asigna virus 2A self-cleaving peptide
Tet-On: Tetracycline-inducible expression system
TMD: transmembrane domain
TME: tumor microenvironment
TMM: Trimmed mean of M-values (normalization method)
TNF: Tumor necrosis factor
TNFAIP3/A20: TNF alpha-induced protein 3 (A20)
TNFR: tumor necrosis factor receptor
TNFR1: Tumor necrosis factor receptor 1 (TNFRSF1A, p55/CD120a)
TNFR2: Tumor necrosis factor receptor 2 (TNFRSF1B, p75/CD120b)
TNFRDD: TNFR death domain
TNFRSF9/4-1BB: TNF receptor superfamily member 9 (4-1BB, CD137)
TNFRSF18/GITR: TNF receptor superfamily member 18 (glucocorticoid-induced TNFR-related protein)
TNFRΔDD1: TNFR1 death domain deletion variant
TOP2A: DNA topoisomerase II alpha
TRADD: Tumor necrosis factor Receptor Type 1-Associated DEATH Domain
TRE: Tet response element
tTA: Tetracycline-controlled transactivator
TU: Transcriptional unit
UMI: Unique molecular identifier
UTR: Untranslated region
VR: Velocity Receptors
XF: Xeno-Free

## Author contributions

Y.R.S and J.N.L conceptualized the project. Y.R.S, M.R.K, and E.C.F created reagents, designed, and performed experiments. Y.R.S, M.R.K, E.C.F, and J.W created reagents for the experiments. Y.R.S and X.S.B analyzed the data. Y.R.S drafted the original manuscript, and Y.R.S, X.S.B., and M.R.K created the figures. N.M and D.J.O provided intellectual contributions to the analysis pipeline and assay design. J.N.L. supervised the work. All authors edited and approved the final manuscript.

## Conflict of interest

Y.S. and J.L. have applied for patent protection of inventions related to this study.

## Supporting information

**SupplementaryInformation.pdf**: additional figures, notes, and tables

**SupplementaryData1.zip** RNA-seq analysis files

**SupplementaryData2.zip**: plasmid maps, list of all plasmids generated in this study, and plasmid doses used in all experiments

**SupplementaryData3.zip** source data use to generate figures

**Additional Data**: In addition to the supplementary data discussed above, an associated Zenodo deposit^76^ contains: (1) Raw flow cytometry files; (2) Time lapse microscopy movies used for motility analyses; and (3) Plasmid maps in Snapgene format for plasmids used in this study.

**Code availability**.

Software used to analyze cell motility is available in the project repository at https://github.com/leonardlab/camps. The repository may be updated over time; version 1.0.0 was published with this study. Data in this paper were analyzed using the software versions and settings given below.

Cell detection and tracking were performed in Fiji/ImageJ 1.54p using TrackMate v8.1.6, run headlessly via the Jython script stage1_trackmate.py (Jython 2.7.4) under Java 21.0.7 (Azul Zulu) on macOS 26.5.2. Spots were detected with the Laplacian-of-Gaussian detector on channel 1, with sub-pixel localization and median filtering enabled, a detector threshold of 0.2 and no initial quality filter; the detector radius was 18.0 px for phase-contrast data and 10.0 px for GFP data. Spots were linked with the Sparse LAP tracker using a maximum frame gap of 2 frames, with track splitting and merging disabled; maximum linking and gap-closing distances were 46.0 px and 55.0 px respectively for phase-contrast data, and 25.0 px and 30.0 px for GFP data.

Per-cell mean squared displacement was computed with compute_msd.py under Python 3.11.4 with numpy 2.4.6 and pandas 2.3.3, using a pixel calibration of 0.68626 µm/px and a lag of 3 frames (540 s) for phase-contrast data, and 1.02939 µm/px and a lag of 5 frames (600 s) for GFP data.

Gaussian-mixture fitting, statistical testing and figure generation were performed under Python 3.11.15 with numpy 2.4.6, pandas 2.3.3, scipy 1.17.1, statsmodels 0.14.6, scikit-learn 1.9.0 and matplotlib 3.11.0. An environment specification is provided in the repository.

## Acknowledgements

This work was supported in part by the National Cancer Institute of the National Institutes of Health under Award Number F30CA275362 and by the National Institute of General Medical Sciences of the NIH under Award Number T32GM008152 (to Hossein Ardehali). Xavier Bower and Yannick Schreiber were supported in part by the National Institutes of Health Training Grant (T32GM153505 and T32GM008449, respectively) through Northwestern University’s Biotechnology Training Program. Flow cytometry was performed at the Single Cell Genomics Facility at Northwestern University (RRID:SCR_026652), graciously supported by the Department of Neurobiology, the Department of Molecular Biosciences, and the Northwestern University Office for Research. The content is solely the responsibility of the authors and does not necessarily represent the official views of the National Institutes of Health.

## Notes

https://doi.org/10.5281/zenodo.22073097

