## Supplementary Information for "Genetically inducible coordinators of cytokine signaling pathways for interrogating T cell motility"

**Contents:**

Supplementary Figures 1–18
Supplementary Notes 1–3
Supplementary Tables 1–5
References Cited in Supplementary Information

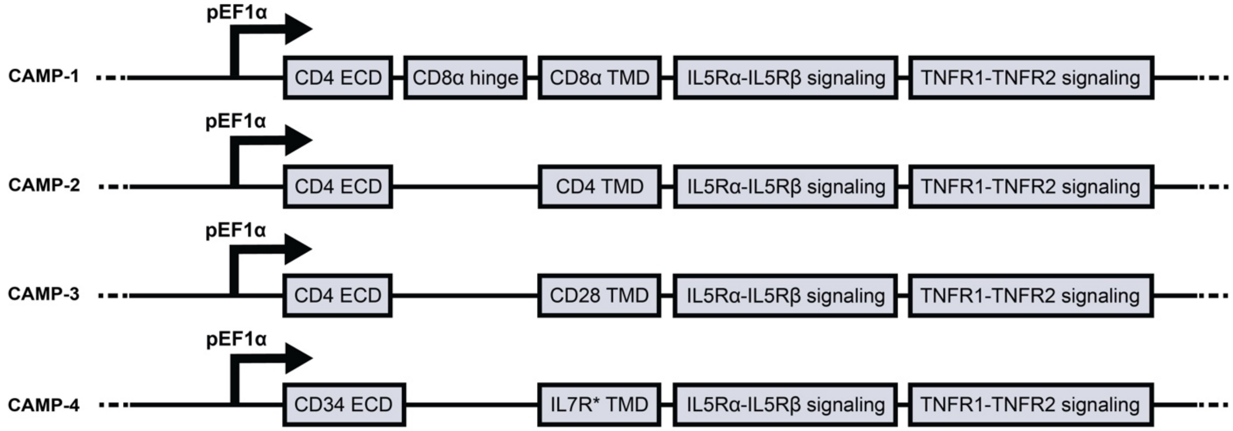


**Supplementary Figure 1. Initial CAMP constructs.** Schematic of the first four CAMP architectures (CAMP-1 to CAMP-4). Each construct is expressed from a pEF1α promoter. All constructs share a common intracellular backbone consisting of tandem IL5Rα-IL5Rβ and TNFR1-TNFR2 signaling modules, and they differ in their extracellular and transmembrane components. CAMP-1 combines a CD4 ECD with a CD8α hinge and CD8α TMD; CAMP-2 and CAMP-3 pair the CD4 ECD directly with a CD4 or CD28 TMD, respectively; and CAMP-4 pairs a CD34 ECD with an IL7R* TMD. Boxes denote protein domains and are not drawn to scale. CAMP: Constitutive Activator of Motility-associate Pathways; pEF1α: human elongation factor-1α promoter; CD4: cluster of differentiation 4; ECD: extracellular domain; CD8α: cluster of differentiation 8α; TMD: transmembrane domain; IL5Rα: interleukin-5 receptor α subunit; IL5Rβ: interleukin-5 receptor β subunit: TNFR1: tumor necrosis factor receptor 1: TNFR2: tumor necrosis factor receptor 2; CD28: cluster of differentiation 28. CD34: cluster of differentiation 34; IL7R*: interleukin-7 receptor subunit α transmembrane domain with Thr244-Ile245insCysProThr mutation.

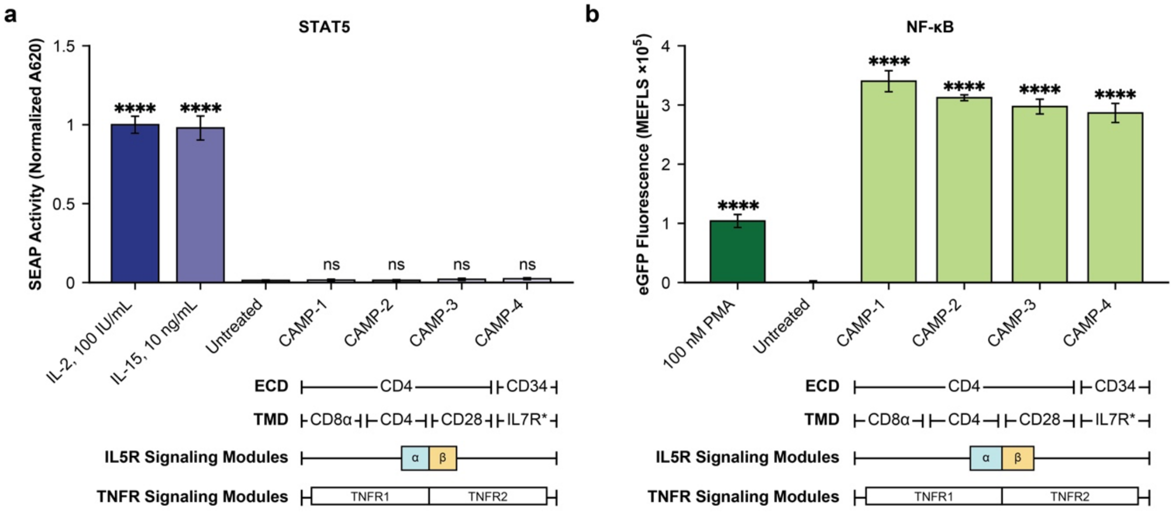


**Supplementary Figure 2. Repeat of experiment shown in Figure 1c,d. (a,b)** Induction of STAT5 (a) and CAMP (b) signaling via the initial set of CAMP designs. Cytokine controls (IL-2, IL-15) induce STAT5 activity, and PMA induces NF-κB. Bars show mean ± s.e.m. (n = 3); significance is versus the untreated condition by one-way ANOVA, followed by Dunnett's multiple comparisons test to evaluate specific comparisons (****p ≤ 0.0001; ns, not significant), under the assumption of equal variance. CAMP: Constitutive Activator of Motility-associated Pathways; VR: Velocity Receptor, ECD: extracellular domain, TMD: transmembrane domain, STAT5: signal transducer and activator of transcription 5, SEAP: secreted embryonic alkaline phosphatase (normalized A620, absorbance at 620 nm), NF-κB: nuclear factor κB, eGFP: enhanced green fluorescent protein, PMA: phorbol 12-myristate 13-acetate, MEFL: molecules of equivalent fluorescein.

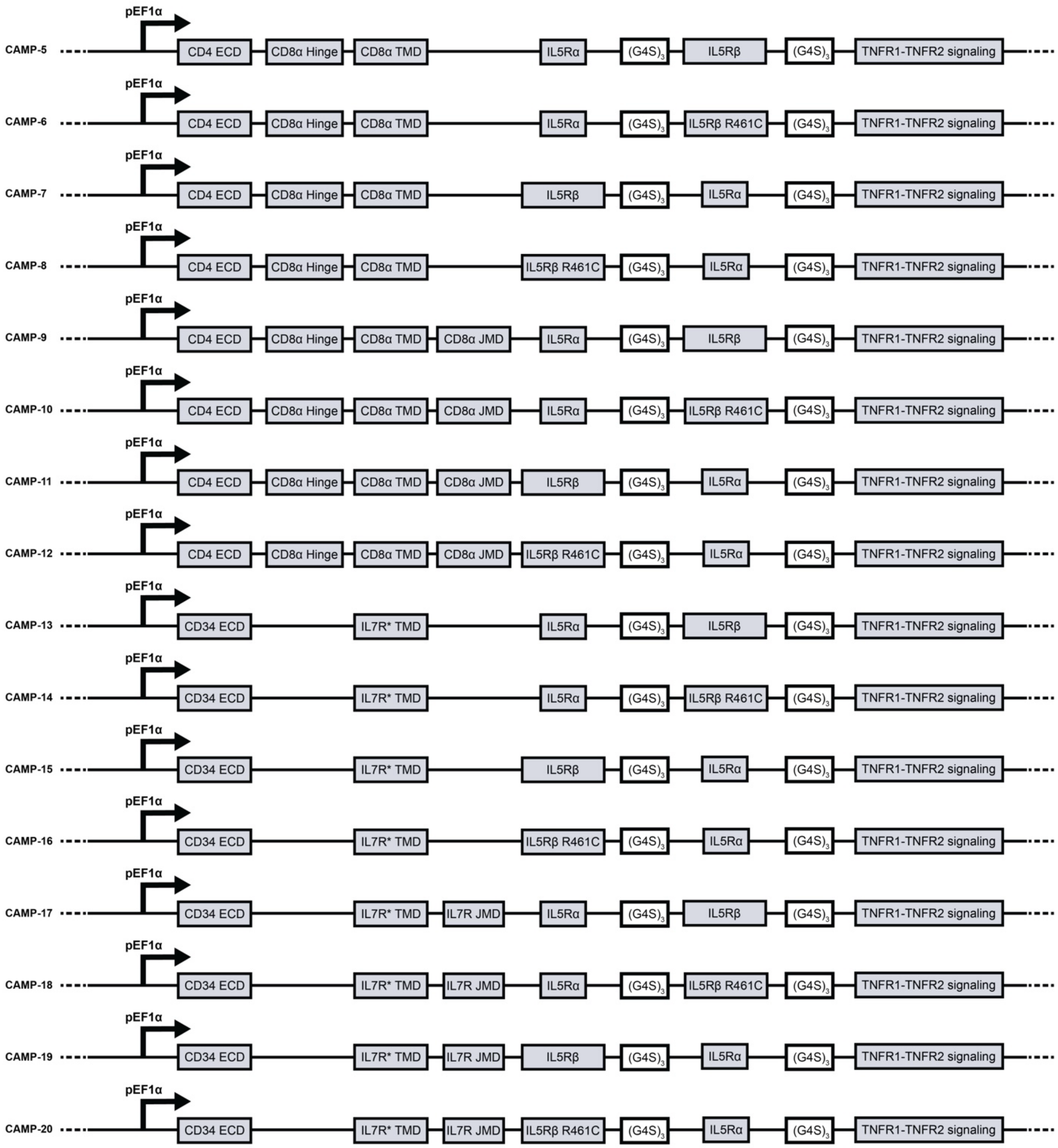


**Supplementary Figure 3. Revised CAMP designs focusing on identifying requirements for inducing STAT5 signaling.** Schematic of CAMP-5 through CAMP-20 architectures. Each construct is expressed from a pEF1α promoter and comprising a transmembrane domain, tandem IL5Rα-IL5Rβ domains joined by $(G4S)_3$ linkers, and a TNFR1-TNFR2 signaling module. The 16 designs represent all combinations of four binary features: the transmembrane domain (CD4 ECD with CD8α hinge and TMD, vs CD34 EVD with IL7R* TMD), presence of a JMD, the order of IL5Rα and IL5Rβ, and wild-type vs. R461C mutant IL5Rβ. Boxes denote protein domains, and are not drawn to scale. $(G4S)_3$: GGGGSGGGGSGGGGS linker, JMD: juxtamembrane domain. Remaining abbreviations are as defined in **Supplementary Figure 1** (initial CAMP constructs).

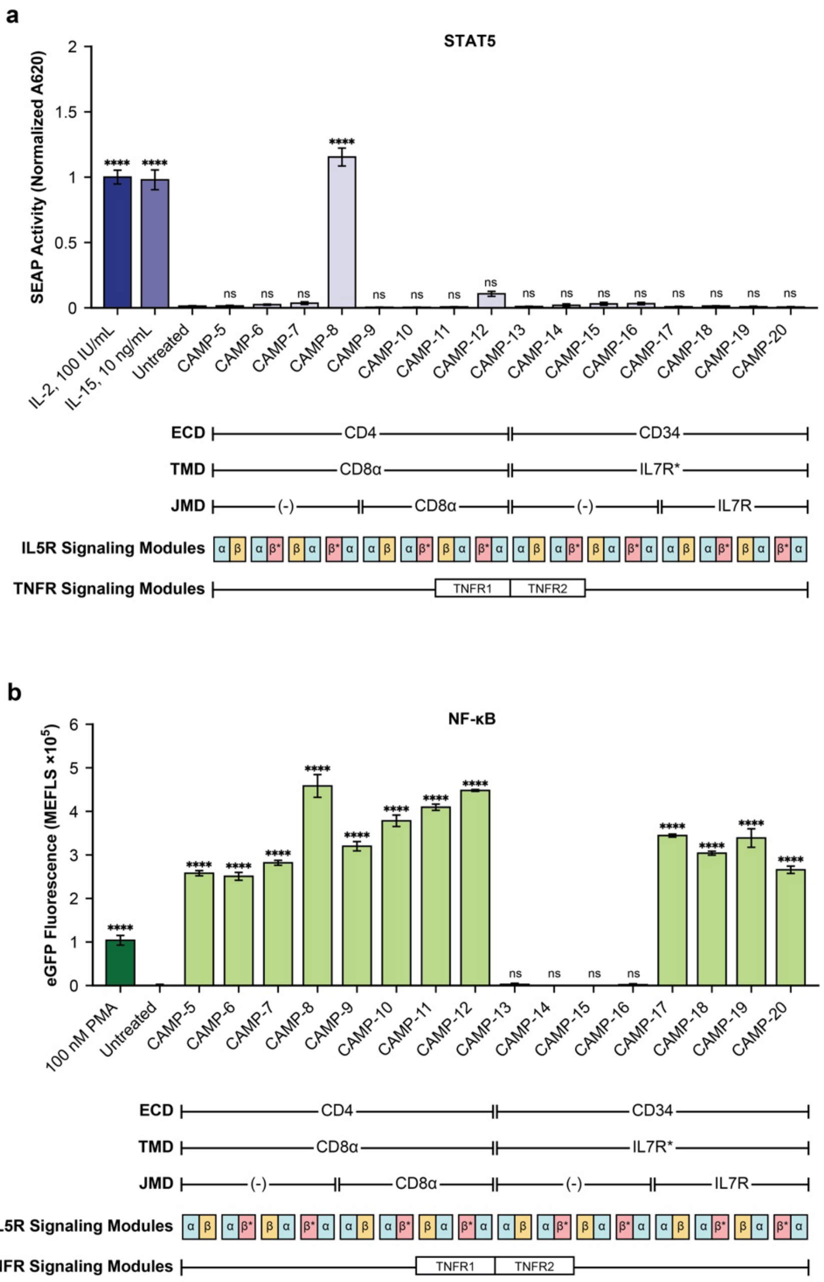


**Supplementary Figure 4. Repeat of experiment show in Figure 2c,d. (a,b)** Induction of STAT5 (a) and NF-κB (b) signaling via new CAMP designs. activity. Bars show mean ± s.e.m. (n = 3); significance is versus the untreated condition by one-way ANOVA, followed by Dunnett's multiple comparisons test to evaluate specific comparisons (****p ≤ 0.0001; ns, not significant), under the assumption of equal variance. ECD: extracellular domain; TMD: transmembrane domain, JMD: juxtamembrane domain, IL5R: interleukin-5 receptor, TNFR: tumor necrosis factor receptor, SEAP: secreted embryonic alkaline phosphatase (normalized A620), NF-κB: nuclear factor κB, PMA: phorbol 12-myristate 13-acetate, MEFL: molecules of equivalent fluorescein.

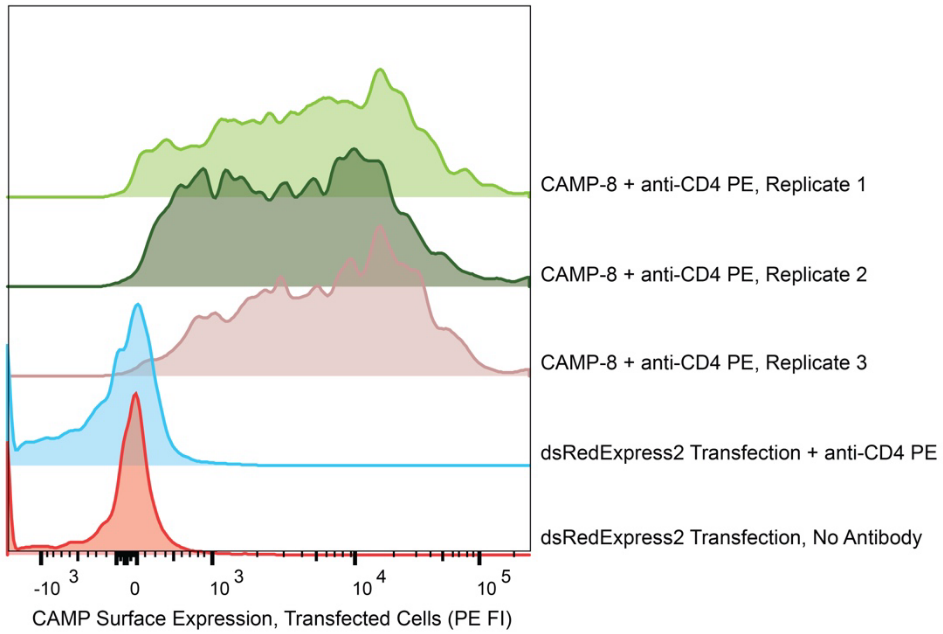


**Supplementary Figure 5. CAMP-8 is expressed on the surface of HEK293FT cells.** HEK293FT cells transfected with mNeonGreen plasmid and CAMP-8 were subsequently stained for CD4 ECD domain using anti-CD4-PE antibody. Flow cytometry gates filtered for cells, single cells, live cells ($DAPI^-$), and transfected cells ($mNeonGreen^+$, shown here).

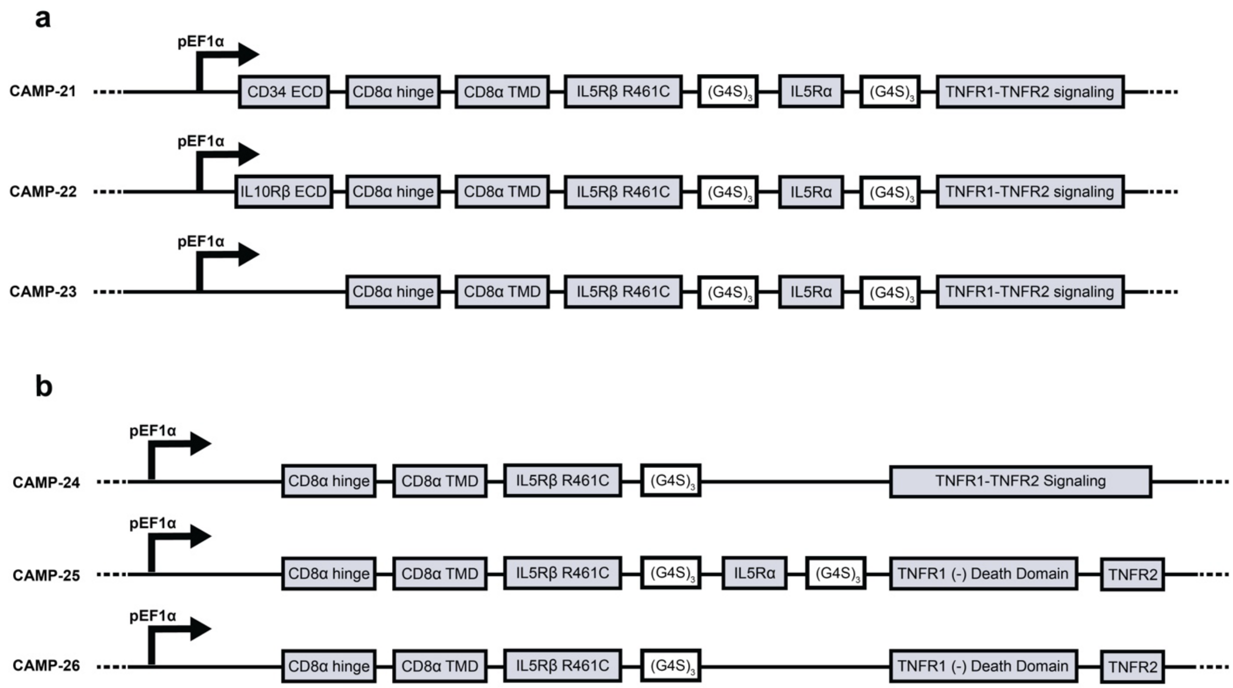


**Supplementary Figure 6. Revised CAMP construct interrogating the role of ECD choice on signaling.** **(a)** CAMP designs probing the role of ECDs on signaling magnitude (CAMP-21 through CAMP-23). Each is expressed from a pEF1α promoter and shares a CD8α hinge and TMD, followed by IL5Rβ R461C, IL5Rα, and a TNFR1–TNFR2 module joined by $(G4S)_3$ linkers, differing only in the ECD: CD34 ECD (CAMP-21), IL10Rβ ECD (CAMP-22), or no ECD (CAMP-23). **(b)** CAMP designs probing the role of intracellular domains (CAMP-24 through CAMP-26), which lack an extracellular domain and vary the intracellular signaling modules: CAMP-24 omits IL5Rα and retains a combined TNFR1–TNFR2 module, CAMP-25 includes a TNFR1 death domain deletion, and CAMP-26 omits both the IL5Rα module and the TNFR1 death domain. Boxes denote protein domains, and they are not drawn to scale. $(G4S)_3$: GGGGSGGGGSGGGGS linker, TNFR1 (-) Death Domain: TNFR1 with deletion of its death domain. Remaining abbreviations are as defined in **Supplementary Figures 1–4** (initial and revised CAMP constructs).

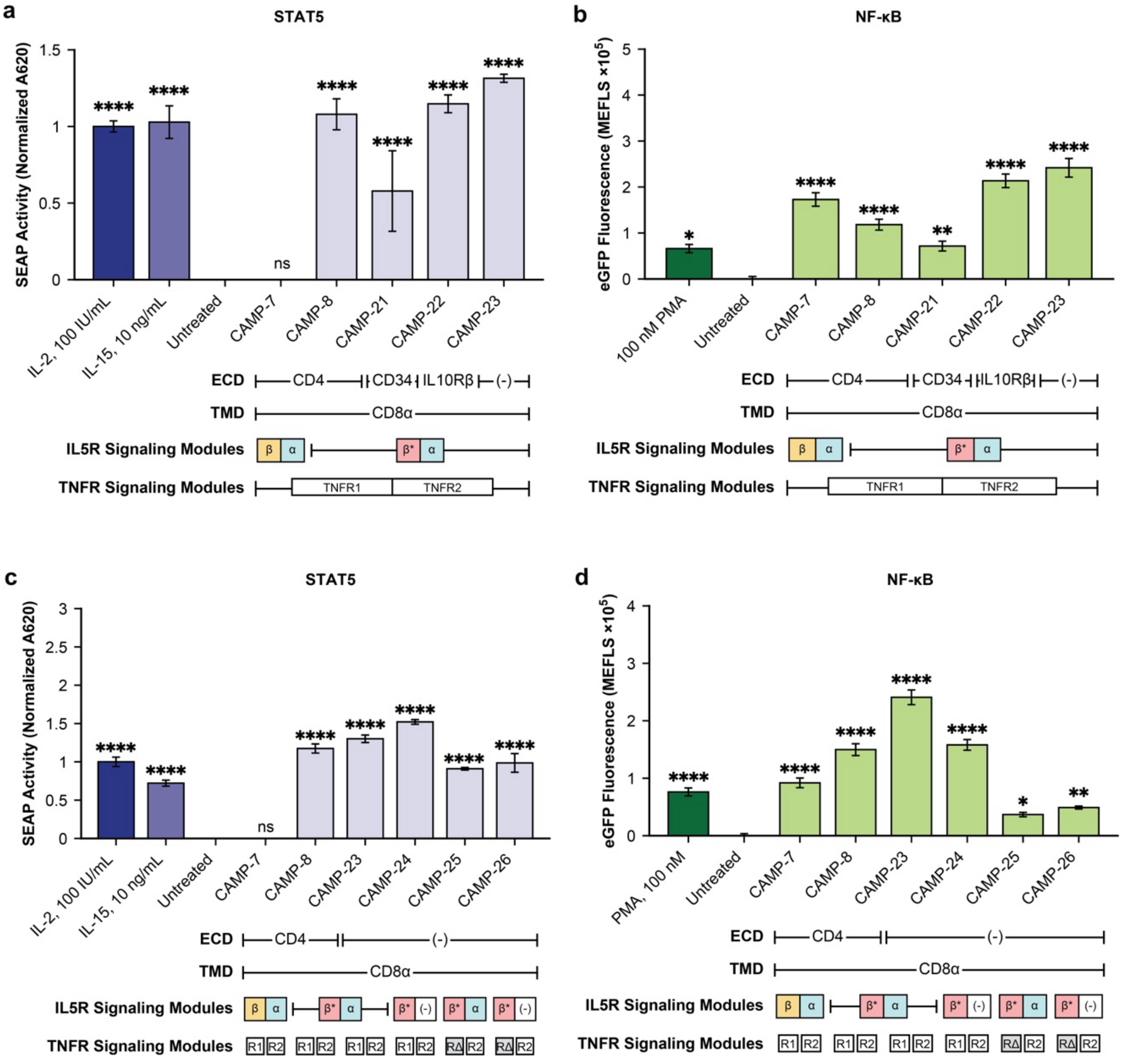


**Supplementary Figure 7. Repeat of experiments shown in Figure 3. (a,b)** Induction of STAT5 (a) and NF-κB.(b) signaling via CAMP designs probing the role of ECD on signaling magnitude. **(c,d)** Induction of STAT5 (c) and NF-κB (d) signaling via CAMP designs probing the role of intracellular domains. Bars show mean ± s.e.m. (n = 3); significance is versus the untreated condition by one-way ANOVA, followed by Dunnett's multiple comparisons test to evaluate specific comparisons (*p ≤ 0.05, **p ≤ 0.01, ***p ≤ 0.001, ****p ≤ 0.0001; ns, not significant), under the assumption of equal variance. ECD: extracellular domain; TMD: transmembrane domain. IL5R: interleukin-5 receptor, IL10Rβ: interleukin-10 receptor β, TNFR: tumor necrosis factor receptor, RΔ: TNFR1 death-domain deletion variant module, SEAP: secreted embryonic alkaline phosphatase (normalized A620), NF-κB: nuclear factor κB, PMA: phorbol 12-myristate 13-acetate, MEFL: molecules of equivalent fluorescein.

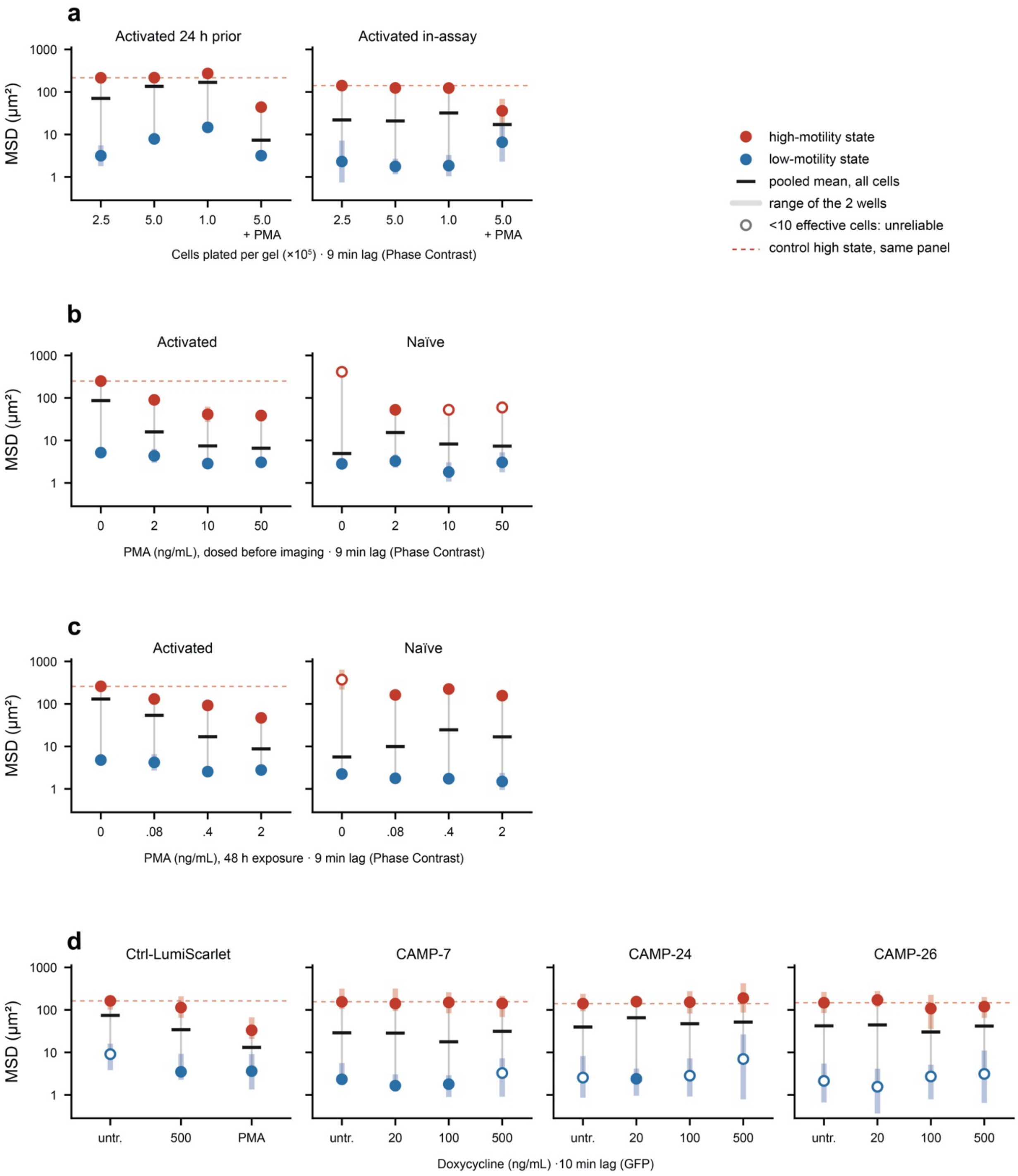


**Supplementary Figure 8. Single-cell motility states across cell density, PMA exposure, and doxycycline induction. (a)** Cell density and activation timing. Mean squared displacement (MSD) versus plating density for primary human T cells activated 24 h before plating or activated in-assay, each including a PMA-treated condition at one density. **(b)** Motility under acute PMA stimulation. PMA was added immediately prior to imaging (0 h), to either activated or naïve cells. **(c)** Motility under chronic PMA stimulation. Motility was evaluated after exposure to PMA for 48 prior to imaging (-48 h), using either activated or naïve cells. **(d)** Doxycycline titration of CAMP-engineered primary human T cells. n = 2 wells per condition for experiments (a–c) and n = 3 for experiment (n = 2 for the run 1 PMA condition) (d). Phase contrast (a–c; 9 -min MSD lag) and GFP (d; 10 min lag) data are not on a common absolute scale and should be compared only within, not across, these blocks. MSD: mean squared displacement; PMA: phorbol 12-myristate 13-acetate; GFP: green fluorescent protein.

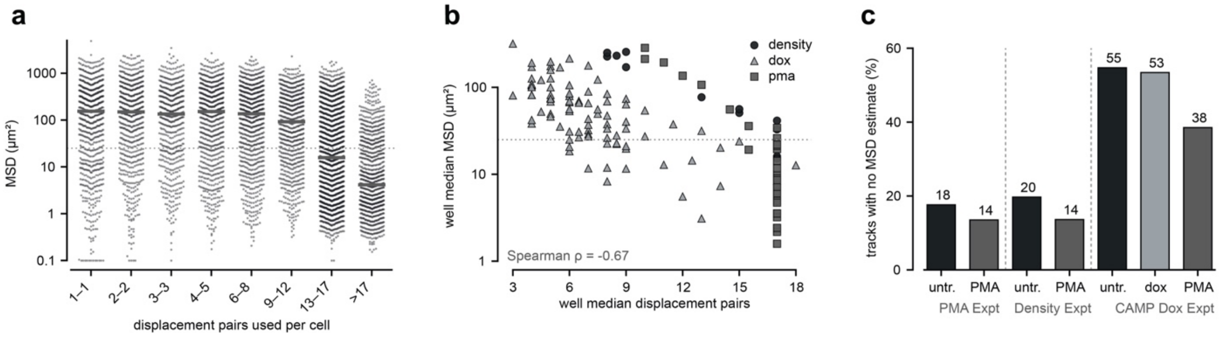


**Supplementary Figure 9. Track-length filtering biases per-cell motility estimates against the untreated condition.** Untreated wells both lose more cells to track-length filtering and retain shorter tracks among the survivors, and shorter tracks are scored as higher motility, so the surviving untreated population is biased toward higher motility. Treated-versus-untreated differences reported elsewhere may therefore be inflated, and they should be interpreted as upper bounds. To test whether this bias accounts for the reported treatment effects, we repeated three PMA comparisons using only well-tracked cells (those with ≥12 displacement pairs), a subset in which treated and untreated wells no longer differ in track length (reproduced in si_confound_checks.py in the code repository). Each comparison is the fold reduction in mean per-cell MSD, pooled across all cells of the condition, in PMA-treated versus untreated cells of the same arm. For pre-activated cells at 5 × $10^5$ cells per gel **(Figure 4b)**, the reduction is 11.3-fold among well-tracked cells versus 10.5-fold using all cells. For activated cells receiving PMA at 0 h, comparing 50 ng/mL with 0 ng/mL **(Figure 4c, upper left)**, it is 13.2-fold versus 10.7-fold; for activated cells receiving PMA at −48 h, comparing 2 ng/mL with 0 ng/mL **(Figure 4c, upper right)**, 8.0-fold versus 9.4-fold. The effects therefore persist at similar magnitude when the track-length difference is removed, so track-length bias does not account for them. **(a)** Per-cell MSD as a function of the number of displacement pairs contributing to it, binned; horizontal bars are within-bin medians. Cells supported by fewer displacement pairs return systematically higher MSD estimates. **(b)** Per-well median MSD versus per-well median displacement-pair count across 137 wells, colored by experimental arm (density, doxycycline, PMA). Wells with shorter retained tracks show higher median MSD (Spearman $\rho = -0.67$). **(c)** Fraction of tracked cells yielding no MSD estimate, by condition within each experimental arm. MSD: mean squared displacement; PMA: phorbol 12-myristate 13-acetate.

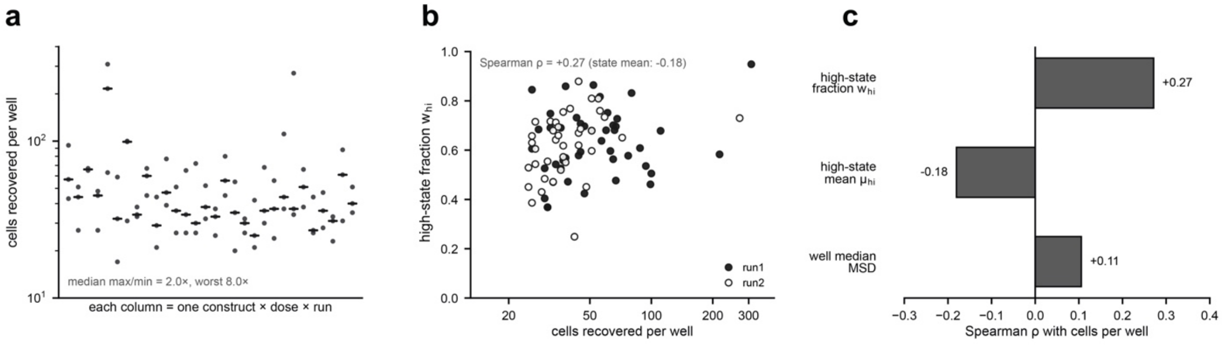


**Supplementary Figure 10. Cell recovery varies between replicate wells and tracks the high-motility state weight, not its position.** Cell recovery affects how many cells are assigned to the high-motility state but not the position of that state, so occupancy-based readouts require attention to recovery while state positions do not. One occupancy-based result in the main text is the rise in the fraction of naïve cells occupying the high-motility state under PMA treatment **(Figure 4c, lower row)**. Across the 16 naïve wells of that experiment, this fraction averages 0.14 in untreated wells (n = 4) and 0.44 in PMA-treated wells (n = 12), an unadjusted increase of +0.30. Adding cells recovered per well as a covariate in a linear model attributes +0.05 of that increase to recovery differences and leaves +0.25 attributable to PMA (p = 0.0004) (reproduced in si_confound_checks.py in the code repository). Recovery variation is therefore not of a scale that could explain the occupancy shift, though it accounts for a small part of it. **(a)** Cells recovered per well, with each column a single construct × dose × run combination and points the replicate wells within it; horizontal bars mark within-group medians. Replicate wells differ substantially in recovery (median max/min = 2.0×; worst case 8.0×). **(b)** Per-well high-motility state fraction ($w_{hi}$) versus cells recovered per well, split by run. Recovery correlates positively with the fraction of cells in the high-motility state (Spearman $\rho$ = +0.27) but negatively and more weakly with where that state sits (state mean; $\rho$ = −0.18). **(c)** Spearman correlation of each per-well readout with cells recovered per well, summarizing (b): the high-state fraction ($w_{hi}$, +0.27) tracks recovery, whereas the high-state mean ($\mu_{hi}$, −0.18) and well median MSD (+0.11) are comparatively insensitive to it. Since cell recovery moves the state weight but not the state position, state positions ($\mu_{hi}$) are the density-robust readout and weight-based readouts ($w_{hi}$) should not be interpreted without accounting for recovery. n = 89 wells. MSD: mean squared displacement; $w_{hi}$: fraction of cells in the high-motility state; $\mu_{hi}$: mean MSD of the high-motility state.

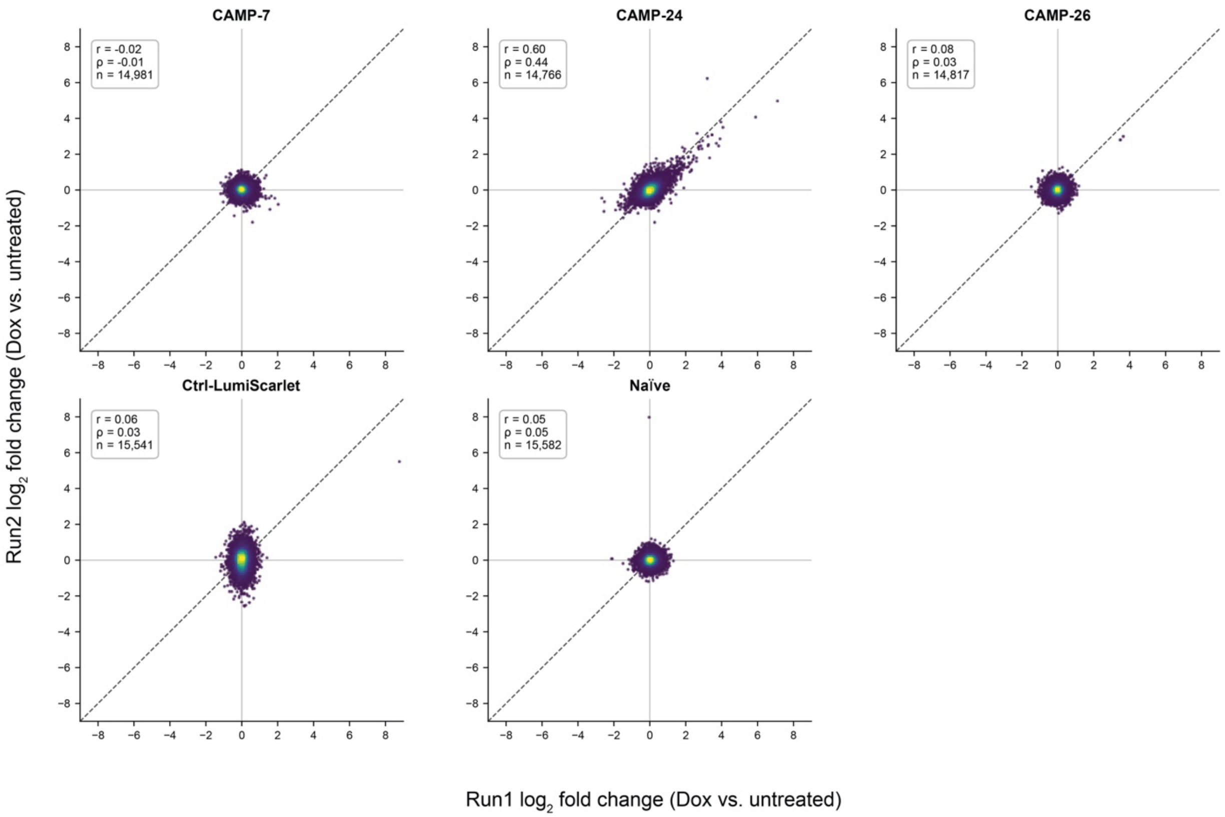


**Supplementary Figure 11. The doxycycline-induced transcriptional response is reproducible across independent runs for CAMP-24.** For each sample series, $\log_2$ fold change (doxycycline vs. untreated) was computed independently within each sequencing run from normalized expression (CPM, where genes with mean CPM ≥ 1 across the four samples in a sample series), and Run 1 is plotted against Run 2. Each point represents a gene, and the color indicates local point density. Perfect agreement (y=x) is indicated by the dotted line. Pearson (*r*), Spearman (*ρ*), and the number of genes plotted (*n*) are outlined in each panel. CAMP-24 shows a correlated transcript expression fold change response with doxycycline induction between both runs (r = 0.60), reflecting a reproducible response to dox-mediated induction of CAMP-24 expression. For the other samples (CAMP-7, CAMP-27, Ctrl-LumiScarlet, and Naïve cells), we observed a near-zero correlation (r ≤ 0.08), reflecting the absence of a reproducible transcriptional response.

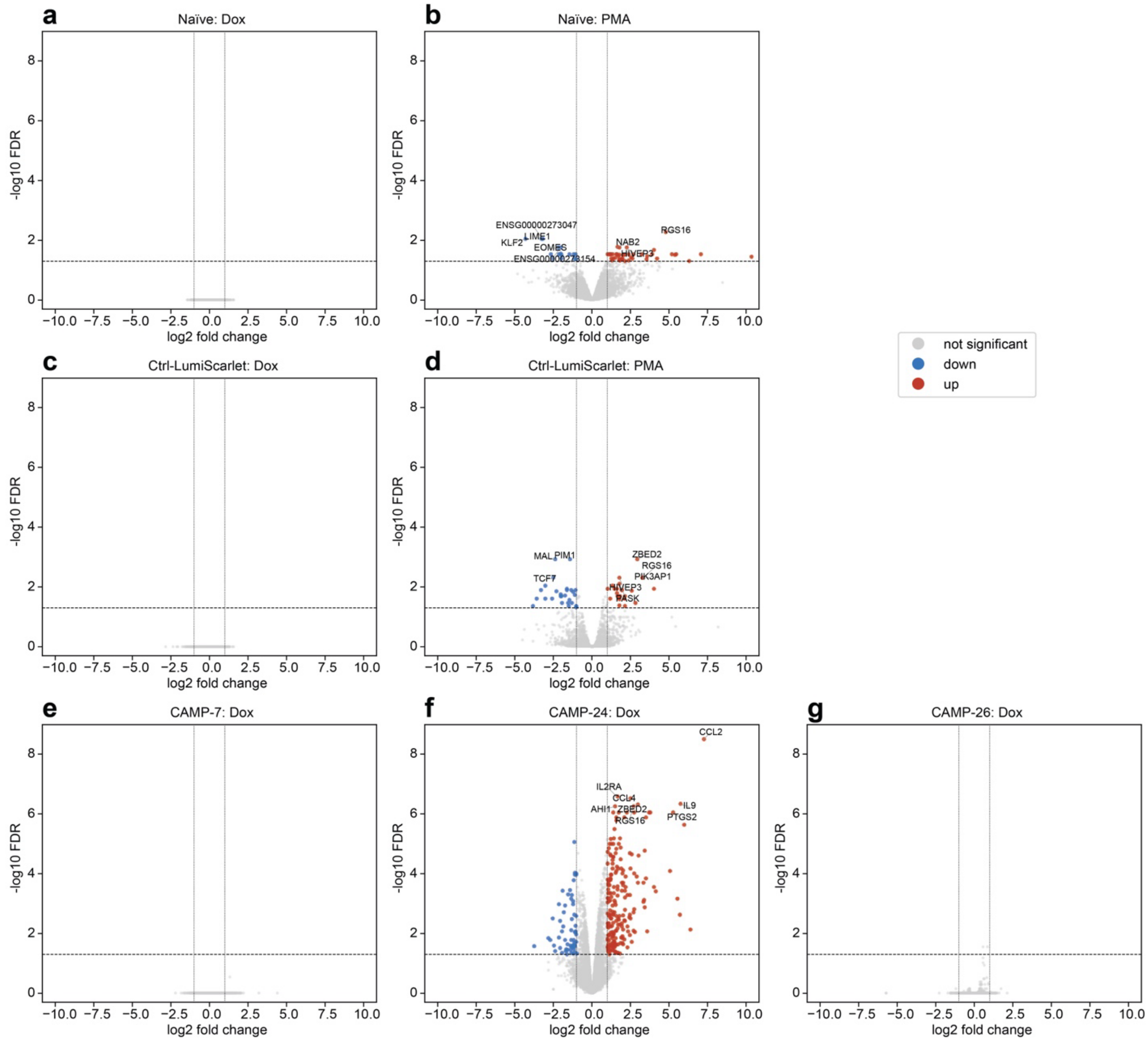


**Supplementary Figure 12. Differential gene expression across stimulus and construct.** Volcano plots of differential gene expression (treated vs. untreated) for each construct-stimulus combination: **(a)** Naïve + dox, **(b)** Naïve + PMA, **(c)** Ctrl-LumiScarlet + Dox, **(d)** Ctrl-LumiScarlet + PMA, **(e)** CAMP-7 + dox, **(f)** CAMP-24 + dox, and **(g)** CAMP-26 + dox. Each point is representative of one gene: red, significantly upregulated relative to untreated; blue, significantly downregulated relative to untreated (both FDR < 0.05 and $|log_2FC| > 1$); gray, no significant difference in expression. The dashed horizontal line marks FDR = 0.05, and the vertical dotted line represents $|log_2FC| = 1$; the x-axis communicates the $log_2$ fold change, and the y-axis $-log_{10}(FDR)$. Significant differential expression was detected in (b) Naïve + PMA (44 up, 16 down), (d) Ctrl-LumiScarlet + PMA (23 up, 28 down), and (f) CAMP-24 + dox (214 up, 60 down); for these conditions with significant differences in gene expression, the 8 most significant (lowest FDR) genes are labeled. Axes are shared across all figure panels. Each condition reflects a cell population of three pooled wells from the same condition, across two sequencing runs (n=2). Dox: 500 ng/mL doxycycline for 60 h; PMA: 0.4 ng/mL for 60 h.

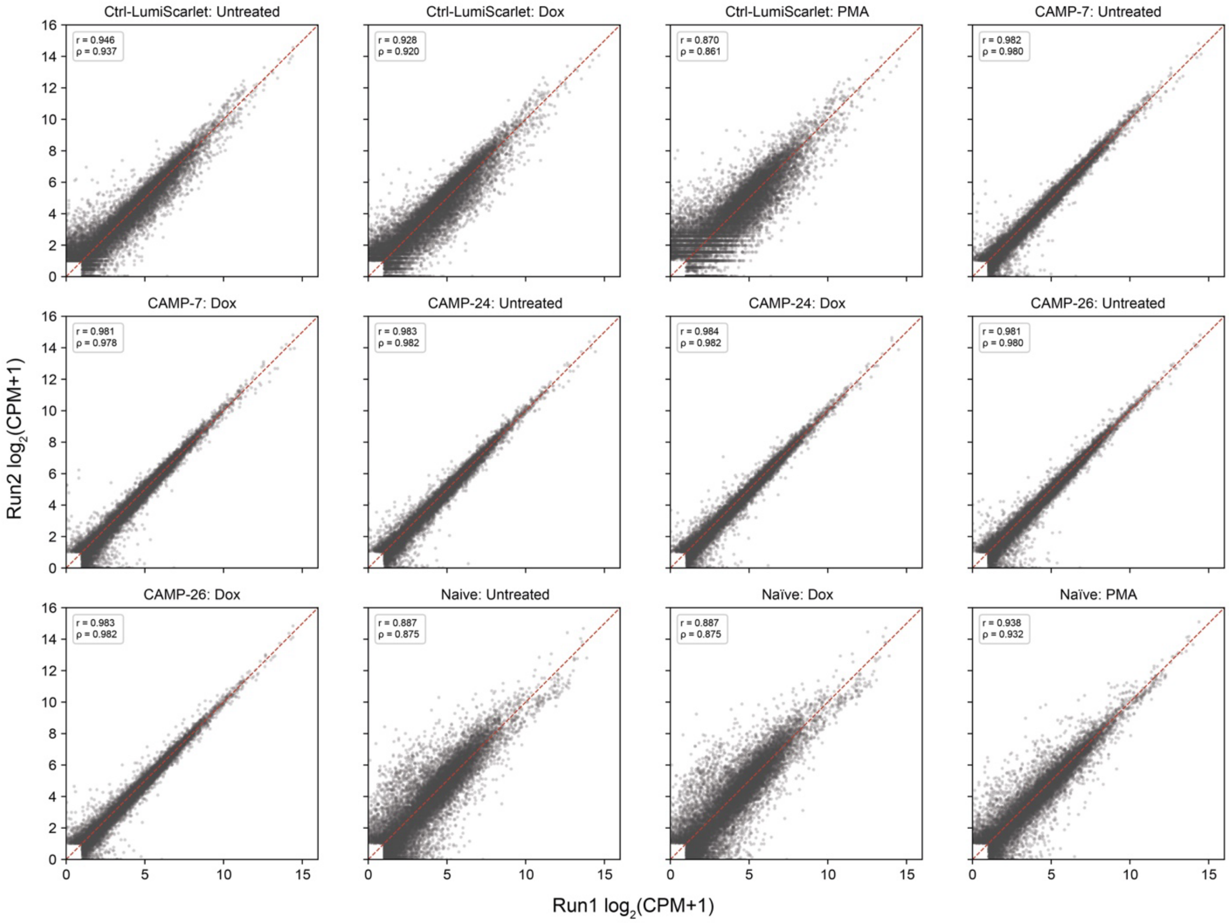


**Supplementary Figure 13. Run-to-run reproducibility of gene expression.** Comparison of per-gene expression between the two RNA-sequencing runs (Run 1 and Run 2) for each construct-treatment condition. In each panel, $\log_2$(CPM+1) in Run 1 (x-axis) is plotted against Run 2 (y-axis). Each point is one gene, restricted to genes with CPM > 1 in at least one run (n ≈ 15,400-17,200 genes per panel). Identity (y=x) is indicated by the red dashed line. Pearson (r) and Spearman (ρ) correlation coefficients are provided for each panel. Agreement was high across all conditions (median r = 0.963, ρ = 0.957; range r ≈ 0.87-0.98). CPM values were obtained from the batch-corrected output of the Plasmidsaurus pipeline, in which the two runs were aligned using the untreated CAMP-7, CAMP-24, and CAMP-26 samples as anchors. The correlations therefore reflect reproducibility after application of batch correction, and the higher agreement of the anchor samples relative to non-anchor naïve and control conditions is expected because the correction was estimated from these samples. Each batch reflects a cell population of three pooled wells from the same condition. Dox: 500 ng/mL doxycycline for 60 h; PMA: 0.4 ng/mL for 60 h.

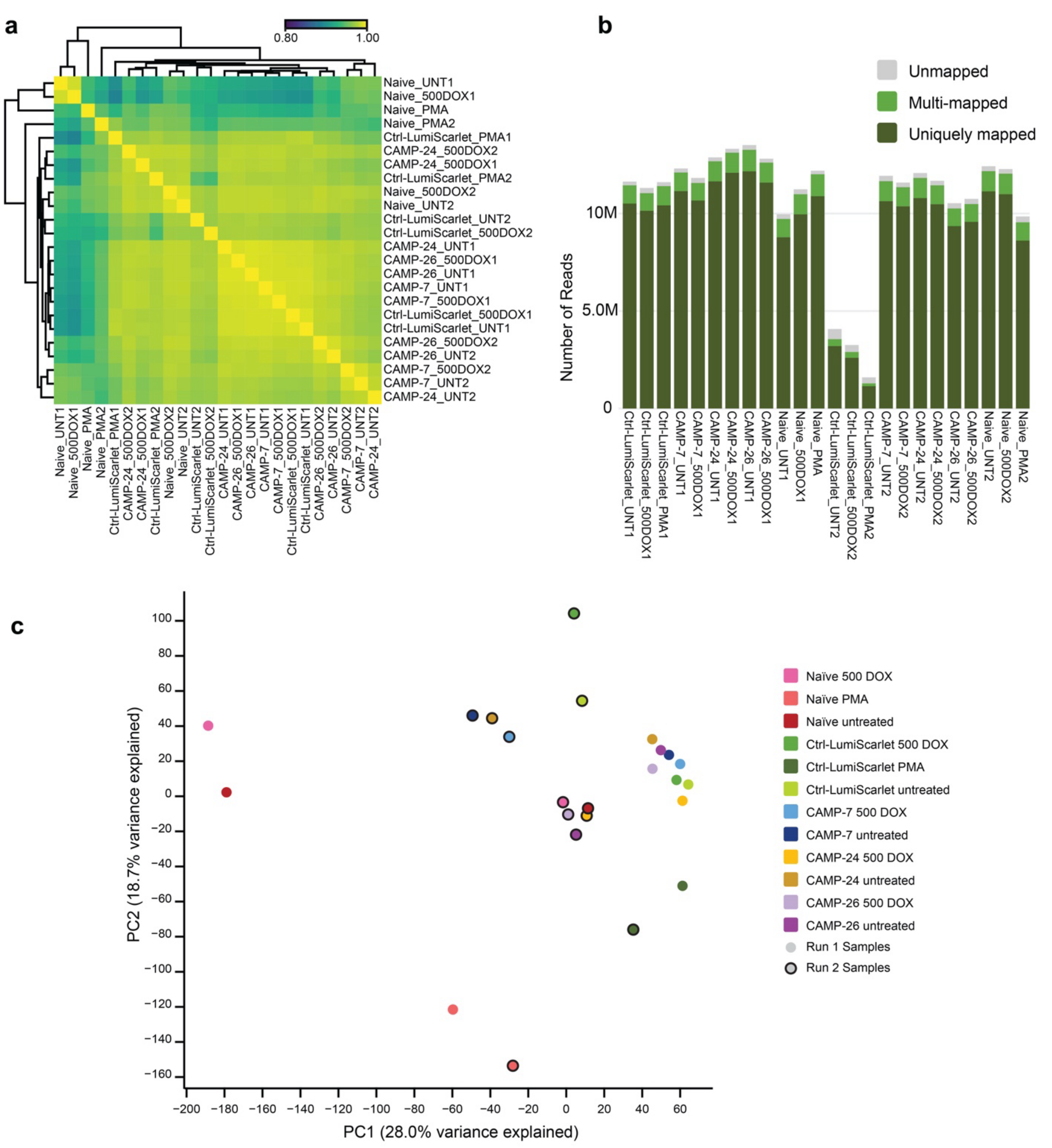


**Supplementary Figure 14. Q. Quality-control metrics for Plasmidsaurus RNA-seq libraries across both sample batches. (a)** Pairwise sample-to-sample correlation heatmap. Each cell provides the Pearson correlation between two libraries' genome-wide expression profiles, computed from TMM-normalized counts; the color scale spans 0.80 (dark) to 1.00 (yellow). Both axes are ordered by unsupervised hierarchical clustering (top and left dendrograms), grouping transcriptionally similar libraries together. Libraries from both RNA-seq batch runs are shown. **(b)** Per-library read-mapping. Stacked bars exhibit the number of deduplicated reads classified as uniquely mapped (dark green), multi-mapped (light green), or unmapped (gray) relative to GRCh38_114 reference genome. **(c)** PCA of the same RNA-seq libraries. Each point represents one sample projected onto the first two principal components, which capture 28.0% (PC1) and 18.7% (PC2) of the total expression variance. Fill colors denote the experimental condition; a black outline marks Run 2 samples vs. Run 1. Samples comprise naïve primary human T cells, or primary human T cells carrying either Ctrl-LumiScarlet or CAMP-7, CAMP-24, or CAMP-26 constructs, each profiled untreated (UNT), after doxycycline induction (500 DOX), or after PMA treatment. Each batch reflects a cell population of three pooled wells from the same condition. TMM: trimmed mean of m-values; PCA: principal component analysis; DOX: 500 ng/mL doxycycline for 60 h; PMA: phorbol 12-myristate 13-acetate 0.4 ng/mL for 60 h.

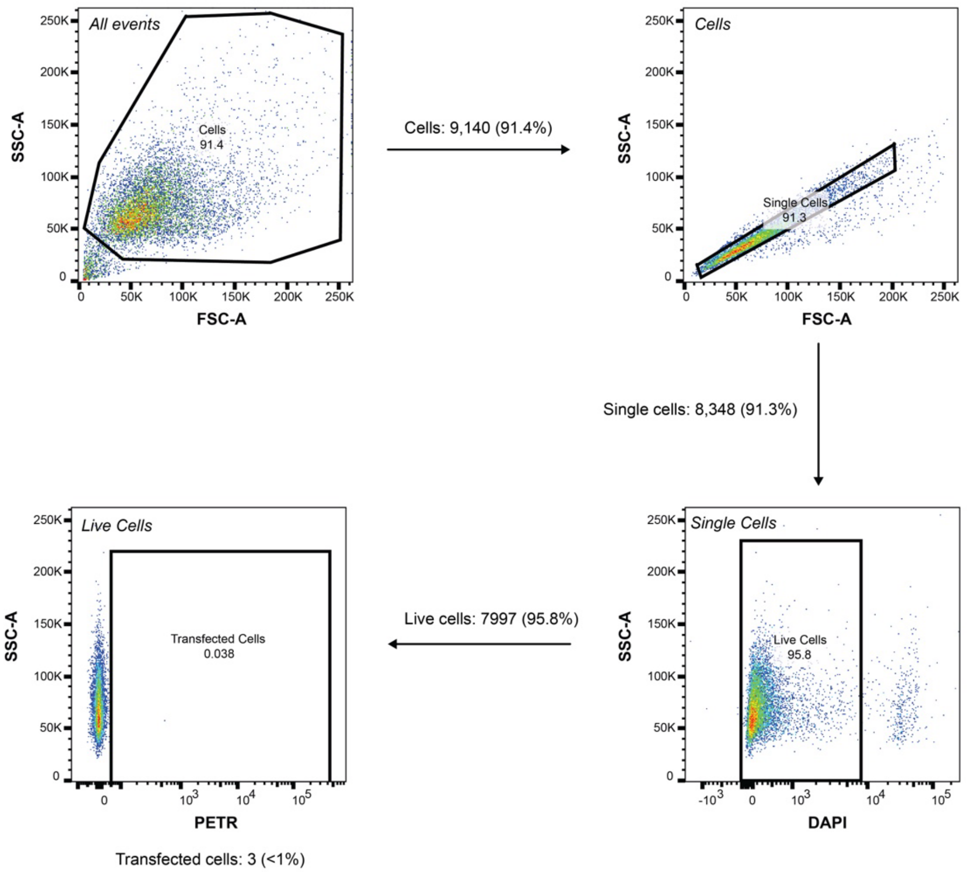


**Supplementary Figure 15. HEK cell flow cytometry gating scheme.** These plots show a sample of HEK293-NEE cells transfected with a modified pcDNA vector that does not contain any coding sequence downstream of the promoter. These cells were not transfected with a dsRedExpress2 (dsRE2) or mNeonGreen (mNG) expressing plasmid; thus they were used to define a negative gate for these fluorophores. In this gating protocol, HEK cells are identified using the SSC-A vs. FSC-A scatter profile. Single cells were identified from this population using the FSC-H vs. FSC-A scatter profile. Live cells were detected from single cells through identification of the DAPI$^-$ population within the SSC-A vs. DAPI profile. In other samples, the “transfected” population was defined as all single, live cells with a greater dsRE2 or mNG signal than the sample of single cells that was transfected with pcDNA only (shown). Note that both dsRE2 and mNG were used as transfection controls in some experiments. The gate defining the transfected population was drawn such that it did not include more than 1% of cells for this pcDNA-transfected negative control sample.

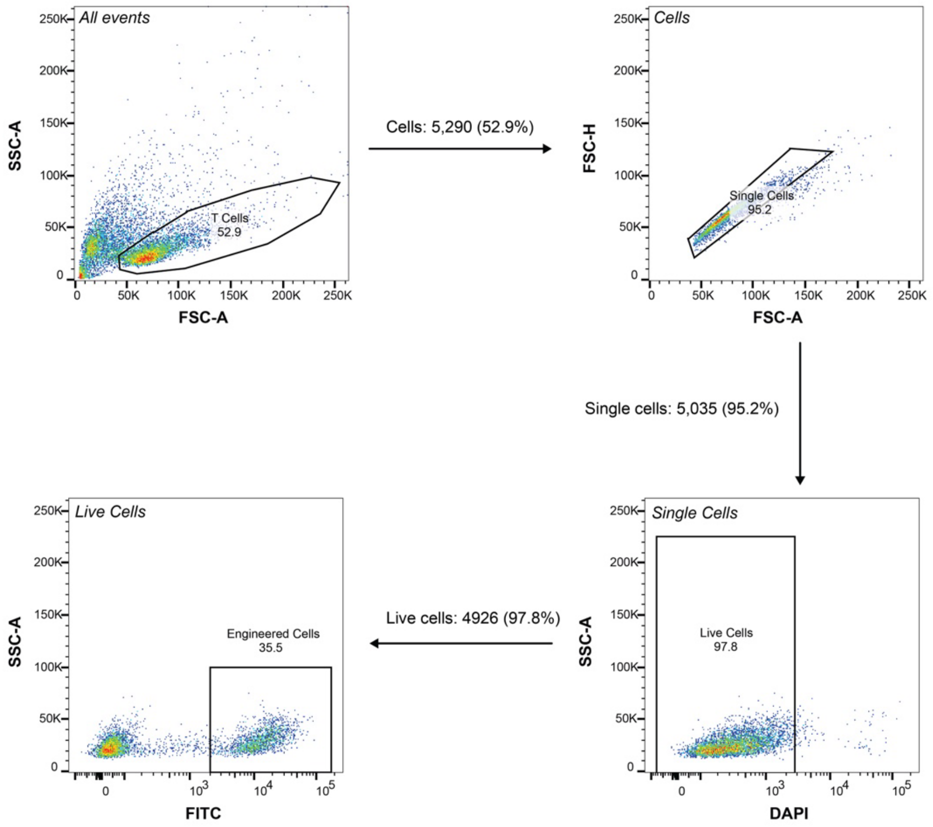


**Supplementary Figure 16. Primary T cell flow cytometry gating scheme.** These plots show a sample of engineered primary human T cells, which were previously electroporated with a PiggyBac transposon vector plasmid containing mNG and a pCMV-driven hyperactive PiggyBac transposase plasmid and expanded for 14 days under treatment with 30 ng/mL IL-4. In this gating scheme, primary human T cells are identified using the SSC-A vs. FSC-A scatter profile. Single cells were identified from this population using the FSC-H vs. FSC-A scatter profile. Live cells were detected from single cells through identification of the DAPI$^-$ population within the SSC-A vs. DAPI profile. The Engineered Cells gate was set based on the separation between fluorescent and non-fluorescent populations in this experimental control sample. The same gate was applied to all samples within each flow cytometry analysis to consistently define engineered cells.

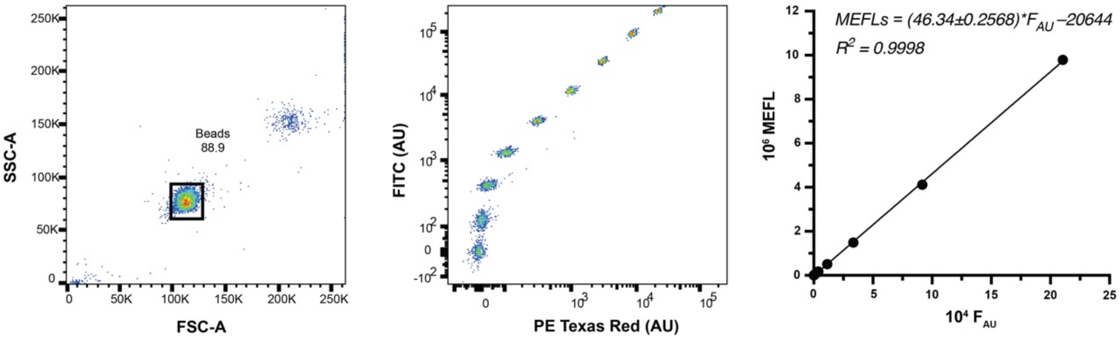


**Supplementary Figure 17. Profile of fluorescent calibration beads.** Spherotech Ultra Rainbow Calibration Particles (URCP) used in this study have nine fluorescent bead populations. Beads are first identified using the FSC-A vs. SSC-A channel profile, and beads are fluorescent in the majority of cytometer channels. In each experiment, identification of individual bead populations was achieved using two channels. The mean intensity of each bead population in the eGFP (FITC) channel was recorded in arbitrary units ($F_{AU}$) and plotted against the number of fluorophores present on the beads described by the manufacturer. A simple linear regression was performed to generate the calibration curve. MFI were converted to MEFLs in each experiment using the $F_{AU}$ multiplier described by the regression; the magnitude and corresponding uncertainty of each multiplier is reported in parentheses on the plot. This plot shows an example of one such calibration curve, one of which was obtained for each such flow cytometry run.

### SUPPLEMENTARY TABLES AND NOTES

**Supplemental Note 1. Nomenclature and modular composition of CAMP chimeric receptors.** Here we summarize the nomenclature used throughout the manuscript and summarized in **Supplementary Table 1**. Each construct carries a short identifier (used in the main text) and a systematic descriptor describing its architecture. Descriptors are read from starting from structures external to the membrane: extracellular domain (ECD)—transmembrane domain (TMD)—intracellular signaling modules (listed proximal to distal). Definitions are given in the key at the top of the table. Constructs were generated in sets throughout the study. The first set of CAMPs compares transmembrane domains on a shared CD4-ECD/IL5Rα–IL5Rβ/TNFR1–TNFR2 backbone, with direct fusions between domains. The second set of CAMPs comprises a combinatorial screen that changes the ECD/TMD module (CD4 ECD paired with CD8α-family TMDs; CD34 ECD paired with the constitutively active IL7R "C7R" TMD), retention of a juxtamembrane segment (+J), the order of the IL5Rα and IL5Rβ subunits, and wild-type versus constitutively active IL5Rβ (R461C). The third set of CAMPs swaps or removes the ECD on receptors validated for function. The fourth set of CAMPs explores minimizing the receptor by removing IL5Rα and/or the TNFR1 death domain. Symbols: "~" denotes a $(G4S)_3$ flexible linker at the two IL5R junctions (present from Generation 2 onward) and "-" denotes a direct fusion. β* = IL5Rβ bearing activating R461C substitution; β = wild-type IL5Rβ (CSF2RB/βc); α = IL5Rα; ΔE = ECD-less (CD4 signal peptide plus CD8α hinge only); +J = juxtamembrane domain retained; T1 and T1ΔDD = TNFR1 with and without its death domain (DD); T2 = TNFR2. The final column lists the figure(s) in which each construct appears. Ctrl-LumiScarlet is a doxycycline-inducible fluorescent reporter used for enrichment and as an assay control, not a receptor.

**Supplementary Table 1. CAMP nomenclature.**

| Field | Components |
|---|---|
| **ECD** | CD4, CD34, IL10RB; ΔE = no ECD (CD4 signal peptide + CD8α hinge only) |
| **TMD** | CD8 (CD8α), CD4, CD28, IL7R* (constitutively active mutant IL7R); +J suffix = juxtamembrane domain retained |
| **Intracellular (proximal to distal)** | β* = IL5RB(R461C, active); β = IL5RB WT [CSF2RB/βc]; α = IL5RA; T1 = TNFR1 (intact death domain); T1ΔDD = TNFR1 lacking death domain; T2 = TNFR2. |
| **Linker** | ~ = $(G4S)_3$ flexible linker (G4S = GGGGS). For constructs with two IL5R domains, a "~" marks a $(G4S)_3$ linker at the two IL5R junctions: between the IL5Rβ and IL5Rα subunits, and between the distal IL5R subunit and TNFR1. A plain hyphen (-) is a direct fusion; all other junctions (including TNFR1–TNFR2) are direct. |

| Short name | Systematic descriptor | Distinguishing feature | Figures |
|---|---|---|---|
| CAMP-1 | CD4-CD8-α-β-T1-T2 | CD8α TMD | 1 |
| CAMP-2 | CD4-CD4-α-β-T1-T2 | CD4 TMD | 1 |
| CAMP-3 | CD4-CD28-α-β-T1-T2 | CD28 TMD | 1 |
| CAMP-4 | CD34-IL7R*+J-α-β-T1-T2 | C7R-style disulfide TMD | 1 |
| CAMP-5 | CD4-CD8-α~β~T1-T2 | CD8α; IL5Rα-proximal; WT IL5Rβ | 2 |
| CAMP-6 | CD4-CD8-α~β*~T1-T2 | CD8α; IL5Rα-proximal; active IL5Rβ (R461C) | 2 |
| CAMP-7 | CD4-CD8-β~α~T1-T2 | CD8α; IL5Rβ-proximal; WT IL5Rβ | 2 |
| CAMP-8 | CD4-CD8-β*~α~T1-T2 | First STAT5-competent architecture; parent of F3 | 2 |
| CAMP-9 | CD4-CD8+J-α~β~T1-T2 | CD8α +JM; IL5Rα-proximal; WT IL5Rβ | 2 |
| CAMP-10 | CD4-CD8+J-α~β*~T1-T2 | CD8α +JM; IL5Rα-proximal; active IL5Rβ (R461C) | 2 |

| | | | |
|---|---|---|---|
| CAMP-11 | CD4-CD8+J-β~α~T1-T2 | CD8α +JM; IL5Rβ-proximal; WT IL5Rβ | 2 |
| CAMP-12 | CD4-CD8+J-β*~α~T1-T2 | CD8α +JM; IL5Rβ-proximal; active IL5Rβ (R461C) | 2 |
| CAMP-13 | CD34-IL7R*-α~β~T1-T2 | C7R (act. IL7R); IL5Rα-proximal; WT IL5Rβ | 2 |
| CAMP-14 | CD34-IL7R*-α~β*~T1-T2 | C7R (act. IL7R); IL5Rα-proximal; active IL5Rβ (R461C) | 2 |
| CAMP-15 | CD34-IL7R*-β~α~T1-T2 | C7R (act. IL7R); IL5Rβ-proximal; WT IL5Rβ | 2 |
| CAMP-16 | CD34-IL7R*-β*~α~T1-T2 | C7R (act. IL7R); IL5Rβ-proximal; active IL5Rβ (R461C) | 2 |
| CAMP-17 | CD34-IL7R*+J-α~β~T1-T2 | C7R (act. IL7R) +JM; IL5Rα-proximal; WT IL5Rβ | 2 |
| CAMP-18 | CD34-IL7R*+J-α~β*~T1-T2 | C7R (act. IL7R) +JM; IL5Rα-proximal; active IL5Rβ (R461C) | 2 |
| CAMP-19 | CD34-IL7R*+J-β~α~T1-T2 | C7R (act. IL7R) +JM; IL5Rβ-proximal; WT IL5Rβ | 2 |
| CAMP-20 | CD34-IL7R*+J-β*~α~T1-T2 | C7R (act. IL7R) +JM; IL5Rβ-proximal; active IL5Rβ (R461C) | 2 |
| CAMP-21 | CD34-CD8-β*~α~T1-T2 | ECD swap (CD34) | 3 |
| CAMP-22 | IL10RB-CD8-β*~α~T1-T2 | ECD swap (IL10RB; lowest output) | 3 |
| CAMP-23 | ΔE-CD8-β*~α~T1-T2 | Minimal (ECD-less); parent of Gen 4 deletion series | 3 |
| CAMP-24 | ΔE-CD8-β*~T1-T2 | − IL5RA | 3, 6 |
| CAMP-25 | ΔE-CD8-β*~α~T1ΔDD-T2 | − TNFR1 death domain | 3, 6 |
| CAMP-26 | ΔE-CD8-β*~T1ΔDD-T2 | − IL5RA and − DD | 3, 6 |
| Ctrl-LumiScarlet | reporter, not a receptor | Dox-inducible reporter; enrichment/assay control | 6 |

**Supplemental Note 2. Statistics.** Here we list additional statistical calculation details by figure.

Below are the outcomes of Dunnett's multiple comparisons tests performed following ordinary one-way ANOVA. In every panel each condition was compared against the untreated control within a single family of comparisons at α = 0.05, and all p values reported below are multiplicity-adjusted (Dunnett-adjusted) values. Null hypotheses were that there existed no effect of the indicated treatment on the measured reporter expression. Multiplicity-adjusted p values are reported.

**Reporter assay in Figure 1c**

(Condition: adjusted p value for comparison untreated vs. indicated condition; 11 comparisons in one family, n = 3 per group, df = 24, α = 0.05)

- CAMP-4: >0.9999 (not significant, n.s.)
- CAMP-2: >0.9999 (n.s.)
- CAMP-3: >0.9999 (n.s.)
- CAMP-1: >0.9999 (n.s.)
- IL-2, 100 IU/mL: <0.0001
- IL-15, 10 ng/mL: <0.0001

**Reporter assay in Figure 1d**

(Condition: adjusted p value for comparison untreated vs. indicated condition; 7 comparisons in one family, n = 3 per group, df = 16, α = 0.05)

- CAMP-4: <0.0001
- CAMP-2: <0.0001
- CAMP-1: <0.0001
- CAMP-3: <0.0001
- 100 nM PMA: 0.0003

**Reporter assay in Figure 2c**

(Condition: adjusted p value for comparison untreated vs. indicated condition; 23 comparisons in one family, n = 3 per group, df = 48, α = 0.05)

- CAMP-5: >0.9999 (not significant, n.s.)
- CAMP-6: >0.9999 (n.s.)
- CAMP-7: >0.9999 (n.s.)
- CAMP-8: <0.0001
- CAMP-9: >0.9999 (n.s.)
- CAMP-10: >0.9999 (n.s.)
- CAMP-11: >0.9999 (n.s.)
- CAMP-12: >0.9999 (n.s.)
- CAMP-13: >0.9999 (n.s.)
- CAMP-14: >0.9999 (n.s.)
- CAMP-15: >0.9999 (n.s.)
- CAMP-16: >0.9999 (n.s.)
- CAMP-17: >0.9999 (n.s.)
- CAMP-18: >0.9999 (n.s.)
- CAMP-19: >0.9999 (n.s.)
- CAMP-20: >0.9999 (n.s.)
- IL-2, 100 IU/mL: <0.0001
- IL-15, 10 ng/mL: <0.0001

**Reporter assay in Figure 2d**

(Condition: adjusted p value for comparison untreated vs. indicated condition; 19 comparisons in one family, n = 3 per group, df = 40, α = 0.05)

- CAMP-5: <0.0001
- CAMP-6: <0.0001

- CAMP-7: <0.0001
- CAMP-8: <0.0001
- CAMP-9: <0.0001
- CAMP-10: <0.0001
- CAMP-11: <0.0001
- CAMP-12: <0.0001
- CAMP-13: >0.9999 (not significant, n.s.)
- CAMP-14: >0.9999 (n.s.)
- CAMP-15: >0.9999 (n.s.)
- CAMP-16: >0.9999 (n.s.)
- CAMP-17: <0.0001
- CAMP-18: <0.0001
- CAMP-19: <0.0001
- CAMP-20: <0.0001
- 100 nM PMA: <0.0001

**Reporter assay in Figure 3a**
(Condition: adjusted p value for comparison untreated vs. indicated condition; 14 comparisons in one family, n = 3 per group, df = 30, α = 0.05)

- CAMP-7: >0.9999 (not significant, n.s.)
- CAMP-8: <0.0001
- CAMP-22: <0.0001
- CAMP-21: 0.0494
- CAMP-23: <0.0001
- IL-2, 100 IU/mL: <0.0001
- IL-15, 10 ng/mL: <0.0001

**Reporter assay in Figure 3b**
(Condition: adjusted p value for comparison untreated vs. indicated condition; 7 comparisons in one family, n = 3 per group, df = 16, α = 0.05)

- CAMP-7: <0.0001
- CAMP-8: <0.0001
- CAMP-22: <0.0001
- CAMP-21: 0.0058
- CAMP-23: <0.0001
- 100 nM PMA: 0.0107

**Reporter assay in Figure 3c**
(Condition: adjusted p value for comparison untreated vs. indicated condition; 15 comparisons in one family, n = 3 per group, df = 32, α = 0.05)

- CAMP-7: >0.9999 (not significant, n.s.)
- CAMP-8: <0.0001
- CAMP-23: <0.0001
- CAMP-24: <0.0001
- CAMP-25: <0.0001
- CAMP-26: <0.0001
- IL-2, 100 IU/mL: <0.0001
- IL-15, 10 ng/mL: <0.0001

**Reporter assay in Figure 3d**
(Condition: adjusted p value for comparison untreated vs. indicated condition; 8 comparisons in one family, n = 3 per group, df = 18, α = 0.05)

- CAMP-7: <0.0001
- CAMP-8: <0.0001
- CAMP-23: <0.0001

- CAMP-24: <0.0001
- CAMP-25: 0.0461
- CAMP-26: 0.0058
- PMA, 100 nM: <0.0001

Below are the outcomes of pooled-error ordinary least squares (OLS) tests on the log10 of the per-well arithmetic mean MSD signal, in which a single within-condition variance was estimated per panel, followed by the Benjamini-Hochberg (BH) procedure. Null hypotheses were that there existed no effect of the indicated treatment on the measured signal. These tests supersede the two-sample Welch's t-tests on the same comparisons, which are reported alongside each result for reference.

**MSD assay in Figure 4b**

(Timing of PMA addition: BH-adjusted p value for comparison $5×10^5$ cells untreated vs. $5×10^5$ cells + PMA; 2 wells per condition, pooled residual df = 4)

- 24 h prior: 0.000310
  - Ratio of group means = 0.09710 (equivalently 10.30-fold); pooled SD of log10 signal = 0.07280; uncorrected p = 0.000155; Welch's t-test on the same comparison (superseded), p = 0.0192 at df = 1.190.
- In-assay: 0.0163
  - Ratio of group means = 0.5092 (equivalently 1.960-fold); pooled SD of log10 signal = 0.07350; uncorrected p = 0.0163; Welch's t-test on the same comparison (superseded), p = 0.202 at df = 1.030.

**MSD assay in Figure 4c**

(Timing of PMA addition, dose: BH-adjusted p value for comparisons vs. the 0 ng/mL condition of the matching panel; 8 wells per panel, pooled residual df = 4)

- Activated, PMA added at 0 h, 0 vs. 2 ng/mL: 0.00841
  - Ratio of group means = 0.2381; t = −4.839 at df = 4; pooled SD of log10 signal = 0.1288; uncorrected p = 0.00841; Welch's t-test on the same comparison (superseded), p = 0.0173.
- Activated, PMA added at 0 h, 0 vs. 10 ng/mL: 0.00262
  - Ratio of group means = 0.1103; t = −7.434 at df = 4; pooled SD of log10 signal = 0.1288; uncorrected p = 0.00175; Welch's t-test on the same comparison (superseded), p = 0.0241.
- Activated, PMA added at 0 h, 0 vs. 50 ng/mL: 0.00254
  - Ratio of group means = 0.09090; t = −8.086 at df = 4; pooled SD of log10 signal = 0.1288; uncorrected p = 0.00127; Welch's t-test on the same comparison (superseded), p = 0.0508.
- Activated, PMA added at −48 h, 0 vs. 0.08 ng/mL: 0.00841
  - Ratio of group means = 0.5001; t = −4.954 at df = 4; pooled SD of log10 signal = 0.06074; uncorrected p = 0.00774; Welch's t-test on the same comparison (superseded), p = 0.0975.
- Activated, PMA added at −48 h, 0 vs. 0.4 ng/mL: 0.00204
  - Ratio of group means = 0.2642; t = −9.519 at df = 4; pooled SD of log10 signal = 0.06074; uncorrected p = 0.000680; Welch's t-test on the same comparison (superseded), p = 0.0137.
- Activated, PMA added at −48 h, 0 vs. 2 ng/mL: 0.000514
  - Ratio of group means = 0.1043; t = −16.16 at df = 4; pooled SD of log10 signal = 0.06074; uncorrected p = $8.57×10^{-5}$; Welch's t-test on the same comparison (superseded), p = 0.00554.

Below are the outcomes of two-tailed Welch's t-tests on the log10 of the per-well arithmetic mean MSD signal. Null hypotheses were that there existed no effect of the indicated treatment on the measured signal. These comparisons were not included in the pooled-error families above and are reported without multiplicity adjustment.

**MSD assay in Figure 4c, sequential dose comparisons**

(Timing of PMA addition, dose step: p value for comparison between successive doses; 2 wells per condition)

- Activated, PMA added at 0 h, 0 vs. 2 ng/mL: 0.0173
  - Ratio of group means = 0.2381; t = −7.672 at df = 1.971.

- Activated, PMA added at 0 h, 2 vs. 10 ng/mL: 0.115 (not significant, n.s.)
  - Ratio of group means = 0.4632; t = −2.990 at df = 1.714.
- Activated, PMA added at 0 h, 10 vs. 50 ng/mL: 0.663 (n.s.)
  - Ratio of group means = 0.8242; t = −0.5151 at df = 1.795.
- Activated, PMA added at −48 h, 0 vs. 0.08 ng/mL: 0.0975 (n.s.)
  - Ratio of group means = 0.5001; t = −5.929 at df = 1.055.
- Activated, PMA added at −48 h, 0.08 vs. 0.4 ng/mL: 0.0925 (n.s.)
  - Ratio of group means = 0.5282; t = −6.106 at df = 1.069.
- Activated, PMA added at −48 h, 0.4 vs. 2 ng/mL: 0.0301
  - Ratio of group means = 0.3948; t = −5.825 at df = 1.944.

**MSD assay in Figure 6b**

(Cell line, independent run: p value for comparison untreated vs. indicated dose or PMA treatment: dox doses administered at 20 ng/mL, 100 ng/mL, and 500 ng/mL; PMA doses administered at 0.4 ng/mL n = 2–3 wells per condition)

- Ctrl-LumiScarlet, run 1, untreated vs. 500: 0.545 (not significant, n.s.)
- Ctrl-LumiScarlet, run 1, untreated vs. PMA: 0.0621 (n.s.)
- Ctrl-LumiScarlet, run 2, untreated vs. 500: 0.190 (n.s.)
- Ctrl-LumiScarlet, run 2, untreated vs. PMA: 0.0249
- CAMP-7, run 1, untreated vs. 20: 0.691 (n.s.)
- CAMP-7, run 1, untreated vs. 100: 0.407 (n.s.)
- CAMP-7, run 1, untreated vs. 500: 0.560 (n.s.)
- CAMP-7, run 2, untreated vs. 20: 0.185 (n.s.)
- CAMP-7, run 2, untreated vs. 100: 0.696 (n.s.)
- CAMP-7, run 2, untreated vs. 500: 0.892 (n.s.)
- CAMP-24, run 1, untreated vs. 20: 0.112 (n.s.)
- CAMP-24, run 1, untreated vs. 100: 0.166 (n.s.)
- CAMP-24, run 1, untreated vs. 500: 0.582 (n.s.)
- CAMP-24, run 2, untreated vs. 20: 0.982 (n.s.)
- CAMP-24, run 2, untreated vs. 100: 0.442 (n.s.)
- CAMP-24, run 2, untreated vs. 500: 0.925 (n.s.)
- CAMP-26, run 1, untreated vs. 20: 0.313 (n.s.)
- CAMP-26, run 1, untreated vs. 100: 0.835 (n.s.)
- CAMP-26, run 1, untreated vs. 500: 0.915 (n.s.)
- CAMP-26, run 2, untreated vs. 20: 0.951 (n.s.)
- CAMP-26, run 2, untreated vs. 100: 0.196 (n.s.)
- CAMP-26, run 2, untreated vs. 500: 0.374 (n.s.)

Below is the outcome of an ordinary least squares (OLS) test on the log10 of the per-well arithmetic mean MSD signal with independent run included as a blocking factor. The null hypothesis was that there existed no effect of PMA treatment on the measured signal once run-to-run variation was accounted for.

**MSD assay in Figure 6b, runs pooled**

(Cell line: p value for comparison untreated vs. PMA treatment across both runs)

- Ctrl-LumiScarlet, untreated vs. PMA: 0.000102
  - Ratio of group means = 0.1730; n = 6 untreated wells and 5 PMA-treated wells pooled across runs.

Below are the outcomes of Dunnett's multiple comparisons tests performed following ordinary one-way ANOVA. In every panel each condition was compared against the untreated control within a single family of comparisons at $\alpha = 0.05$, and all p values reported below are multiplicity-adjusted (Dunnett-adjusted) values. Null hypotheses were that there existed no effect of the indicated treatment on the measured reporter expression. Multiplicity-adjusted p values are reported.

**Reporter assay in Supplementary Figure 2a**

(Condition: adjusted p value for comparison untreated vs. indicated condition; 11 comparisons in one family, n = 3 per group, df = 24, α = 0.05)

- CAMP-4: >0.9999 (not significant, n.s.)
- CAMP-2: >0.9999 (n.s.)
- CAMP-3: >0.9999 (n.s.)
- CAMP-1: >0.9999 (n.s.)
- IL-2, 100 IU/mL: <0.0001
- IL-15, 10 ng/mL: <0.0001

**Reporter assay in Supplementary Figure 2b**

(Condition: adjusted p value for comparison untreated vs. indicated condition; 7 comparisons in one family, n = 3 per group, df = 16, α = 0.05)

- CAMP-4: <0.0001
- CAMP-2: <0.0001
- CAMP-1: <0.0001
- CAMP-3: <0.0001
- 100 nM PMA: <0.0001

**Reporter assay in Supplementary Figure 4a**

(Condition: adjusted p value for comparison untreated vs. indicated condition; 23 comparisons in one family, n = 3 per group, df = 48, α = 0.05)

- CAMP-5: >0.9999 (not significant, n.s.)
- CAMP-6: >0.9999 (n.s.)
- CAMP-7: >0.9999 (n.s.)
- CAMP-8: <0.0001
- CAMP-9: >0.9999 (n.s.)
- CAMP-10: >0.9999 (n.s.)
- CAMP-11: >0.9999 (n.s.)
- CAMP-12: 0.4584 (n.s.)
- CAMP-13: >0.9999 (n.s.)
- CAMP-14: >0.9999 (n.s.)
- CAMP-15: >0.9999 (n.s.)
- CAMP-16: >0.9999 (n.s.)
- CAMP-17: >0.9999 (n.s.)
- CAMP-18: >0.9999 (n.s.)
- CAMP-19: >0.9999 (n.s.)
- CAMP-20: >0.9999 (n.s.)
- IL-2, 100 IU/mL: <0.0001
- IL-15, 10 ng/mL: <0.0001

**Reporter assay in Supplementary Figure 4b**

(Condition: adjusted p value for comparison untreated vs. indicated condition; 19 comparisons in one family, n = 3 per group, df = 40, α = 0.05)

- CAMP-5: <0.0001
- CAMP-6: <0.0001
- CAMP-7: <0.0001
- CAMP-8: <0.0001
- CAMP-9: <0.0001
- CAMP-10: <0.0001
- CAMP-11: <0.0001
- CAMP-12: <0.0001
- CAMP-13: >0.9999 (not significant, n.s.)
- CAMP-14: >0.9999 (n.s.)
- CAMP-15: >0.9999 (n.s.)

- CAMP-16: >0.9999 (n.s.)
- CAMP-17: <0.0001
- CAMP-18: <0.0001
- CAMP-19: <0.0001
- CAMP-20: <0.0001
- 100 nM PMA: <0.0001

**Reporter assay in Supplementary Figure 7a**

(Condition: adjusted p value for comparison untreated vs. indicated condition; 14 comparisons in one family, n = 3 per group, df = 30, α = 0.05)

- CAMP-7: >0.9999 (not significant, n.s.)
- CAMP-8: <0.0001
- CAMP-22: <0.0001
- CAMP-21: <0.0001
- CAMP-23: <0.0001
- IL-2, 100 IU/mL: <0.0001
- IL-15, 10 ng/mL: <0.0001

**Reporter assay in Supplementary Figure 7b**

(Condition: adjusted p value for comparison untreated vs. indicated condition; 7 comparisons in one family, n = 3 per group, df = 16, α = 0.05)

- CAMP-7: <0.0001
- CAMP-8: <0.0001
- CAMP-22: <0.0001
- CAMP-21: <0.0001
- CAMP-23: <0.0001
- 100 nM PMA: <0.0001

**Reporter assay in Supplementary Figure 7c**

(Condition: adjusted p value for comparison untreated vs. indicated condition; 15 comparisons in one family, n = 3 per group, df = 32, α = 0.05)

- CAMP-7: >0.9999 (not significant, n.s.)
- CAMP-8: <0.0001
- CAMP-23: <0.0001
- CAMP-24: <0.0001
- CAMP-25: <0.0001
- CAMP-26: <0.0001
- IL-2, 100 IU/mL: 0.0001
- IL-15, 10 ng/mL: <0.0001

**Reporter assay in Supplementary Figure 7d**

(Condition: adjusted p value for comparison untreated vs. indicated condition; 8 comparisons in one family, n = 3 per group, df = 18, α = 0.05)

- CAMP-7: <0.0001
- CAMP-8: <0.0001
- CAMP-23: <0.0001
- CAMP-24: <0.0001
- CAMP-25: 0.5100 (not significant, n.s.)
- CAMP-26: 0.1736 (n.s.)
- PMA, 100 nM: <0.0001

**Supplementary Note 3. Automating a GUI-based image-analysis pipeline with an AI coding assistant**.

**Overview.** Interactive image-analysis programs such as Fiji/ImageJ and TrackMate are well suited to analyzing individual movies, but their manual, click-based operation is difficult to apply at scale. Processing many movies by hand is slow and prone to inconsistency, the exact sequence of settings is hard to record, and re-analyzing a dataset with a different parameter requires repeating the entire procedure. This note describes how we converted our manual TrackMate cell-tracking workflow into an automated pipeline with the assistance of an AI coding tool (Claude Code, Anthropic). It is intended for researchers who use Fiji or comparable software but who have not previously written scripts or worked at a command line, and it assumes no programming background. Cell tracking serves as the example, but the same procedure applies to most analyses carried out through a graphical interface.

**Procedure**. The procedure comprises six steps. The first two are carried out by the researcher, who defines the analysis; the AI assistant performs the coding in the third and fourth; and the last two are performed together. Specific terminology for this approach is provided in **Supplementary Table 2.**

1. Perform the complete analysis once in the graphical interface and record every step, including the value entered in each dialog box. For the work described here we captured a series of 28 annotated screenshots documenting movie import, correction of the frame axis, spot detection, the spot-quality threshold (2.0 for a permissive setting or 4.0 for a stringent one), the tracking distances, and each export operation. This record is the specification that the assistant reproduces, and greater detail produces a more faithful result.
2. Write a short description of the required inputs and outputs, and save one representative example of each. We recorded that the pipeline should accept AVI movies and produce a spot-statistics table (CSV), a track-overlay video, and a TIFF image, and we included the first several lines of a manually exported CSV to define the exact output format. Supplying a real example is the most reliable way to obtain a matching result, because the assistant can compare its output against a known-correct file.
3. Provide the screenshots, the description, and the example files to the assistant, and ask it to reproduce the workflow as a script that runs without the graphical interface. The assistant examines the supplied files, writes the script, runs it, and reports the outcome. Reading the resulting code is not necessary to begin, though it is advisable (see step 5).
4. Ask the assistant to wrap the script in a Makefile so that the full set of movies can be processed with a single command, the outputs are collected in one location, and the main settings can be changed from the command line. This converts an individual script into a reusable pipeline: the make command then processes every movie, and a command such as make -B QUALITY=2.0 re-runs the analysis with a new threshold.
5. Compare the automated output against the manual result before relying on the pipeline. Automatically generated code can appear correct while producing subtly wrong results, and a direct comparison against a trusted manual analysis is the only dependable check. We compared the two analyses for one movie and required them to agree. Because TrackMate assigns new internal identifiers on each run, the comparison was made on the basis of spot position rather than identifier; the automated and manual analyses yielded the same 228 tracks with identical membership, and all 2,478 tracked spots corresponded one to one. The comparison also revealed two errors that were then corrected: the exported table was initially tab-delimited rather than comma-delimited, which opened without complaint but was misread by the downstream step, and the automated export initially retained detection artifacts that the manual export excludes.
6. Once the pipeline has been validated, refine it as needed. Parameter changes that previously required hours of repeated manual operations become single commands, which allowed us to evaluate detection and tracking settings across the entire dataset and to observe their effect on the aggregate motility statistics rather than inferring it from a single movie. We also connected a second analysis stage, the calculation of mean-squared displacement, so that raw movies are processed through to final plots in one command.

**Application to cell tracking**. The assistant used here was Claude Code (Anthropic; model Claude Opus 4.7, May & June 2026), directed interactively, with the author providing guidance and review throughout.

The resulting pipeline comprises a Jython macro (trackmate_batch.py) that reproduces the manual TrackMate procedure (Laplacian-of-Gaussian detection with a 5 px radius and sub-pixel localization at a quality threshold of 4.0; Simple LAP tracking with 30 px linking, 15 px gap closing, and a maximum gap of two frames) and writes, for each movie, a spot-statistics table, a track-overlay AVI and TIFF, and a TrackMate session file. A Makefile locates the Fiji installation, identifies the input movies whether they are stored loosely in a folder or grouped into per-batch or per-sample subfolders, processes them, and passes the resulting tables to the downstream mean-squared-displacement analysis (compute_msd.py and plot_msd.py). The complete dataset is regenerated from the raw movies with a single make command. The code, the workflow screenshots, and the input description are archived in the project repository on Github (see **Supporting Information** section of main manuscript).

**Practical considerations**. Several practical points should be emphasized. Providing a genuine example of each output is more effective than describing it in words. Validation against a manual result should be treated as a required step rather than an optional one, since plausible output may still be incorrect. Comparisons should be made on physically meaningful quantities such as position, frame, or intensity, because internal identifiers may differ between runs even when the underlying results are identical. The generated code should be read by the researcher or reviewed by a colleague with programming experience, as the assistant is an efficient means of implementation but not a substitute for judgment about how an analysis should be performed. Every analysis setting should be made adjustable from the command line so that results remain reproducible and parameters can be varied systematically. Finally, software versions should be recorded, since differences among versions of Fiji, the build tool, or the supporting language can affect the results; the present work used macOS on Apple Silicon, Fiji/TrackMate, GNU Make 3.81, and Python 3 with NumPy and pandas for the mean-squared-displacement stage.

**Extending the approach to other analyses**. The approach is not specific to Fiji or to image analysis. Any analysis that can be expressed as code, and whose software provides a scripting interface, a macro system, or a means of exporting data for external processing, can be automated in the same way. In synthetic biology this includes flow-cytometry gating and population statistics, plate-reader growth and expression kinetics, microscopy segmentation and quantification, and sequencing-based readouts, among others. The essential elements are the same in every case: document the manual procedure and its settings, supply representative input and output examples, delegate the implementation to the assistant, and validate the automated result against a trusted manual analysis before applying it broadly. The initial effort of documenting and validating a workflow is recovered whenever the analysis must be applied to further samples or repeated with revised parameters.

**Getting started**. Getting started requires a computer with the relevant analysis software installed, access to Claude Code or a comparable AI coding assistant, and one manually completed analysis to serve as both the specification and the validation target. The expertise that cannot be delegated is knowledge of the correct analysis; the programming can be. Researchers who wish to apply this approach but are unsure how to begin are welcome to contact Xavier Bower.

**Supplementary Table 2. Automated image analysis pipeline terminology.**

| Term | Definition |
|---|---|
| **AI coding assistant (Claude Code)** | A command-line program that accepts instructions in ordinary English, reads and writes the files in a project folder, generates and runs code, and reports the results. It revises its work in response to feedback. |
| **Command line** | A text interface for issuing instructions to the computer in place of graphical controls. On macOS it is provided by the Terminal / iTerm2 application. |
| **Script (macro)** | A text file containing a sequence of instructions that carries out an analysis automatically. Fiji executes scripts written in the Jython language. |
| **Headless execution** | Running Fiji without its graphical interface, so that analyses proceed in the background and no dialog boxes require attention. This is what allows many files to be processed in one run. |
| **Makefile** | A configuration file, executed with the make command, that locates input files, applies a defined procedure to each, and regenerates only the outputs whose inputs have changed. |
| **Parameter** | An analysis setting, such as the spot-quality threshold, that is exposed for adjustment without editing the underlying code. |

**Supplementary Note 4. Advice for developing an analysis method with an AI coding assistant**

**Overview**. **Supplementary Note 3** describes automating an image-analysis procedure that had already been decided: the manual TrackMate workflow existed, and the task was to reproduce it in code. This note covers development of other analytical methods used in the project, where the analysis itself was still open to definition. When the motility data were first obtained, we had not yet decided how motility should be summarized, for example whether the cells are better described as two distinct populations, or states, than by one cut-off, and which comparisons the experiment could support. Ultimately, the research team defined a set of guiding questions and determined which analysis could answer such questions, and an AI coding assistant (Claude Code, Anthropic) wrote and ran the analysis.

One pass takes minutes, and a session contains tens of passes (**Supplementary Figure 18**). When moving to an assistant-enabled exploration process, trying out an analysis went from a day of work to a few minutes, so we could afford to test many choices. The assistant did not decide which analysis was correct. Every choice in this paper was made by the research team, and several choices were made by rejecting what the assistant first produced. Because nothing is retained between sessions of interaction with the assistant, each session ends by writing its conclusions and caveats into a numbered notebook entry that the next session reads: ten entries make up the analytical record for this project. Specific terminology for this approach is provided in **Supplemental Table 3.**

**Procedure**. The human research team leads the first three steps, the assistant does the fourth step, and the last two steps involve collaboration between the researchers and the assistant.

1. **Define your metric of interest specifically.** Asking for a plot leaves the underlying quantity implicit, and the assistant will pick one for you. When we asked instead what the fold change metric between two conditions should be, seven reasonable ways of computing it disagreed by up to sevenfold, and on 9 of 22 comparisons they disagreed about which direction the effect went. Choosing among them is a scientific judgement.
2. **Give the assistant the facts it cannot work out from the data.** Pixel size is the clearest case: our phase contrast images captured at 3 min intervals are 0.686 µm/pixel, and our phase contrast and fluorescence images captured at 2 min intervals are 1.029 µm/pixel. MSD depends on the square of that number, so the wrong value inflates every result by 2.25-fold and looks like a real difference between imaging channels. Frame intervals and treatment doses matter the same way. A mislabeled (by 1000x) dose in the filenames was originally interpreted as the true dose by the assistant and had to be corrected.
3. **Write the analysis plan down before running the comparisons.** Cheap analysis removes the friction that used to limit how many versions you tried, but that friction is useful in many cases—it encourages careful thought. Here, we recorded our readouts, our comparisons and our correction for testing many at once before computing any of them. This rigor differentiates choosing an appropriate test from choosing the most convenient looking.
4. **Have the assistant implement good coding practices.** In many cases, we asked for the analysis to be written as library functions with their own tests, with the scripts around them only reading files, writing files and printing a summary. Insisting on this early allowed later sessions to reproducibly leverage finalized code from earlier sessions for relevant parts of analysis without having to repeat implementation and testing.
5. **Check all results against possible claims.** The agent will attempt to interpret the data, often appropriately explaining limitations and suggesting further analysis or experiments that could clarify possible claims. However, critical review of all possible claims is important—unchallenged ideas will be kept in agent context, so incorrect interpretations need to be pushed back against. Furthermore, the agent has limited spatio-visual capabilities compared to text-based capabilities, and may fail to notice or underprioritize important results, which must be highlighted and explained by the human reviewer. In addition, the reviewer bears the responsibility for confirming accuracy of assembly, for instance ensuring that numbers in captions are consistent with the numbers in figures, and that stated claims are well supported as opposed to plausible.

6. **Write down what happened, including what is unresolved.** A tidy summary is less useful than one carrying the caveats and any disagreement left open, because the next session needs to inherit the doubts along with the conclusions. Agents have large context windows and could benefit from additional information in a way that a human researcher might not.

**Practical advice for researcher-agent interactions**

- State what counts as an independent measurement. The assistant will test at whatever level the data happens to be organized in (e.g. cells vs. wells), and this can produce confident wrong answers.
- Write the plan down before running the comparisons.
- Report and record what a negative result could have detected, rather than the negative result alone.
- Take pixel sizes, frame intervals and doses from the bench record, not from file names.
- Keep figure styling choices in one reference file so that panels stay consistent when one is revised.
- Treat anything existing only inside an agent session as potentially losable until it's written to files. Our shared style file survived a restart because it was a file; some code that used it did not, and it had to be rewritten.
- Record software and assistant versions used, and disclose the use of the tool.

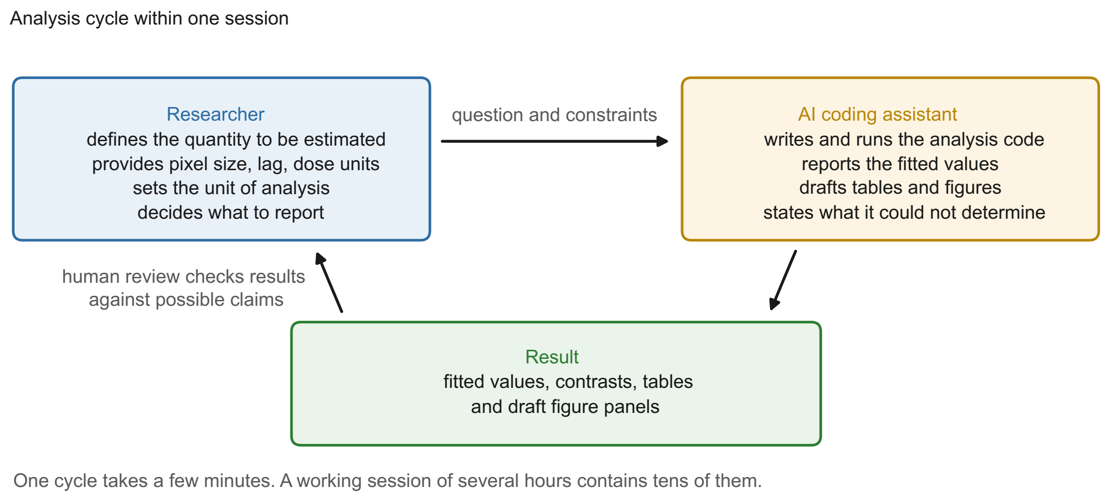


**Supplementary Figure S18. A human-in-the-loop process for analytical method development.** Summary of a typical development cycle used in this study. The researcher states the question and supplies the constraints to which the answer must adhere, including pixel size, frame interval, doses and what counts as an independent measurement. The assistant (e.g., Claude Code; see **Supplementary Note 3**) writes and runs the analysis and reports apparent conclusions, including anything it could not determine from the data. The researcher then checks the result to determine what claims are supported, and what questions should be asked to refine or clarify the claims with further analysis rounds.

**Supplementary Table 3. Human-in-the-loop process for analytical development terminology.**

| Term | Definition |
|---|---|
| **Session** | One working period with the assistant, usually a few hours. Nothing is retained between sessions, so anything a later session needs has to be written to a file. |
| **Notebook entry** | The numbered file recording one piece of work: the instructions given to the assistant, the results, and the conclusions and caveats. It is both the laboratory record and the input to the next session. |
| **Unit of analysis** | The level at which measurements are independent, and therefore the level at which they are counted. Here it is the well, not the individual cell. |
| **Readout** | One number per well, such as the median MSD or the center of the fast state, used to compare conditions. |

**Supplementary Table 4. Imaging, tracking, and motility-analysis parameters: Experiment 1 (phase contrast, Figure 4).[a,b]**

| Parameter | Setting | Physical value / note |
|---|---|---|
| **Acquisition** | | |
| Imaging modality | Phase contrast | Single channel |
| File format | 8-bit AVI | Grayscale |
| Frame size | 960 × 720 px | ≈ 659 × 494 μm field of view |
| Pixel size (UM_PER_PIXEL) | 0.68626 μm/px | — |
| Frame interval | 3 min | 180 s |
| Number of frames | 20 | — |
| Sequence duration | ≈ 57 min | 20 frames × 3 min interval (19 intervals) |
| **Detection (TrackMate LoG detector)** | | |
| Estimated blob diameter (DIAMETER) | 36.0 px | 24.7 μm |
| Detection threshold (THRESHOLD) | 0.20 | — |
| Quality threshold (QUALITY) | 0 (off) | No quality filter applied |
| Median filtering (MEDIAN_FILTERING) | On | — |
| Sub-pixel localization (SUBPIXEL) | On | — |
| **Tracking (LAP tracker)** | | |
| Tracker (TRACKER) | LAP | Linear assignment problem |
| Max linking distance (LINKING_MAX_DISTANCE) | 46 px | 31.6 μm |
| Max gap-closing distance (GAP_CLOSING_MAX_DISTANCE) | 55 px | 37.7 μm |
| Max frame gap (MAX_FRAME_GAP) | 2 frames | 6 min |
| **MSD / motility analysis** | | |
| MSD lag (LAG) | 3 frames | 9 min |
| Min. consecutive frames (MIN_CONSEC) | 12 frames [c] | 36 min |
| Motile threshold (MOTILE_THRESHOLD) | MSD > 25 μm² | — |
| Primary MSD endpoint | msd_overlap_partial | Overlapping-window, time-averaged MSD at the 9-min lag |

[a] Cells were imaged by phase contrast and tracked in TrackMate v7; mean squared displacement (MSD) was computed per cell at a fixed frame lag. Parameter names in parentheses correspond to the analysis pipeline's Makefile variables.

[b] Physical distances = pixel value × 0.68626 μm/px; field of view = frame size × pixel size; lag and gap times = frame count × 3 min frame interval.

[c] MIN_CONSEC constrains only the "complete" MSD variants (msd_overlap_complete, msd_single_complete); the reported endpoint (msd_overlap_partial) requires only one valid lag-window and is not bounded by it. Results reported in primary figures constitute msd_overlap_partial data.

**Supplementary Table 5. Imaging, tracking, and motility-analysis parameters: Experiment 2 (two-channel: GFP and phase contrast, Figure 6).**[a,b]

| Parameter | Phase contrast | GFP |
|---|---|---|
| **Acquisition (both channels)** | | |
| File format | 8-bit AVI (grayscale) | |
| Frame size | 640 × 480 px (≈ 659 × 494 µm field of view) | |
| Pixel size (UM_PER_PIXEL) | 1.02939 µm/px | |
| Frame interval | 2 min (120 s) | |
| Number of frames | 30 | |
| Sequence duration | ≈ 58 min (30 frames × 2 min interval, 29 intervals) | |
| **Detection (TrackMate LoG detector)** | | |
| Estimated blob diameter (DIAMETER) | 24.0 px (24.7 µm) | 20.0 px (20.6 µm) |
| Detection threshold (THRESHOLD) | 0.20 | |
| Quality threshold (QUALITY) | 0 (off) | |
| Median filtering (MEDIAN_FILTERING) | On | |
| Sub-pixel localization (SUBPIXEL) | On | |
| **Tracking (LAP tracker)** | | |
| Tracker (TRACKER) | LAP (linear assignment problem) | |
| Max linking distance (LINKING_MAX_DISTANCE) | 25 px (25.7 µm) | |
| Max gap-closing distance (GAP_CLOSING_MAX_DISTANCE) | 30 px (30.9 µm) | |
| Max frame gap (MAX_FRAME_GAP) | 2 frames (4 min) | |
| **MSD / motility analysis** | | |
| MSD lag (LAG) | 5 frames (10 min) | |
| Min. consecutive frames (MIN_CONSEC) | 12 frames (24 min) [c] | |
| Motile threshold (MOTILE_THRESHOLD) | MSD > 25 µm² | |
| Primary MSD endpoint | msd_overlap_partial (overlapping-window, time-averaged MSD at the 10-min lag) | |

[a] The same fields were imaged in two channels and each channel was tracked with using a separate Makefile command. Settings were identical across channels except the detection blob diameter (see Detection). Names in parentheses correspond to the analysis pipeline's Makefile variables.

[b] Physical distances = pixel value × 1.02939 µm/px; field of view = frame size × pixel size; lag and gap times = frame count × 2 min frame interval.

[c] MIN_CONSEC constrains only the "complete" MSD variants (msd_overlap_complete, msd_single_complete); the reported endpoint (msd_overlap_partial) requires only one valid lag-window and is not bounded by it.

**Supplementary Table 6. Flow cytometry laser configurations and settings.**

| | | | | |
|---|---|---|---|---|
| **355 nm** UV | **BUV 396** | — | 379/28 | BUV 395 |
| | **DAPI** | 410 LP | 450/50 | DAPI, Hoechst, eBFP, Calcein Blue AM, FVS 440UV, mTagBFP2, LIVE/DEAD Fixable Blue |
| **405 nm** Violet | **Pacific Blue** | — | 450/50 | Pacific Blue, Alexa Fluor 405, BV421, FVS450, V450, LIVE/DEAD Fixable Violet |
| | **AmCyan** | 505 LP | 525/50 | AmCyan, BV510, FVS510, V500, Pacific Orange |
| | **BV605** | 595 LP | 600/30 | BV605, Qdot 605, FVS575V |
| | **BV650** | 635 LP | 670/30 | BV650, Qdot 655 |
| | **BV711** | 685 LP | 730/45 | BV711, BV750 |
| **488 nm** Blue | **SSC** | — | 488/10 | *Side scatter (light-scatter detector; not a fluorescence channel)* |
| | **FITC** | 505 LP | 530/30 | FITC, Alexa Fluor 488, eGFP, eYFP, CFSE, Calcein AM, FVS 520, mNeonGreen, fluorescein |
| | **PerCP** | 685 LP | 716/40 | PerCP-Cy5.5, PerCP, RB705 [a] |
| **552 nm** Yellow-Green | **PE** | — | 582/15 | PE, tdTomato, eRFP, Alexa Fluor 561, FVS570, RY586, Cy3 |
| | **PE-TexasRed** | 600 LP | 610/20 | PE-Texas Red, mCherry, propidium iodide, Alexa Fluor 594, FVS620, RY610, DsRed-Express |
| | **PE-Cy5** | 635 LP | 670/20 | PE-Cy5, 7-AAD, RY655 [b] |
| | **PE-Cy7** | 750 LP | 780/60 | PE-Cy7, RY743, RY775 |
| **640 nm** Red | **APC** | — | 670/30 | APC, Alexa Fluor 647, Cy5, DRAQ5, DRAQ7, FVS 660, DyLight 650 |
| | **AlexaFluor700** | 685 LP | 730/45 | Alexa Fluor 700, APC-Cy5.5, DRAQ5, DRAQ7 |
| | **APC-Cy7** | 750 LP | 780/60 | APC-Cy7, DRAQ7, Alexa Fluor 750 |
| **685 nm** Far Red | **Alexa750** | — | 787/42 | Alexa Fluor 750, APC-Cy7, FVS 780 [c] |

**Notes**

— The first detector in each laser array collects emitted light directly and has no longpass mirror in the path.

[a] For PerCP: replace the 716/40 bandpass with 670/30 and the 685 LP mirror with 635 LP.

[b] For PE-Cy5.5: replace the 670/20 bandpass with 720/20.

[c] For Alexa Fluor 700: replace the 787/42 bandpass with 720/20.

*Bandpass filters are given as center wavelength / bandwidth in nm (e.g. 530/30 transmits 515–545 nm). Fluorochrome lists are representative, not exhaustive.*
