## Supplementary Data 1 for "Genetically inducible coordinators of cytokine signaling pathways for interrogating T cell motility": 8WTJWN-methods.txt

Methods

Quality of the fastq files was assessed using FastQC v0.12.1. Reads were then
quality filtered using fastp v0.24.0 with poly-X tail trimming, 3' quality-based
tail trimming, a minimum Phred quality score of 15, and a minimum length
requirement of 50 bp. Quality-filtered reads were aligned to the reference genome
using STAR aligner v2.7.11 with non-canonical splice junction removal and output
of unmapped reads, followed by coordinate sorting using samtools v1.22.1. PCR and
optical duplicates were removed using UMI-based deduplication with UMIcollapse
v1.1.0. Alignment quality metrics, strand specificity, and read distribution
across genomic features were assessed using RSeQC v5.0.4 and Qualimap v2.3, with
results aggregated into a comprehensive quality control report using MultiQC
v1.32. Gene-level expression quantification was performed using featureCounts
(subread package v2.1.1) with strand-specific counting, multi-mapping read
fractional assignment, exons and three prime UTR as the feature identifiers, 
and grouped by gene_id. Final gene counts were annotated with gene biotype and 
other metadata extracted from the reference GTF file.  Sample-sample correlations 
for sample-sample heatmap and PCA were calculated on normalized counts 
(TMM, trimmed mean of M-values) using Pearson correlation. Differential expression 
was done with edgeR v4.0.16 using standard practice including filtering for 
low-expressed genes with edgeR::filterByExprwith default values. Functional 
enrichment, when available, is performed using gene set enrichment analysis with 
gseapy v0.12 using the MSigDB Hallmark gene set.
