## Supplementary Data 1 for "Genetically inducible coordinators of cytokine signaling pathways for interrogating T cell motility": 8WTJWN_2_pYRS0239_500DOX1.html

Loading report..

Report
generated on 2026-08-05, 21:51 UTC
based on data in:

- `/tmp/nxf.aRc1pe5tTN/rustqc`
- `/tmp/nxf.aRc1pe5tTN/8WTJWN_2_filtered_fastqc.zip`
- `/tmp/nxf.aRc1pe5tTN/8WTJWN_2_QC-short-reads.json`
- `/tmp/nxf.aRc1pe5tTN/8WTJWN_2_Log.final.out`
- `/tmp/nxf.aRc1pe5tTN/8WTJWN_2_feature-counts.tsv.summary`

Copy Prompt


Summarize table

| Overrepresented sequence | Reports | Occurrences | % of all reads |
| --- | --- | --- | --- |
| GTACATTCCACAAGCATTGCCTTCTTATTTTACTTCTTTTAGCTGTTTAA | 1 | 23506 | 0.1153% |
| CCCTAATACCTGCCACCCCACTCTTAATCAGTGGTGGAAGAACGGTCTCA | 1 | 21539 | 0.1056% |

Close
