## Supplementary Data 1 for "Genetically inducible coordinators of cytokine signaling pathways for interrogating T cell motility": 8WTJWN_3_pYRS0239_PMA1.html

Loading report..

Report
generated on 2026-08-05, 21:20 UTC
based on data in:

- `/tmp/nxf.2C81Wrl75s/rustqc`
- `/tmp/nxf.2C81Wrl75s/8WTJWN_3_filtered_fastqc.zip`
- `/tmp/nxf.2C81Wrl75s/8WTJWN_3_QC-short-reads.json`
- `/tmp/nxf.2C81Wrl75s/8WTJWN_3_Log.final.out`
- `/tmp/nxf.2C81Wrl75s/8WTJWN_3_feature-counts.tsv.summary`

Copy Prompt


Summarize table

| Overrepresented sequence | Reports | Occurrences | % of all reads |
| --- | --- | --- | --- |
| CCCTAATACCTGCCACCCCACTCTTAATCAGTGGTGGAAGAACGGTCTCA | 1 | 25492 | 0.1333% |
| GTACATTCCACAAGCATTGCCTTCTTATTTTACTTCTTTTAGCTGTTTAA | 1 | 24574 | 0.1285% |
| CTCTTAATCAGTGGTGGAAGAACGGTCTCAGAACTGTTTGTTTCAATTGG | 1 | 19651 | 0.1027% |
| GGCCCAAGGTGTCCTGCAGGCTGTAATGCAGTTTAATCAGAGTGCCATTT | 1 | 19508 | 0.1020% |

Close
