## Supplementary Data 1 for "Genetically inducible coordinators of cytokine signaling pathways for interrogating T cell motility": 8WTJWN_7_pYRS0255_500DOX1.html

MultiQC Report


Loading report..

v1.33


Theme

- Light
- Dark
- Auto

Highlight

 Rename

 Show / Hide


AI Analysis

 Export

 Settings

 Citations

 About

- General Stats
- QualiMap
  - Genomic origin of reads
  - Gene Coverage Profile
- RSeQC
  - Read Distribution
  - Infer experiment
- featureCounts
- STAR
  - Summary Statistics
  - Alignment Scores
- FastQC
  - Sequence Quality Histograms
  - Per Sequence Quality Scores
  - Per Base Sequence Content
  - Per Sequence GC Content
  - Per Base N Content
  - Sequence Length Distribution
  - Overrepresented sequences by sample
  - Adapter Content
  - Status Checks
- Software Versions

Loading report..

Report
generated on 2026-08-05, 21:53 UTC
based on data in:

- `/tmp/nxf.4Ys4yyPebK/rustqc`
- `/tmp/nxf.4Ys4yyPebK/8WTJWN_7_filtered_fastqc.zip`
- `/tmp/nxf.4Ys4yyPebK/8WTJWN_7_QC-short-reads.json`
- `/tmp/nxf.4Ys4yyPebK/8WTJWN_7_Log.final.out`
- `/tmp/nxf.4Ys4yyPebK/8WTJWN_7_feature-counts.tsv.summary`

##### AI Summary

Provider: , model:

Chat with Seqera AI

1 samples had less than 1% of reads made up of overrepresented sequences

### Adapter Content 1 Help

The cumulative percentage count of the proportion of your
library which has seen each of the adapter sequences at each position.

Close
