## Supplementary Data 1 for "Genetically inducible coordinators of cytokine signaling pathways for interrogating T cell motility": 8WTJWN_8_pYRS0257_UNT1.html

Loading report..

Report
generated on 2026-08-05, 21:52 UTC
based on data in:

- `/tmp/nxf.wBcgPsbbwe/rustqc`
- `/tmp/nxf.wBcgPsbbwe/8WTJWN_8_filtered_fastqc.zip`
- `/tmp/nxf.wBcgPsbbwe/8WTJWN_8_QC-short-reads.json`
- `/tmp/nxf.wBcgPsbbwe/8WTJWN_8_Log.final.out`
- `/tmp/nxf.wBcgPsbbwe/8WTJWN_8_feature-counts.tsv.summary`

Copy Prompt


Summarize table

| Overrepresented sequence | Reports | Occurrences | % of all reads |
| --- | --- | --- | --- |
| GTACATTCCACAAGCATTGCCTTCTTATTTTACTTCTTTTAGCTGTTTAA | 1 | 24888 | 0.1233% |
| CCCTAATACCTGCCACCCCACTCTTAATCAGTGGTGGAAGAACGGTCTCA | 1 | 22933 | 0.1136% |

Close
