## Supplementary Data 1 for "Genetically inducible coordinators of cytokine signaling pathways for interrogating T cell motility": 8WTJWN_10_Naive_UNT1.html

Loading report..

Report
generated on 2026-08-05, 21:50 UTC
based on data in:

- `/tmp/nxf.VqQaAS3ftM/rustqc`
- `/tmp/nxf.VqQaAS3ftM/8WTJWN_10_filtered_fastqc.zip`
- `/tmp/nxf.VqQaAS3ftM/8WTJWN_10_QC-short-reads.json`
- `/tmp/nxf.VqQaAS3ftM/8WTJWN_10_Log.final.out`
- `/tmp/nxf.VqQaAS3ftM/8WTJWN_10_feature-counts.tsv.summary`

Copy Prompt


Summarize table

| Overrepresented sequence | Reports | Occurrences | % of all reads |
| --- | --- | --- | --- |
| CCCTAATACCTGCCACCCCACTCTTAATCAGTGGTGGAAGAACGGTCTCA | 1 | 38206 | 0.1958% |
| CTCTTAATCAGTGGTGGAAGAACGGTCTCAGAACTGTTTGTTTCAATTGG | 1 | 30530 | 0.1565% |

Close
