## Supplementary Data 1 for "Genetically inducible coordinators of cytokine signaling pathways for interrogating T cell motility": 8WTJWN_11_Naive_500DOX1.html

Loading report..

Report
generated on 2026-08-05, 21:59 UTC
based on data in:

- `/tmp/nxf.2niPYGt9LU/rustqc`
- `/tmp/nxf.2niPYGt9LU/8WTJWN_11_filtered_fastqc.zip`
- `/tmp/nxf.2niPYGt9LU/8WTJWN_11_QC-short-reads.json`
- `/tmp/nxf.2niPYGt9LU/8WTJWN_11_Log.final.out`
- `/tmp/nxf.2niPYGt9LU/8WTJWN_11_feature-counts.tsv.summary`

Copy Prompt


Summarize table

| Overrepresented sequence | Reports | Occurrences | % of all reads |
| --- | --- | --- | --- |
| CCCTAATACCTGCCACCCCACTCTTAATCAGTGGTGGAAGAACGGTCTCA | 1 | 37822 | 0.1866% |
| CTCTTAATCAGTGGTGGAAGAACGGTCTCAGAACTGTTTGTTTCAATTGG | 1 | 30645 | 0.1512% |

Close
