## Supplementary Data 1 for "Genetically inducible coordinators of cytokine signaling pathways for interrogating T cell motility": 8WTJWN_12_Naive_PMA.html

Loading report..

Report
generated on 2026-08-05, 21:56 UTC
based on data in:

- `/tmp/nxf.r629aSS8eS/rustqc`
- `/tmp/nxf.r629aSS8eS/8WTJWN_12_filtered_fastqc.zip`
- `/tmp/nxf.r629aSS8eS/8WTJWN_12_QC-short-reads.json`
- `/tmp/nxf.r629aSS8eS/8WTJWN_12_Log.final.out`
- `/tmp/nxf.r629aSS8eS/8WTJWN_12_feature-counts.tsv.summary`

Copy Prompt


Summarize table

| Overrepresented sequence | Reports | Occurrences | % of all reads |
| --- | --- | --- | --- |
| CCCTAATACCTGCCACCCCACTCTTAATCAGTGGTGGAAGAACGGTCTCA | 1 | 37029 | 0.1899% |
| CTCTTAATCAGTGGTGGAAGAACGGTCTCAGAACTGTTTGTTTCAATTGG | 1 | 29085 | 0.1491% |
| GTACATTCCACAAGCATTGCCTTCTTATTTTACTTCTTTTAGCTGTTTAA | 1 | 23578 | 0.1209% |

Close
