## Supplementary Data 1 for "Genetically inducible coordinators of cytokine signaling pathways for interrogating T cell motility": TYJH7L-multiqc-report.html

Loading report..

Report
generated on 2026-08-05, 21:59
based on data in:

- `/tmp/tmprl1uvqtq/TYJH7L_2_multiqc`
- `/tmp/tmprl1uvqtq/TYJH7L_3_multiqc`
- `/tmp/tmprl1uvqtq/TYJH7L_5_multiqc`
- `/tmp/tmprl1uvqtq/TYJH7L_6_multiqc`
- `/tmp/tmprl1uvqtq/TYJH7L_7_multiqc`
- `/tmp/tmprl1uvqtq/TYJH7L_8_multiqc`
- `/tmp/tmprl1uvqtq/TYJH7L_10_multiqc`
- `/tmp/tmprl1uvqtq/TYJH7L_11_multiqc`
- `/tmp/tmprl1uvqtq/TYJH7L_12_multiqc`
- `/tmp/tmprl1uvqtq/TYJH7L_4_multiqc`
- `/tmp/tmprl1uvqtq/TYJH7L_1_multiqc`
- `/tmp/tmprl1uvqtq/TYJH7L_9_multiqc`

Summarize report

Copy report prompt

**Welcome!** Not sure where to start?
Watch a tutorial video
*(6:06)*

don't show again

###### Report AI Summary

More details…

Provider: , model:

Chat with Seqera AI

### General Statistics

###### AI Summary

Provider: , model:

Chat with Seqera AI

Table
 Export...

Copy prompt


Summarize plot

Created with MultiQC

Copy table

 Configure columns

 Sort by highlight

 Scatter plot

 Violin plot
Export as CSV...
Showing ^36^/_36_ rows and ^7^/_22_ columns.

Copy Prompt


Summarize table

| Sample Name | 5'-3' bias | M Aligned | Exonic | Intronic | Intergenic | Overlapping Exon | TIN stdev | TIN | Assigned | Assigned | Total reads | Aligned | Aligned | Uniq aligned | Uniq aligned | Multimapped | Dups | GC | Avg len | Median len | Failed | Seqs |
| --- | --- | --- | --- | --- | --- | --- | --- | --- | --- | --- | --- | --- | --- | --- | --- | --- | --- | --- | --- | --- | --- | --- |
| TYJH7L\_1 |  |  |  |  |  |  |  |  |  |  | 8.9M | 8.4M | 94.2% | 7.6M | 84.8% | 0.8M |  |  |  |  |  |  |
| TYJH7L\_1\_dedup | 0.35 | 3.6M | 2.6M | 0.4M | 0.1M | 0.1M | 18.6 | 21.6 | 3.2M | 74.8% |  |  |  |  |  |  |  |  |  |  |  |  |
| TYJH7L\_1\_filtered |  |  |  |  |  |  |  |  |  |  |  |  |  |  |  |  | 78.1% | 43.0% | 92bp | 94bp | 27% | 8.9M |
| TYJH7L\_2 |  |  |  |  |  |  |  |  |  |  | 7.2M | 6.8M | 94.9% | 6.1M | 85.1% | 0.7M |  |  |  |  |  |  |
| TYJH7L\_2\_dedup | 0.24 | 2.9M | 2.1M | 0.3M | 0.1M | 0.1M | 17.7 | 19.2 | 2.7M | 75.3% |  |  |  |  |  |  |  |  |  |  |  |  |
| TYJH7L\_2\_filtered |  |  |  |  |  |  |  |  |  |  |  |  |  |  |  |  | 78.5% | 42.0% | 92bp | 94bp | 27% | 7.2M |
| TYJH7L\_3 |  |  |  |  |  |  |  |  |  |  | 3.3M | 3.0M | 90.7% | 2.7M | 80.4% | 0.3M |  |  |  |  |  |  |
| TYJH7L\_3\_dedup | 0.22 | 1.3M | 0.9M | 0.2M | 0.1M | 0.1M | 17.5 | 17.7 | 1.1M | 70.0% |  |  |  |  |  |  |  |  |  |  |  |  |
| TYJH7L\_3\_filtered |  |  |  |  |  |  |  |  |  |  |  |  |  |  |  |  | 73.8% | 42.0% | 92bp | 94bp | 27% | 3.3M |
| TYJH7L\_4 |  |  |  |  |  |  |  |  |  |  | 20.1M | 19.8M | 98.6% | 18.1M | 89.8% | 1.8M |  |  |  |  |  |  |
| TYJH7L\_4\_dedup | 0.41 | 11.7M | 8.6M | 1.5M | 0.2M | 0.4M | 20.6 | 30.0 | 10.6M | 76.9% |  |  |  |  |  |  |  |  |  |  |  |  |
| TYJH7L\_4\_filtered |  |  |  |  |  |  |  |  |  |  |  |  |  |  |  |  | 78.2% | 45.0% | 92bp | 94bp | 27% | 20.1M |
| TYJH7L\_5 |  |  |  |  |  |  |  |  |  |  | 19.1M | 18.8M | 98.7% | 17.2M | 90.0% | 1.7M |  |  |  |  |  |  |
| TYJH7L\_5\_dedup | 0.45 | 11.4M | 8.5M | 1.4M | 0.2M | 0.4M | 20.7 | 30.2 | 10.4M | 78.0% |  |  |  |  |  |  |  |  |  |  |  |  |
| TYJH7L\_5\_filtered |  |  |  |  |  |  |  |  |  |  |  |  |  |  |  |  | 77.9% | 45.0% | 93bp | 94bp | 27% | 19.1M |
| TYJH7L\_6 |  |  |  |  |  |  |  |  |  |  | 20.6M | 20.3M | 98.7% | 18.5M | 90.0% | 1.8M |  |  |  |  |  |  |
| TYJH7L\_6\_dedup | 0.51 | 11.8M | 9.0M | 1.3M | 0.2M | 0.4M | 20.6 | 29.8 | 11.0M | 79.6% |  |  |  |  |  |  |  |  |  |  |  |  |
| TYJH7L\_6\_filtered |  |  |  |  |  |  |  |  |  |  |  |  |  |  |  |  | 79.3% | 45.0% | 93bp | 94bp | 27% | 20.6M |
| TYJH7L\_7 |  |  |  |  |  |  |  |  |  |  | 19.9M | 19.7M | 98.8% | 18.0M | 90.4% | 1.7M |  |  |  |  |  |  |
| TYJH7L\_7\_dedup | 0.50 | 11.5M | 8.6M | 1.4M | 0.2M | 0.4M | 20.8 | 30.9 | 10.4M | 77.6% |  |  |  |  |  |  |  |  |  |  |  |  |
| TYJH7L\_7\_filtered |  |  |  |  |  |  |  |  |  |  |  |  |  |  |  |  | 77.9% | 45.0% | 93bp | 94bp | 27% | 19.9M |
| TYJH7L\_8 |  |  |  |  |  |  |  |  |  |  | 19.4M | 19.1M | 98.6% | 17.4M | 89.9% | 1.7M |  |  |  |  |  |  |
| TYJH7L\_8\_dedup | 0.46 | 10.3M | 7.6M | 1.3M | 0.2M | 0.4M | 20.6 | 29.6 | 9.3M | 77.3% |  |  |  |  |  |  |  |  |  |  |  |  |
| TYJH7L\_8\_filtered |  |  |  |  |  |  |  |  |  |  |  |  |  |  |  |  | 79.2% | 45.0% | 93bp | 94bp | 27% | 19.4M |
| TYJH7L\_9 |  |  |  |  |  |  |  |  |  |  | 19.3M | 19.1M | 98.6% | 17.4M | 90.0% | 1.7M |  |  |  |  |  |  |
| TYJH7L\_9\_dedup | 0.46 | 10.5M | 7.9M | 1.3M | 0.2M | 0.4M | 20.6 | 30.6 | 9.6M | 77.4% |  |  |  |  |  |  |  |  |  |  |  |  |
| TYJH7L\_9\_filtered |  |  |  |  |  |  |  |  |  |  |  |  |  |  |  |  | 78.5% | 45.0% | 92bp | 94bp | 27% | 19.3M |
| TYJH7L\_10 |  |  |  |  |  |  |  |  |  |  | 19.9M | 19.6M | 98.7% | 17.9M | 90.2% | 1.7M |  |  |  |  |  |  |
| TYJH7L\_10\_dedup | 0.56 | 12.2M | 9.1M | 1.5M | 0.2M | 0.4M | 20.9 | 31.5 | 11.0M | 77.4% |  |  |  |  |  |  |  |  |  |  |  |  |
| TYJH7L\_10\_filtered |  |  |  |  |  |  |  |  |  |  |  |  |  |  |  |  | 77.1% | 45.0% | 93bp | 94bp | 27% | 19.9M |
| TYJH7L\_11 |  |  |  |  |  |  |  |  |  |  | 18.9M | 18.6M | 98.7% | 17.0M | 90.0% | 1.6M |  |  |  |  |  |  |
| TYJH7L\_11\_dedup | 0.55 | 12.0M | 9.1M | 1.4M | 0.2M | 0.4M | 20.8 | 30.8 | 11.1M | 78.0% |  |  |  |  |  |  |  |  |  |  |  |  |
| TYJH7L\_11\_filtered |  |  |  |  |  |  |  |  |  |  |  |  |  |  |  |  | 77.0% | 45.0% | 93bp | 94bp | 27% | 18.9M |
| TYJH7L\_12 |  |  |  |  |  |  |  |  |  |  | 19.3M | 19.0M | 98.4% | 17.1M | 88.7% | 1.9M |  |  |  |  |  |  |
| TYJH7L\_12\_dedup | 0.41 | 9.5M | 7.3M | 0.9M | 0.2M | 0.4M | 20.3 | 29.0 | 9.0M | 78.6% |  |  |  |  |  |  |  |  |  |  |  |  |
| TYJH7L\_12\_filtered |  |  |  |  |  |  |  |  |  |  |  |  |  |  |  |  | 81.9% | 45.0% | 93bp | 94bp | 27% | 19.3M |

 Scatter plot

 Violin plot
Export as CSV...
Showing ^12^/_12_ rows and ^10^/_19_ columns.

Copy Prompt


Summarize table

| Sample Name | Total reads | Aligned | Aligned | Uniq aligned | Uniq aligned | Multimapped | Avg. read len | Avg. mapped len | Splices | Annotated splices | GT/AG splices | GC/AG splices | AT/AC splices | Non-canonical splices | Mismatch rate | Del rate | Del len | Ins rate | Ins len |
| --- | --- | --- | --- | --- | --- | --- | --- | --- | --- | --- | --- | --- | --- | --- | --- | --- | --- | --- | --- |
| TYJH7L\_1 | 8.9M | 8.4M | 94.2% | 7.6M | 84.8% | 0.8M | 91.0bp | 91.6bp | 1.4M | 1.4M | 1.4M | 0.0M | 0.0M | 0.0M | 0.2% | 0.0% | 1.6bp | 0.0% | 1.4bp |
| TYJH7L\_2 | 7.2M | 6.8M | 94.9% | 6.1M | 85.1% | 0.7M | 91.0bp | 91.4bp | 1.0M | 1.0M | 1.0M | 0.0M | 0.0M | 0.0M | 0.2% | 0.0% | 1.6bp | 0.0% | 1.5bp |
| TYJH7L\_3 | 3.3M | 3.0M | 90.7% | 2.7M | 80.4% | 0.3M | 91.0bp | 91.2bp | 0.5M | 0.5M | 0.5M | 0.0M | 0.0M | 0.0M | 0.3% | 0.0% | 1.6bp | 0.0% | 1.5bp |
| TYJH7L\_4 | 20.1M | 19.8M | 98.6% | 18.1M | 89.8% | 1.8M | 92.0bp | 91.9bp | 3.5M | 3.5M | 3.5M | 0.0M | 0.0M | 0.0M | 0.2% | 0.0% | 1.5bp | 0.0% | 1.4bp |
| TYJH7L\_5 | 19.1M | 18.8M | 98.7% | 17.2M | 90.0% | 1.7M | 92.0bp | 92.0bp | 3.3M | 3.3M | 3.3M | 0.0M | 0.0M | 0.0M | 0.2% | 0.0% | 1.5bp | 0.0% | 1.4bp |
| TYJH7L\_6 | 20.6M | 20.3M | 98.7% | 18.5M | 90.0% | 1.8M | 92.0bp | 92.1bp | 3.6M | 3.6M | 3.6M | 0.0M | 0.0M | 0.0M | 0.2% | 0.0% | 1.5bp | 0.0% | 1.4bp |
| TYJH7L\_7 | 19.9M | 19.7M | 98.8% | 18.0M | 90.4% | 1.7M | 92.0bp | 92.0bp | 3.4M | 3.4M | 3.4M | 0.0M | 0.0M | 0.0M | 0.2% | 0.0% | 1.5bp | 0.0% | 1.4bp |
| TYJH7L\_8 | 19.4M | 19.1M | 98.6% | 17.4M | 89.9% | 1.7M | 92.0bp | 92.0bp | 3.4M | 3.3M | 3.3M | 0.0M | 0.0M | 0.0M | 0.2% | 0.0% | 1.5bp | 0.0% | 1.4bp |
| TYJH7L\_9 | 19.3M | 19.1M | 98.6% | 17.4M | 90.0% | 1.7M | 92.0bp | 91.9bp | 3.4M | 3.3M | 3.3M | 0.0M | 0.0M | 0.0M | 0.2% | 0.0% | 1.5bp | 0.0% | 1.4bp |
| TYJH7L\_10 | 19.9M | 19.6M | 98.7% | 17.9M | 90.2% | 1.7M | 92.0bp | 92.0bp | 3.4M | 3.3M | 3.4M | 0.0M | 0.0M | 0.0M | 0.2% | 0.0% | 1.5bp | 0.0% | 1.4bp |
| TYJH7L\_11 | 18.9M | 18.6M | 98.7% | 17.0M | 90.0% | 1.6M | 92.0bp | 92.0bp | 3.3M | 3.3M | 3.3M | 0.0M | 0.0M | 0.0M | 0.2% | 0.0% | 1.5bp | 0.0% | 1.4bp |
| TYJH7L\_12 | 19.3M | 19.0M | 98.4% | 17.1M | 88.7% | 1.9M | 92.0bp | 92.0bp | 3.6M | 3.6M | 3.6M | 0.0M | 0.0M | 0.0M | 0.2% | 0.0% | 1.5bp | 0.0% | 1.4bp |

Copy Prompt


Summarize table

| Overrepresented sequence | Reports | Occurrences | % of all reads |
| --- | --- | --- | --- |
| CCCTAATACCTGCCACCCCACTCTTAATCAGTGGTGGAAGAACGGTCTCA | 12 | 300953 | 0.1536% |
| GGCCCAAGGTGTCCTGCAGGCTGTAATGCAGTTTAATCAGAGTGCCATTT | 12 | 287984 | 0.1470% |
| GTACATTCCACAAGCATTGCCTTCTTATTTTACTTCTTTTAGCTGTTTAA | 10 | 272268 | 0.1390% |
| CTCTTAATCAGTGGTGGAAGAACGGTCTCAGAACTGTTTGTTTCAATTGG | 10 | 177714 | 0.0907% |
| CTGTAATGCAGTTTAATCAGAGTGCCATTTTTTTTTTTGTTCAAATGATT | 1 | 22037 | 0.0112% |
| AATATGCACTGTACATTCCACAAGCATTGCCTTCTTATTTTACTTCTTTT | 1 | 19430 | 0.0099% |
| GTTTAAATGACTGTGCTGCCCCTTTCACATCAAAGAACTACTGACAACGA | 1 | 19308 | 0.0099% |
| GTAAAAGACTGGTTAATGATAACAATGCATCGTAAAACCTTCAGAAGGAA | 2 | 10966 | 0.0056% |

Close
