## Supplementary Data 1 for "Genetically inducible coordinators of cytokine signaling pathways for interrogating T cell motility": TYJH7L_1_pYRS0239_UNT2.html

Loading report..

Report
generated on 2026-08-05, 21:54 UTC
based on data in:

- `/tmp/nxf.jPqOVIlWVj/rustqc`
- `/tmp/nxf.jPqOVIlWVj/TYJH7L_1_filtered_fastqc.zip`
- `/tmp/nxf.jPqOVIlWVj/TYJH7L_1_QC-short-reads.json`
- `/tmp/nxf.jPqOVIlWVj/TYJH7L_1_Log.final.out`
- `/tmp/nxf.jPqOVIlWVj/TYJH7L_1_feature-counts.tsv.summary`

Copy Prompt


Summarize table

| Overrepresented sequence | Reports | Occurrences | % of all reads |
| --- | --- | --- | --- |
| CCCTAATACCTGCCACCCCACTCTTAATCAGTGGTGGAAGAACGGTCTCA | 1 | 14287 | 0.1598% |
| GGCCCAAGGTGTCCTGCAGGCTGTAATGCAGTTTAATCAGAGTGCCATTT | 1 | 13156 | 0.1471% |
| CTCTTAATCAGTGGTGGAAGAACGGTCTCAGAACTGTTTGTTTCAATTGG | 1 | 10854 | 0.1214% |
| GTACATTCCACAAGCATTGCCTTCTTATTTTACTTCTTTTAGCTGTTTAA | 1 | 9396 | 0.1051% |

Close
