## Supplementary Data 1 for "Genetically inducible coordinators of cytokine signaling pathways for interrogating T cell motility": TYJH7L_2_pYRS0239_500DOX2.html

Loading report..

Report
generated on 2026-08-05, 21:55 UTC
based on data in:

- `/tmp/nxf.4ZJysCQGN5/rustqc`
- `/tmp/nxf.4ZJysCQGN5/TYJH7L_2_filtered_fastqc.zip`
- `/tmp/nxf.4ZJysCQGN5/TYJH7L_2_QC-short-reads.json`
- `/tmp/nxf.4ZJysCQGN5/TYJH7L_2_Log.final.out`
- `/tmp/nxf.4ZJysCQGN5/TYJH7L_2_feature-counts.tsv.summary`

Copy Prompt


Summarize table

| Overrepresented sequence | Reports | Occurrences | % of all reads |
| --- | --- | --- | --- |
| CCCTAATACCTGCCACCCCACTCTTAATCAGTGGTGGAAGAACGGTCTCA | 1 | 13410 | 0.1871% |
| GGCCCAAGGTGTCCTGCAGGCTGTAATGCAGTTTAATCAGAGTGCCATTT | 1 | 11345 | 0.1583% |
| CTCTTAATCAGTGGTGGAAGAACGGTCTCAGAACTGTTTGTTTCAATTGG | 1 | 9829 | 0.1371% |
| GTAAAAGACTGGTTAATGATAACAATGCATCGTAAAACCTTCAGAAGGAA | 1 | 7601 | 0.1060% |

Close
