## Supplementary Data 1 for "Genetically inducible coordinators of cytokine signaling pathways for interrogating T cell motility": TYJH7L_3_pYRS0239_PMA2.html

Loading report..

Report
generated on 2026-08-05, 21:51 UTC
based on data in:

- `/tmp/nxf.8n8DitHxtF/rustqc`
- `/tmp/nxf.8n8DitHxtF/TYJH7L_3_filtered_fastqc.zip`
- `/tmp/nxf.8n8DitHxtF/TYJH7L_3_QC-short-reads.json`
- `/tmp/nxf.8n8DitHxtF/TYJH7L_3_Log.final.out`
- `/tmp/nxf.8n8DitHxtF/TYJH7L_3_feature-counts.tsv.summary`

Copy Prompt


Summarize table

| Overrepresented sequence | Reports | Occurrences | % of all reads |
| --- | --- | --- | --- |
| CCCTAATACCTGCCACCCCACTCTTAATCAGTGGTGGAAGAACGGTCTCA | 1 | 5492 | 0.1644% |
| GGCCCAAGGTGTCCTGCAGGCTGTAATGCAGTTTAATCAGAGTGCCATTT | 1 | 5076 | 0.1519% |
| CTCTTAATCAGTGGTGGAAGAACGGTCTCAGAACTGTTTGTTTCAATTGG | 1 | 3962 | 0.1186% |
| GTAAAAGACTGGTTAATGATAACAATGCATCGTAAAACCTTCAGAAGGAA | 1 | 3365 | 0.1007% |

Close
