## Supplementary Data 1 for "Genetically inducible coordinators of cytokine signaling pathways for interrogating T cell motility": TYJH7L_4_pYRS0240_UNT2.html

Loading report..

Report
generated on 2026-08-05, 22:03 UTC
based on data in:

- `/tmp/nxf.wljkBQI5un/rustqc`
- `/tmp/nxf.wljkBQI5un/TYJH7L_4_filtered_fastqc.zip`
- `/tmp/nxf.wljkBQI5un/TYJH7L_4_QC-short-reads.json`
- `/tmp/nxf.wljkBQI5un/TYJH7L_4_Log.final.out`
- `/tmp/nxf.wljkBQI5un/TYJH7L_4_feature-counts.tsv.summary`

Copy Prompt


Summarize table

| Overrepresented sequence | Reports | Occurrences | % of all reads |
| --- | --- | --- | --- |
| CCCTAATACCTGCCACCCCACTCTTAATCAGTGGTGGAAGAACGGTCTCA | 1 | 31889 | 0.1584% |
| GGCCCAAGGTGTCCTGCAGGCTGTAATGCAGTTTAATCAGAGTGCCATTT | 1 | 30361 | 0.1508% |
| GTACATTCCACAAGCATTGCCTTCTTATTTTACTTCTTTTAGCTGTTTAA | 1 | 29723 | 0.1476% |
| CTCTTAATCAGTGGTGGAAGAACGGTCTCAGAACTGTTTGTTTCAATTGG | 1 | 22220 | 0.1104% |

Close
