## Supplementary Data 1 for "Genetically inducible coordinators of cytokine signaling pathways for interrogating T cell motility": TYJH7L_5_pYRS0240_500DOX2.html

Loading report..

Report
generated on 2026-08-05, 21:53 UTC
based on data in:

- `/tmp/nxf.WEzLzfPWQm/rustqc`
- `/tmp/nxf.WEzLzfPWQm/TYJH7L_5_filtered_fastqc.zip`
- `/tmp/nxf.WEzLzfPWQm/TYJH7L_5_QC-short-reads.json`
- `/tmp/nxf.WEzLzfPWQm/TYJH7L_5_Log.final.out`
- `/tmp/nxf.WEzLzfPWQm/TYJH7L_5_feature-counts.tsv.summary`

Copy Prompt


Summarize table

| Overrepresented sequence | Reports | Occurrences | % of all reads |
| --- | --- | --- | --- |
| CCCTAATACCTGCCACCCCACTCTTAATCAGTGGTGGAAGAACGGTCTCA | 1 | 29495 | 0.1546% |
| GTACATTCCACAAGCATTGCCTTCTTATTTTACTTCTTTTAGCTGTTTAA | 1 | 28651 | 0.1502% |
| GGCCCAAGGTGTCCTGCAGGCTGTAATGCAGTTTAATCAGAGTGCCATTT | 1 | 28615 | 0.1500% |
| CTCTTAATCAGTGGTGGAAGAACGGTCTCAGAACTGTTTGTTTCAATTGG | 1 | 21055 | 0.1104% |

Close
