## Supplementary Data 1 for "Genetically inducible coordinators of cytokine signaling pathways for interrogating T cell motility": TYJH7L_6_pYRS0255_UNT2.html

Loading report..

Report
generated on 2026-08-05, 21:59 UTC
based on data in:

- `/tmp/nxf.tNLfZAC4BN/rustqc`
- `/tmp/nxf.tNLfZAC4BN/TYJH7L_6_filtered_fastqc.zip`
- `/tmp/nxf.tNLfZAC4BN/TYJH7L_6_QC-short-reads.json`
- `/tmp/nxf.tNLfZAC4BN/TYJH7L_6_Log.final.out`
- `/tmp/nxf.tNLfZAC4BN/TYJH7L_6_feature-counts.tsv.summary`

Copy Prompt


Summarize table

| Overrepresented sequence | Reports | Occurrences | % of all reads |
| --- | --- | --- | --- |
| CCCTAATACCTGCCACCCCACTCTTAATCAGTGGTGGAAGAACGGTCTCA | 1 | 31798 | 0.1545% |
| GTACATTCCACAAGCATTGCCTTCTTATTTTACTTCTTTTAGCTGTTTAA | 1 | 29411 | 0.1429% |
| GGCCCAAGGTGTCCTGCAGGCTGTAATGCAGTTTAATCAGAGTGCCATTT | 1 | 28905 | 0.1404% |
| CTCTTAATCAGTGGTGGAAGAACGGTCTCAGAACTGTTTGTTTCAATTGG | 1 | 22896 | 0.1112% |

Close
