## Supplementary Data 1 for "Genetically inducible coordinators of cytokine signaling pathways for interrogating T cell motility": TYJH7L_7_pYRS0255_500DOX2.html

Loading report..

Report
generated on 2026-08-05, 21:55 UTC
based on data in:

- `/tmp/nxf.hvSzhdSdil/rustqc`
- `/tmp/nxf.hvSzhdSdil/TYJH7L_7_filtered_fastqc.zip`
- `/tmp/nxf.hvSzhdSdil/TYJH7L_7_QC-short-reads.json`
- `/tmp/nxf.hvSzhdSdil/TYJH7L_7_Log.final.out`
- `/tmp/nxf.hvSzhdSdil/TYJH7L_7_feature-counts.tsv.summary`

Copy Prompt


Summarize table

| Overrepresented sequence | Reports | Occurrences | % of all reads |
| --- | --- | --- | --- |
| GTACATTCCACAAGCATTGCCTTCTTATTTTACTTCTTTTAGCTGTTTAA | 1 | 26488 | 0.1330% |
| GGCCCAAGGTGTCCTGCAGGCTGTAATGCAGTTTAATCAGAGTGCCATTT | 1 | 25535 | 0.1282% |
| CCCTAATACCTGCCACCCCACTCTTAATCAGTGGTGGAAGAACGGTCTCA | 1 | 25398 | 0.1275% |

Close
