## Supplementary Data 1 for "Genetically inducible coordinators of cytokine signaling pathways for interrogating T cell motility": TYJH7L_8_pYRS0257_UNT2.html

Loading report..

Report
generated on 2026-08-05, 22:22 UTC
based on data in:

- `/tmp/nxf.lyfz5LzC0N/rustqc`
- `/tmp/nxf.lyfz5LzC0N/TYJH7L_8_filtered_fastqc.zip`
- `/tmp/nxf.lyfz5LzC0N/TYJH7L_8_QC-short-reads.json`
- `/tmp/nxf.lyfz5LzC0N/TYJH7L_8_Log.final.out`
- `/tmp/nxf.lyfz5LzC0N/TYJH7L_8_feature-counts.tsv.summary`

Copy Prompt


Summarize table

| Overrepresented sequence | Reports | Occurrences | % of all reads |
| --- | --- | --- | --- |
| CCCTAATACCTGCCACCCCACTCTTAATCAGTGGTGGAAGAACGGTCTCA | 1 | 29790 | 0.1536% |
| GGCCCAAGGTGTCCTGCAGGCTGTAATGCAGTTTAATCAGAGTGCCATTT | 1 | 29482 | 0.1520% |
| GTACATTCCACAAGCATTGCCTTCTTATTTTACTTCTTTTAGCTGTTTAA | 1 | 29193 | 0.1505% |
| CTCTTAATCAGTGGTGGAAGAACGGTCTCAGAACTGTTTGTTTCAATTGG | 1 | 20653 | 0.1065% |

Close
