## Supplementary Data 1 for "Genetically inducible coordinators of cytokine signaling pathways for interrogating T cell motility": TYJH7L_9_pYRS0257_500DOX2.html

Loading report..

Report
generated on 2026-08-05, 21:59 UTC
based on data in:

- `/tmp/nxf.n64k39zlOL/rustqc`
- `/tmp/nxf.n64k39zlOL/TYJH7L_9_filtered_fastqc.zip`
- `/tmp/nxf.n64k39zlOL/TYJH7L_9_QC-short-reads.json`
- `/tmp/nxf.n64k39zlOL/TYJH7L_9_Log.final.out`
- `/tmp/nxf.n64k39zlOL/TYJH7L_9_feature-counts.tsv.summary`

Copy Prompt


Summarize table

| Overrepresented sequence | Reports | Occurrences | % of all reads |
| --- | --- | --- | --- |
| GTACATTCCACAAGCATTGCCTTCTTATTTTACTTCTTTTAGCTGTTTAA | 1 | 30110 | 0.1556% |
| GGCCCAAGGTGTCCTGCAGGCTGTAATGCAGTTTAATCAGAGTGCCATTT | 1 | 28364 | 0.1466% |
| CCCTAATACCTGCCACCCCACTCTTAATCAGTGGTGGAAGAACGGTCTCA | 1 | 25293 | 0.1307% |

Close
