## Supplementary Data 1 for "Genetically inducible coordinators of cytokine signaling pathways for interrogating T cell motility": TYJH7L_10_Naive_UNT2.html

Loading report..

Report
generated on 2026-08-05, 21:59 UTC
based on data in:

- `/tmp/nxf.5DWE7we3Wp/rustqc`
- `/tmp/nxf.5DWE7we3Wp/TYJH7L_10_filtered_fastqc.zip`
- `/tmp/nxf.5DWE7we3Wp/TYJH7L_10_QC-short-reads.json`
- `/tmp/nxf.5DWE7we3Wp/TYJH7L_10_Log.final.out`
- `/tmp/nxf.5DWE7we3Wp/TYJH7L_10_feature-counts.tsv.summary`

Copy Prompt


Summarize table

| Overrepresented sequence | Reports | Occurrences | % of all reads |
| --- | --- | --- | --- |
| CCCTAATACCTGCCACCCCACTCTTAATCAGTGGTGGAAGAACGGTCTCA | 1 | 27529 | 0.1385% |
| GTACATTCCACAAGCATTGCCTTCTTATTTTACTTCTTTTAGCTGTTTAA | 1 | 26529 | 0.1335% |
| GGCCCAAGGTGTCCTGCAGGCTGTAATGCAGTTTAATCAGAGTGCCATTT | 1 | 26021 | 0.1309% |
| CTCTTAATCAGTGGTGGAAGAACGGTCTCAGAACTGTTTGTTTCAATTGG | 1 | 19922 | 0.1002% |

Close
