## Supplementary Data 1 for "Genetically inducible coordinators of cytokine signaling pathways for interrogating T cell motility": TYJH7L_11_Naive_500DOX2.html

Loading report..

Report
generated on 2026-08-05, 21:57 UTC
based on data in:

- `/tmp/nxf.nEiLrgaA9e/rustqc`
- `/tmp/nxf.nEiLrgaA9e/TYJH7L_11_filtered_fastqc.zip`
- `/tmp/nxf.nEiLrgaA9e/TYJH7L_11_QC-short-reads.json`
- `/tmp/nxf.nEiLrgaA9e/TYJH7L_11_Log.final.out`
- `/tmp/nxf.nEiLrgaA9e/TYJH7L_11_feature-counts.tsv.summary`

Copy Prompt


Summarize table

| Overrepresented sequence | Reports | Occurrences | % of all reads |
| --- | --- | --- | --- |
| CCCTAATACCTGCCACCCCACTCTTAATCAGTGGTGGAAGAACGGTCTCA | 1 | 27103 | 0.1436% |
| GTACATTCCACAAGCATTGCCTTCTTATTTTACTTCTTTTAGCTGTTTAA | 1 | 25946 | 0.1375% |
| GGCCCAAGGTGTCCTGCAGGCTGTAATGCAGTTTAATCAGAGTGCCATTT | 1 | 25476 | 0.1350% |
| CTCTTAATCAGTGGTGGAAGAACGGTCTCAGAACTGTTTGTTTCAATTGG | 1 | 18899 | 0.1001% |

Close
