## Supplementary Data 1 for "Genetically inducible coordinators of cytokine signaling pathways for interrogating T cell motility": TYJH7L_12_Naive_PMA2.html

Loading report..

Report
generated on 2026-08-05, 21:53 UTC
based on data in:

- `/tmp/nxf.su6QmDCOmQ/rustqc`
- `/tmp/nxf.su6QmDCOmQ/TYJH7L_12_filtered_fastqc.zip`
- `/tmp/nxf.su6QmDCOmQ/TYJH7L_12_QC-short-reads.json`
- `/tmp/nxf.su6QmDCOmQ/TYJH7L_12_Log.final.out`
- `/tmp/nxf.su6QmDCOmQ/TYJH7L_12_feature-counts.tsv.summary`

Copy Prompt


Summarize table

| Overrepresented sequence | Reports | Occurrences | % of all reads |
| --- | --- | --- | --- |
| CCCTAATACCTGCCACCCCACTCTTAATCAGTGGTGGAAGAACGGTCTCA | 1 | 39469 | 0.2046% |
| GTACATTCCACAAGCATTGCCTTCTTATTTTACTTCTTTTAGCTGTTTAA | 1 | 36821 | 0.1909% |
| GGCCCAAGGTGTCCTGCAGGCTGTAATGCAGTTTAATCAGAGTGCCATTT | 1 | 35648 | 0.1848% |
| CTCTTAATCAGTGGTGGAAGAACGGTCTCAGAACTGTTTGTTTCAATTGG | 1 | 27424 | 0.1422% |
| CTGTAATGCAGTTTAATCAGAGTGCCATTTTTTTTTTTGTTCAAATGATT | 1 | 22037 | 0.1142% |
| AATATGCACTGTACATTCCACAAGCATTGCCTTCTTATTTTACTTCTTTT | 1 | 19430 | 0.1007% |
| GTTTAAATGACTGTGCTGCCCCTTTCACATCAAAGAACTACTGACAACGA | 1 | 19308 | 0.1001% |

Close
