## Supplementary Data 1 for "Genetically inducible coordinators of cytokine signaling pathways for interrogating T cell motility": README_RNAseq.txt

================================================================================
Supplemental Data Genetically inducible coordinators of cytokine signaling pathways for probing T cell motility phenomena — RNA-seq
================================================================================

Bulk RNA-seq of engineered primary human T cells, sequenced and analyzed by
Plasmidsaurus. Two sequencing runs (TYJH7L and 8WTJWN) were pooled; all
comparisons labelled "bothruns" combine samples from both.

Contents:
  Pooled_DGE_compiled.xlsx            differential expression, compiled
  Pooled_GSEA_Analysis_Results.xlsx   gene set enrichment, compiled
  20260731_RNA-Seq-Plasmidsaurus-Files_TYJH7L/   raw delivery, run 1
  20260727_RNA-Seq-Plasmidsaurus-Files_8WTJWN/   raw delivery, run 2

The two workbooks are the Plasmidsaurus per-comparison CSV outputs collected
into one file each; nothing was recomputed.

Constructs: pYRS0239 = Ctrl-LumiScarlet, pYRS0240 = CAMP-7, pYRS0255 =
CAMP-24, pYRS0257 = CAMP-26. Naive = unmodified T cells.


--------------------------------------------------------------------------------
DGE WORKBOOK
--------------------------------------------------------------------------------

"Contents" sheet lists every comparison with its construct ID and source
filename. Seven comparison tabs, named sample_baselinevstreatment_run:

  Naive              untreated vs 500 ng/mL dox;  untreated vs PMA
  Ctrl-LumiScarlet   untreated vs 500 ng/mL dox;  untreated vs PMA
  CAMP-7             untreated vs 500 ng/mL dox
  CAMP-24            untreated vs 500 ng/mL dox
  CAMP-26            untreated vs 500 ng/mL dox


  logFC is treatment vs baseline (positive = higher in the treated group).
  Rows with FDR < 0.05 are shaded light green.


--------------------------------------------------------------------------------
GSEA WORKBOOK
--------------------------------------------------------------------------------

21 sheets: the same seven comparisons above, each run against three gene set
libraries — GO Biological Process 2025, MSigDB Hallmark 2025, and Reactome
2024. Sheet names follow sample_untvstreatment_library.


--------------------------------------------------------------------------------
PLASMIDSAURUS DELIVERY FOLDERS
--------------------------------------------------------------------------------

Unmodified as downloaded. Each contains:

  RNA-seq_results.xlsx               all per-comparison results in one book
  <run>-expression-matrix.csv/.tsv   normalised counts, all genes x samples
  <run>-methods.txt                  library prep, sequencing, and analysis
                                     methods as provided by Plasmidsaurus
  <run>-multiqc-report.html          aggregate QC
  <run>-per-sample-multiqc-report/   per-sample QC
  <run>-mapping-stats/               and -mapping-stats-reads.csv
  <run>-gene-biotype-5plus-reads     biotype breakdown (.html and summary .csv)
  <run>-correlation-heatmap.png      sample-sample correlation
  <run>-pca-scatter.html             PCA


--------------------------------------------------------------------------------
NOTES
--------------------------------------------------------------------------------

- The dox comparison is written "untvs500dox" in the DGE workbook and
  "untvsdox" in the GSEA workbook. Same comparison, 500 ng/mL dox.
- Several GSEA sheet names are truncated (e.g. "Ctrl-LumiScarlet_untvsdox_GOBP2")
  because Excel caps sheet names at 31 characters.
