## Supplementary figures and images for "Genetically inducible coordinators of cytokine signaling pathways for interrogating T cell motility"

### 8WTJWN-correlation-heatmap.png

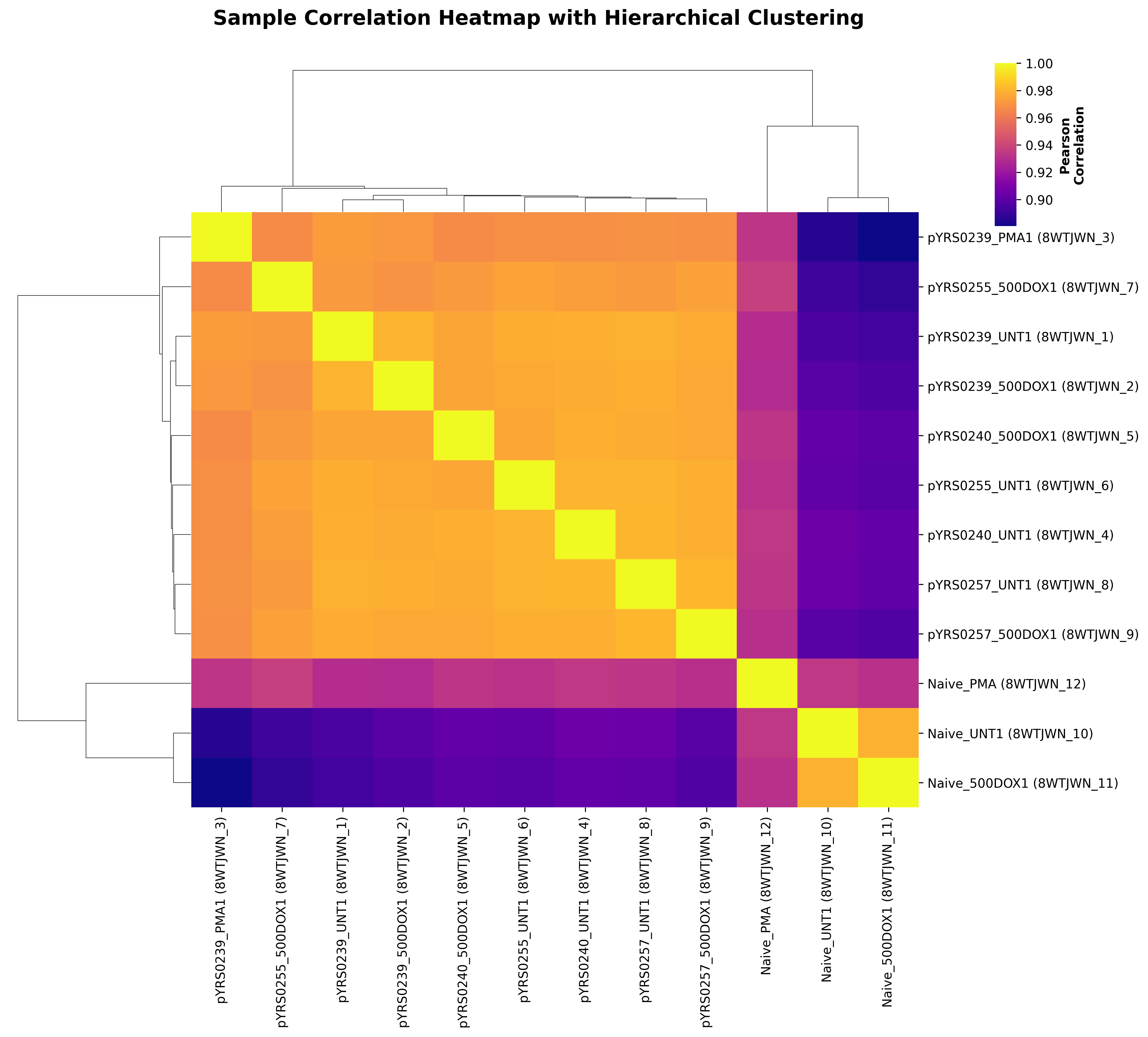

### TYJH7L-correlation-heatmap.png

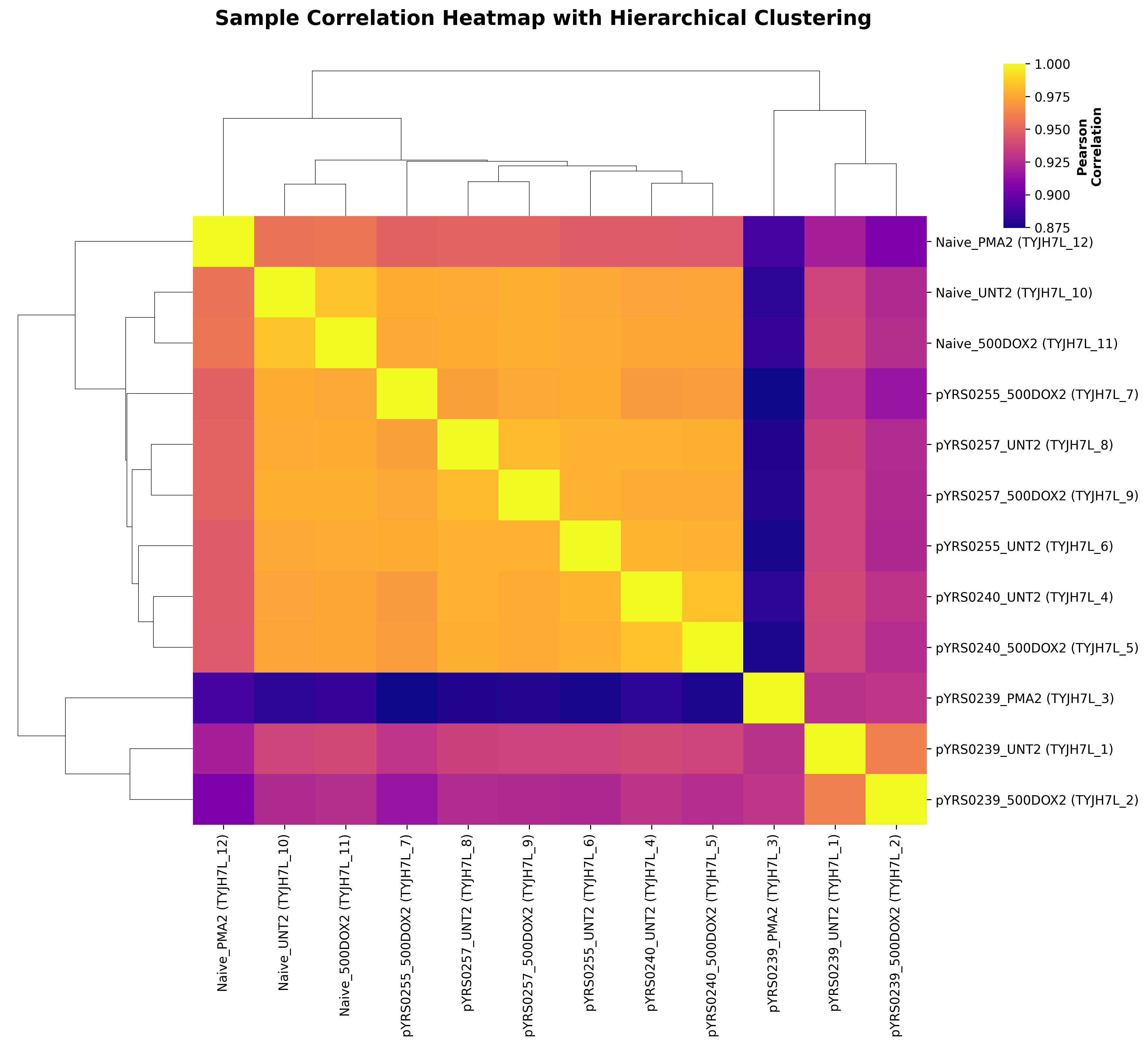
