## Supplementary Data 2 for "Genetically inducible coordinators of cytokine signaling pathways for interrogating T cell motility": README_Plasmids.txt

================================================================================
Genetically inducible coordinators of cytokine signaling pathways for probing T cell motility phenomena — Plasmids used in this study
Plasmids_for_Experiments_Figure_Annotated.xlsx
================================================================================

Every plasmid used in the study, grouped by function across five sheets, with
the figure panels each one supports.

  Sheet                     Entries   Contents
  -----------------------   -------   ------------------------------------
  TUPVs                        32     Transcriptional unit plasmids —
                                      the CAMP receptor constructs and the
                                      modules used to build the transposon
                                      vectors
  Transposon Vectors            6      PiggyBac transposon plasmids used to
                                      engineer primary human T cells
  Transposase Vectors           1      hyperactive PiggyBac transposase
  Fluorescent Proteins          2      dsRedExpress2 and mNeonGreen
                                      transfection/expression controls
  Empty Vector Filler DNA       1      empty pcDNA backbone used to equalize
                                      total DNA mass across transfections

All five sheets share the same six columns:

  Plasmid Number and Name   full construct name
  CAMP identifier           the CAMP-n name used in the figures; "—" where
                            the plasmid is not a CAMP
  Reference number          the pYRS/pJW/pPD/pHIE number used in filenames
                            throughout the deposited raw data
  Purpose                   what the plasmid is for
  Relevant Figure Panels    where it appears
  Notable Components        promoter, domains, and markers

Use the "Reference number" column to connect this table to the raw data: it is
the identifier that appears in the flow cytometry filenames (e.g. CAMP_255 =
pYRS0255), the movie folder names, and the RNA-seq comparison names.


NOTES

- CAMP identifiers run CAMP-1 through CAMP-26, plus Ctrl-LumiScarlet. Not all
  numbers appear in the main figures; several were built and characterized but
  are reported only in supplementary panels or not at all.
- Four constructs appear twice in TUPVs — once as the standalone receptor and
  once as a dox-inducible module (marked "(module)" in the CAMP identifier
  column); this is because the module version was used to build the corresponding
  transposon vector.
