## Supplementary Data 3 for "Genetically inducible coordinators of cytokine signaling pathways for interrogating T cell motility": README_SupplementalData3.txt

================================================================================
Supplemental Data 3 — FigureData_SupplementalData3.xlsx
Source data for all main and supplementary figure panels for Genetically inducible coordinators of cytokine signaling pathways for probing T cell motility phenomena
================================================================================

One sheet per figure panel, named for the panel it generates (Fig1c, Fig4b,
SuppFig8a, ...). The "Legend" sheet lists all 12 experiments with a one-line
description and the panels each one supports; start there.

Sheets fall into three layouts.


1. REPORTER ASSAY PANELS
   Fig1c, Fig1d, Fig2c, Fig2d, Fig3a-3d, SuppFig2a-b, SuppFig4a-b, SuppFig7a-d

   Sample | Mean | SEM | N

   One row per construct. HEK-Blue panels report A620 SEAP absorbance; HEK-NEE panels report mNeonGreen MFI in FITC channel. N = 3 technical replicates throughout.


2. T CELL MOTILITY PANELS
   Fig4b, Fig4c, Fig6b, SuppFig8a-d, SuppFig9a-c, SuppFig10a-c

   Per-well summary statistics from a two-component mixture model fitted to
   single-cell motility. Fig4b/4c/6b and SuppFig9/10 are per-well; SuppFig8 is
   aggregated per condition.

   Column glossary:
     n_cells, n_used, n_raw   cells contributing to the fit; n_raw before
                              track filtering, frac_dropped = fraction removed
     med_y                    median of per-cell log10 values
     med_um2                  median squared displacement of raw µm² values
     mu_lo, mu_hi             mean of the low- and high-motility components
                              (same log10 scale as med_y)
     w_hi                     mixing weight of the high-motility component
     n_lo_eff, n_hi_eff       effective N per component
     lag_sec                  MSD lag time, seconds (600 s = 10 min)
     lo_min/lo_max,           range across replicates (SuppFig8 only)
       hi_min/hi_max
     converged, weak_low,     fit diagnostics; rows flagged weak or
       weak_high, weak_sep,   non-converged should be interpreted with care
       sigma_unstable
     well                     matches the movie filenames in https://doi.org/10.5281/zenodo.22069801


3. FLOW CYTOMETRY GATING
   Fig5c

   FlowJo gating table, one row per .fcs sample: event counts and percent of
   parent at each gate (T cells, singlets, live, transposon-engineered).
   Row labels match the .fcs filenames in https://doi.org/10.5281/zenodo.22069801.


--------------------------------------------------------------------------------
NOTES
--------------------------------------------------------------------------------

- Acute PMA doses are 2, 10, and 50 ng/mL. Chronic PMA doses (0.08, 0.4,
  2 ng/mL) carry no unit in the well strings.
- Constructs: pYRS0239 = Ctrl-LumiScarlet, pYRS0240 = CAMP-7, pYRS0255 =
  CAMP-24, pYRS0257 = CAMP-26. Full plasmid key in Supplementary Table 1.
- Replicates are n = 3 for the reporter assays and the dox motility experiment, n = 2
  for the density and PMA motility experiments.
